# Polystyrene nanoplastics promote neurodegeneration by catalyzing TDP43 hyperphosphorylation

**DOI:** 10.1101/2024.11.11.622894

**Authors:** Winanto, Li-Yi Tan, Wai Hon Chooi, Cheryl Yi-Pin Lee, Wan Yun Ho, Yong Shan Lim, Boon Seng Soh, Emma Sanford, Chong-Lun Tan, Yih-Cherng Liou, Cathy Chia-Yu Huang, Shuo-Chien Ling, Shi-Yan Ng

**Affiliations:** Institute of Molecular and Cell Biology (Cell and Biological Therapies Division), A*STAR Research Entities, Singapore 138673 Institute of Molecular and Cell Biology (IMCB), Agency for Science, Technology and Research (A*STAR), 61 Biopolis Drive, Proteos, Singapore 138673, Republic of Singapore; National University of Singapore, Faculty of Science (Department of Biological Science), Singapore 117543; National University of Singapore, Yong Loo Lin School of Medicine (Department of Physiology), Singapore 117456; Programs in Neuroscience and Behavioral Disorders, Duke-NUS Medical School, Singapore, SG; Department of Life Sciences, National Central University, Taoyuan, Taiwan; Healthy Longevity Translational Research Programme, Yong Loo Lin School of Medicine, National University of Singapore, Singapore, Singapore, SG; National Neuroscience Institute, Singapore 308433

## Abstract

The ubiquity of polystyrene nanoplastics (PS-NPs) in our environment raises substantial concerns about their potential impact on human health. Recent studies have shown that PS-NPs cross the blood-brain-barrier and accumulate in the central nervous system (CNS), raising the concerns on the causal role of PS-NP exposure and neurodegenerative diseases. In this study, we utilized human-induced pluripotent stem cell-derived neurons to examine the effects of PS-NPs on neuronal function and health. Our results revealed that PS-NPs penetrate neuronal cells in a size-dependent manner, and are bound by various cellular proteins, including TDP43, a key protein implicated in amyotrophic lateral sclerosis (ALS). Interestingly, CK1 and GSK3β kinases that are known to phosphorylate TDP-43 were found to be associated with PS-NPs. This observation suggests that PS-NPs may play a role in facilitating conditions that lead to TDP43 phosphorylation. We further demonstrate that exposure to healthy motor neurons with PS-NPs resulted in ALS-like phenotypes, characterized by hyperphosphorylated TDP-43, disrupted neuronal morphology, impaired mitochondrial respiration, and accelerated motor neuron death. These findings suggest that PS-NPs contribute to the pathogenesis of neurodegenerative diseases such as ALS and highlight the urgent need for strategies to limit human exposure to nanoplastics.

## INTRODUCTION

Huge amounts of plastics are being used in the developed world and it is estimated that individuals could ingest approximately 5 grams of plastic weekly from drinking water as well as seafood consumption [1–4]. Plastics do not undergo biodegradation, but instead break down into smaller pieces such as microplastics (between 0.1 to 5 mm in size) [5] and nanoplastics (less than 100 nm) [6], which eventually get transferred up the food chain. In aquatic organisms, nanoplastics exposure and bioaccumulation have been shown to result in endocrine disruption [7], intestinal damage [8], reduced reproduction [9–11] and locomotive defects [12, 13]. There is accumulating evidence demonstrating the nanoplastics cross the blood-brain barrier in mammals and disrupt neuronal function and behavior [14, 15]. In mice, nanoplastics can be detected in the brain two hours after oral ingestion, through an uptake mechanism that involves cholesterol [19]. Mice chronically exposed to high doses of nanoplastics also show progressive loss of blood-brain barrier integrity, neuronal dysfunction, elevated neuroinflammatory response and Parkinson’s disease-like neurodegeneration [16].

Given that polystyrene is a major environmental pollutant that is most commonly used in food packaging and can leech into the environment and enter food that we ingest [17], we questioned if polystyrene nanoplastics and microplastics impact neuronal health and function, which contributes to neurodegeneration. Polystyrene nanoplastics (PS-NPs) have been shown to cross the blood-brain barrier [15] and are taken up by cells in the central nervous system. However, it is still unclear if and how PS-NPs contribute to neurodegeneration in humans. In this study, we combine proteomics tools and human induced pluripotent stem cell-based modeling and demonstrated that PS-NP exposure leads to neurodegenerative phenotypes that resemble Amyotrophic Lateral Sclerosis (ALS). We found that PS-NPs are readily internalized by neurons and interact with Tar-DNA binding protein 43-kDa (TDP43) and its associated kinases. These interactions may potentially bring these molecules into closer proximity, which could facilitate conditions favorable for TDP43 hyperphosphorylation. In human motor neurons, PS-NPs also result in defective mitochondrial respiration metabolism, disease-associated neuronal morphologies, and accelerated motor neuron death. Our data therefore adds onto the growing body of evidence strongly suggesting that PS-NPs contribute to the pathogenesis of neurodegenerative diseases, and our mechanistic study suggests that this is a result of the catalytic scaffold function of PS-NPs.

## EXPERIMENTAL METHODS

### Maintenance and differentiation of ReN-VM neural stem cells (NSCs)

The human neural stem cell line, ReN-VM was maintained on Matrigel-coated (Corning, 356234) plates with N2B27 media comprising of 1:1 mix of DMEM/F12 (Gibco™, 11330032) and Neuro Medium (Miltenyi Biotec, 130-093-570), and supplemented with 1% GlutaMAX supplement (Gibco, 35050079), 1% MEM Non-Essential Amino Acids Solution (Gibco, 11140050), 1% N2 supplement and 2% NeuroBrew-21 (Miltenyi Biotec, 130-093-566), as well as 10 ng/ml of human bFGF (Miltenyi Biotec 130-093-838) and 20 ng/ml of human EGF (Miltenyi Biotec 130-093-825). To differentiate ReN-VM into neurons, ReN-VM were grown in N2B27 media without bFGF and EGF for 7 days.

To generate ReN-VM spheroids, ReN-VM NSCs were encapsulated in 30X Matrigel (Corning, 356234) at 40,000 cells per 5 μl Matrigel droplet. These droplets were incubated at 37°C for 2-3hours before the addition of sufficient N2B27 media with bFGF and EGF to cover every droplet. After four days, droplets were transferred to a low-attachment 10-cm plate and switched to the N2B27 media without bFGF and EGF, and placed on an orbital shaker for the next 7 days.

### Culture and maintenance of human pluripotent stem cells (hPSCs)

The human induced pluripotent stem cell (hPSC) line BJ-iPS and embryonic stem cell line H7 (WiCell) were maintained on feeder-free Matrigel coated dishes in StemMACS™ iPS-Brew XF, human (Miltenyi Biotec, 130-104-368) at 37 °C in a humidified incubator with 5% CO_2_. Periodic passaging of iPSCs were conducted weekly using ReLeSR passaging reagent (STEMCELL Technologies, 100-0484) at 1:6 split ratio every week.

### Generation of spinal motor neurons

Pluripotent stem cells were differentiated into spinal motor neurons (MNs) fate following a previously established protocol [39, 50]. In brief, neural induction of hPSCs was achieved with 4.25μM of CHIR99021(Miltenyi Biotec, 130-103-926) and 0.5 μM of LDN193189 (Miltenyi Biotec, 130-106-540) from days 1 to 7. On day 3 of differentiation, 1 μM of retinoic acid (Sigma-Aldrich, R2625) was added to caudalize the culture. At day 10, cultures were ventralized by adding 1 μM purmorphamine (Miltenyi Biotec, 130-104-465). 10 ng/ml brain-derived neurotrophic factor (BDNF) (Miltenyi Biotec, 130-103-435) and 10 ng/ml of glial-derived neurotrophic factor (Miltenyi Biotec, 130-129-544) were added from day 17 to 28 to promote neuronal maturation. Additionally, to eliminate progenitor cells in culture, a single dose of cytosine β-D-arabinofuranoside (Ara-C, C1768, Sigma Aldrich) at 1 μM was added at day 21 of differentiation.

### Exposure of cultured neurons to polystyrene nanoplastics (PS-NPs)

Fluorescent polystyrene micro- and nano-plastic particles with an emission wavelength of 488-nm were obtained from Phosphorex Inc and Polysciences Asia Pacific Inc. Plastics of three different sizes were used in this study - 50nm (Phosphorex, 2101) (Polyscience Inc, 17149-10), 500nm (Phosphorex, 2103A), and 5000nm (Phosphorex, 2106B). All Phosphorex PS-NPs are supplied as 1% plastic suspensions (10 mg/ml) with density of 1.06 g/cm³. In this study, PS-NPs were exposed to cultured neurons at 2 concentrations: 0.001% and 0.01%, which is equivalent to 1:1000 and 1:100 dilution respectively.

### Flow cytometry

Since PS-NPs emit at 488-nm, flow cytometry was used to determine the percentage of cells with detectable green fluorescence, which is indicative of cellular uptake of PS-NPs. After treatment with respective plastic particles, neurons were dissociated into single cells using Accutase and washed twice with PBS. Cells were resuspended in PBS containing 5% BSA and filtered through a 45-μm cell filter to remove any cell clumps. Flow cytometry was performed using BD FACSAria™ III Cell Sorter (BD Bioscience) and analyzed using FlowJo.

### Annexin V apoptosis assay

Apoptosis was measured using the Annexin V - PE detection Kit I (BD Pharmingen™, 559763) following the manufacturer’s instructions. In brief, following 24 hours PS-NPs treatment, ReN-VM neurons were dissociated into single cells, washed twice with ice-cold PBS, and resuspended in 1x binding buffer at 1 x 10^6^ cells/ ml. 100 µl of this cell suspension were incubated with 5 µl of PE Annexin V and 5 µl propidium iodide at 25°C in the dark. Cells were analyzed immediately with BD FACSymphony™ A3 Cell Analyzer (BD biosciences). Each treatment was performed in triplicates with PBS as a control. Flow cytometry was used to quantify the number of dying and apoptotic cells. To visualize cells undergoing cell death, another independent experiment was performed with ReN-VM neurons seeded on 96 well-plates at 20,000 cells per well. Fluorescent images were captured through a high content microscope Operetta (Perkin Elmer) using the 20x objective. Cell counts and intensity measurements were obtained using the Columbus (Perkin Elmer) software.

### Immunostaining and image acquisition

ReN-VM cells were seeded at 25,000 cells per well of a 96-well optical plate for immunostaining while iPSC-derived neurons were seeded at 75,000 cells per well of a 96-well plate. Cells were fixed in 4% paraformaldehyde for 20mins, permeabilized using 0.1% Triton® X-100 (Promega Corporation. H5142) for 20 minutes and subsequently blocked for 2 hours with immunostaining buffer that contains PBS with 5% of bovine serum albumin (BSA; Hyclone SH30574.02) and 10% of fetal bovine serum (FBS; Gibco 10437028). After blocking, neurons were probed with primary antibodies overnight at 4 °C. The list of primary antibodies used and their respective dilutions are listed in **Table 1**. The following day, primary antibodies were aspirated and secondary antibodies were diluted in the blocking buffer at a concentration of 1:1500 and incubated with cells for 1 hour in room temperature. Cells were then washed thrice with PBS prior to imaging. DAPI (0.1ug/ml) was used to visualize cellular nuclei. Images were obtained using the high content microscope Phenix (Perkin Elmer) using the 20x objective. Cell counts and intensity measurements were obtained using the Columbus (Perkin Elmer) software.

**Table 1:** list of antibodies for immunostaining.

| S/N | Antibody | Dilution | Source |
| --- | --- | --- | --- |
| 1 | Beta-Tubulin III antibody, TUJ1 | 1:1000 | Biologend, 801202 |
| 2 | Islet-1 (ISL1) | 1:1500 | Abcam, AB203406 |
| 3 | SMI32 | 1:2000 | Calbiochem, NE-1023 |
| 4 | TDP43 (C-Terminal) | 1:1000 | Proteintech, 12892-1-AP |
| 5 | Phospho-TDP43 (Ser409/410) | 1:1000 | Proteintech, 22309-1-AP |

**Table 2:** list of antibodies for western blot.

| S/N | Antibody | Dilution | Source |
| --- | --- | --- | --- |
| 1 | TDP43 (C-Terminal) | 1:500 | Proteintech, 12892-1-AP |
| 2 | Phospho-TDP43 (Ser409/410) | 1:500 | Proteintech, 22309-1-AP |
| 3 | Tau | 1:1000 | Tako |
| 4 | Tau phospho Serine 199/202 | 1:1000 | Merck, AB9674 |
| 5 | ApoE (pan) | 1:1000 | Cell Signal, #13366 |
| 6 | GAPDH | 1:1000 | Proteintech, 10494-1-AP |
| 7 | GSK-3β | 1:1000 | BD Biosciences, 610201 |
| 8 | Casein Kinase 1 alpha | 1:1000 | Proteintech, 55192-1-AP |
| 9 | Casein Kinase 1 delta | 1:1000 | Proteintech, 14388-1-AP |

ReNVM spheroids were cryosectioned at 10um per section. Samples were permeabilised using 0.5% TritonX-100 for 20 mins, and blocked with our immunostaining buffer as described above. After blocking, samples were probed with primary antibodies overnight at 4°C. Primary antibody solution washed off with PBS. Secondary antibodies (1:1500) were diluted in the blocking buffer and incubated with cells for 1 hour in room temperature and washed off after.

### Neurite outgrowth assay

ReN-VM neurons were seeded on 96 well plate at 20,000 per well. After PS-NP treatment, plates were fixed 4% paraformaldehyde, permeabilized with 0.1% tritonX and stained with anti-Beta-Tubulin III antibody, TUJ1 to label the neuronal extensions. Neurite lengths were determined using NeuriteTracer, an automated neurite outgrowth plug-in from ImageJ. At least 50 neurites from different neurons were measured and assessed for statistical significance.

### Measurement of mitochondrial ROS

Mitochondrial reactive oxygen species (ROS) was measured using MitoSOX (Invitrogen, M36008), following the manufacturer’s instructions. Briefly, cells were incubated with MitoSOX reagent for 30 minutes at 37°C in 5% CO_2_ incubator. After washing with PBS, cellular imaging was performed using the high content microscope Phenix (Perkin Elmer) using the 20x objective. Cell counts and intensity measurements were obtained using the Columbus (Perkin Elmer) software.

### EdU proliferation assay

Proliferation of ReN-VM neurons were measured using Click-iT EdU Cell Proliferation Kit for Imaging, Alexa Fluor-647 dye (Invitrogen, C10340). After 3 days PS-NP treatment, ReN-VM neurons were pulsed with 10 μM EdU for 3 hours. Cells were washed once with PBS followed by 15 min incubation with Click-iT fixative in the dark at room temperature. Cells were then fixed and counterstained with DAPI, and using the high content microscope Phenix (Perkin Elmer) with the 20x objective. The percentage of EdU-positive cells were then analyzed using Columbus (Perkin Elmer).

### Nanoparticle pull down assay

Human motor neuron lysates were prepared by resuspending 30 million cells in 1 ml of PBS with protease inhibitors added (Roche, 4693132001). Cell suspension was put through freeze thawing cycle thrice to lyse the cells. Thereafter, cell debris were removed by centrifugation at 13,200 rpm for 15 minutes. The supernatant was then transferred to a new 1.5-ml tube and incubated with either 50 nm PS-NPs, 50 nm Silica gold (Sigma Aldrich, 797936) or 50 nm gold (Sigma Aldrich, 753645) for 30 minutes. Thereafter, nanoparticles were pelleted at 13,200 RPM for 20 minutes at 4 °C. Supernatant consisting of unbounded protein was removed. The respective pellets were subjected to washing with non-denaturing buffer consisting of 20mM Tris HCL at pH8, 137 mM NaCl, 2mM EDTA and 1% Nonidet P-40 (NP-40) to remove any non-specific binding. The washed pellet consisting nanoparticle-bound proteins were then analyzed by LC-MS/MS as well as SDS-PAGE and Western blot.

### Western blot

Protein was harvested using RIPA buffer (Life Technologies Pte Ltd, 89900) and protein concentration quantified using the Pierce™ BCA Protein Assay Kits (Thermo Scientific™, 23225) following the manufacturer’s protocol. Lysates were resolved in 8-12% SDS-PAGE gels depending on the molecular weight of protein of interest in the Tris-Glycine-SDS buffer. After resolving, proteins were transferred onto a PVDF membrane and blocked in TBS-T with 5% BSA for an hour. Primary antibodies were diluted in the blocking buffer with membrane overnight at 4 °C. After which, membranes were washed in TBS-T three times for 15 minutes each. The respective secondary antibodies were diluted in the blocking buffer and incubated with membrane for an hour in room temperature. Membranes were then washed three times. Images were obtained using the ChemiDoc after ECL exposure (Bio-Rad Laboratories, 1705060).

### Motor neuron survival assay

Motor neuron survival was calculated based on the numbers of ISL1+ motor neurons in culture from day 28 to day 35 [39]. First, 1 μM AraC was added to motor neuron cultures at day 21 to eliminate progenitor cells. Cells were then reseeded onto 96-well plates at 60,000 cells per well on day 27. Plates with cells treated with or without PS-NPs were then fixed on days 28, 31 and 35. Immunostaining was performed using specific antibodies against ISL1 and SMI32. Dilutions and details of primary antibodies are described in Table 1. The motor neuron survival index was calculated with biological triplicates and a minimum of 5 technical replicates.

### Analysis of mitochondrial respiration with metabolic flux assays

To measure mitochondrial respiration in motor neurons, day 27 cultures were first enriched for motor neurons by using antibodies against PSA-NCAM and L1CAM (both from Miltenyi Biotec) [39]. Thereafter, enriched neurons were seeded onto Seahorse 96-well culture plates at 125,000 cells per well. Oxygen consumption rate (OCR) was measured using the MitoStress assay on the Seahorse XFe96 Analyzer (Agilent Technologies), where cells were subjected to sequential exposure of oligomycin, FCCP, rotenone and antimycin A. Basal respiration, spare respiratory capacity and maximal mitochondria respiration were calculated, 3 biological triplicates with 10 technical replicates each were performed.

### Mass spectrometry protein enrichment analysis

R package clusterProfiler (v. 4.10.0) was used to perform gene ontology enrichment analysis of biological processes of the genes of 479 exclusive PSNP-bound proteins and KEGG pathway overrepresentation analysis of the genes of 1126 BJ PSNP batch 1 and 2 intersect proteins. For gene ontology enrichment analysis, the enrichGO() function was used with the following parameters: org.Hs.eg.db (R package v. 3.18.0) organism, ontology = BP, p-value cut-off = 0.01, q-value cut-off = 0.05, with Benjamini-Hochberg correction for adjusted p-values. Redundant terms were removed using R package enrichplot’s (v. 1.22.0) simplify() function. For KEGG pathway overrepresentation analysis, the enrichKEGG() function was used with the following parameters: Homo sapiens organism (hsa), p-value cut-off = 0.01, q-value cut-off = 0.05, with Benjamini-Hochberg correction for adjusted p-values. For both analyses, the top 10 enriched biological processes/pathways were visualized in a dot plot with enrichplot’s dotplot() function, ordered by gene ratio.

### In vivo motor neuron survival and soma size analyses

The soma sizes of ChAT+ motor neurons in the ventral spinal cord were quantified using ImageJ. The number of ChAT+ neurons per ventral horn was manually counted to assess motor neuron survival. Statistical analyses were conducted using GraphPad Prism.

### Data and statistical analysis

Statistical analyses were performed using graphpad Prism 7. Biological significance between two different groups were performed using unpaired Student’s t test. Comparison between more than two groups were calculated using one-way ANOVA followed by Dunn’s posthoc test. P < 0.05 was considered statistically significant.

## RESULTS

### Polystyrene nanoplastics are efficiently taken up by human neural cells

To investigate if polystyrene microplastics and nanoplastics can be internalized by human neural cells, we differentiated the human neural stem cell (NSC) line, ReN-VM, into neurons by withdrawal of mitogens bFGF and EGF [18], and treated these ReN-VM neurons with polystyrene spheres of varying diameters (50 nm, 500 nm and 5000 nm) at a concentration of 0.001% (w/v) for 6 hours (**Figure 1A**). Thereafter, cell culture media containing MP/NPs were aspirated, cultures were washed, and fresh media was replaced. Since the PS-MP/NP spheres possessed green fluorescence, internalization was determined by immunofluorescence and flow cytometry analyses, which revealed that 50 nm PS-NP spheres were most readily internalized and remains accumulated in the neurons for at least 48 hours after exposure (**Figures 1B-C**). To confirm the bio-accumulation of nanoplastics, ReN-VM cells were cultured as three-dimensional neural spheroids that were continuously exposed to 0.001% 50 nm PS-NP spheres for 72 hours. Spheroids were then harvested for cryosectioning and fluorescence imaging where the green fluorescent PS-NP were observed to penetrate into the core of the spheroids, which has a mean diameter of 720 μm (**Figures 1D-E**), indicating that 50 nm nanoplastics can be efficiently internalized by neural cells.

**Figure 1:**
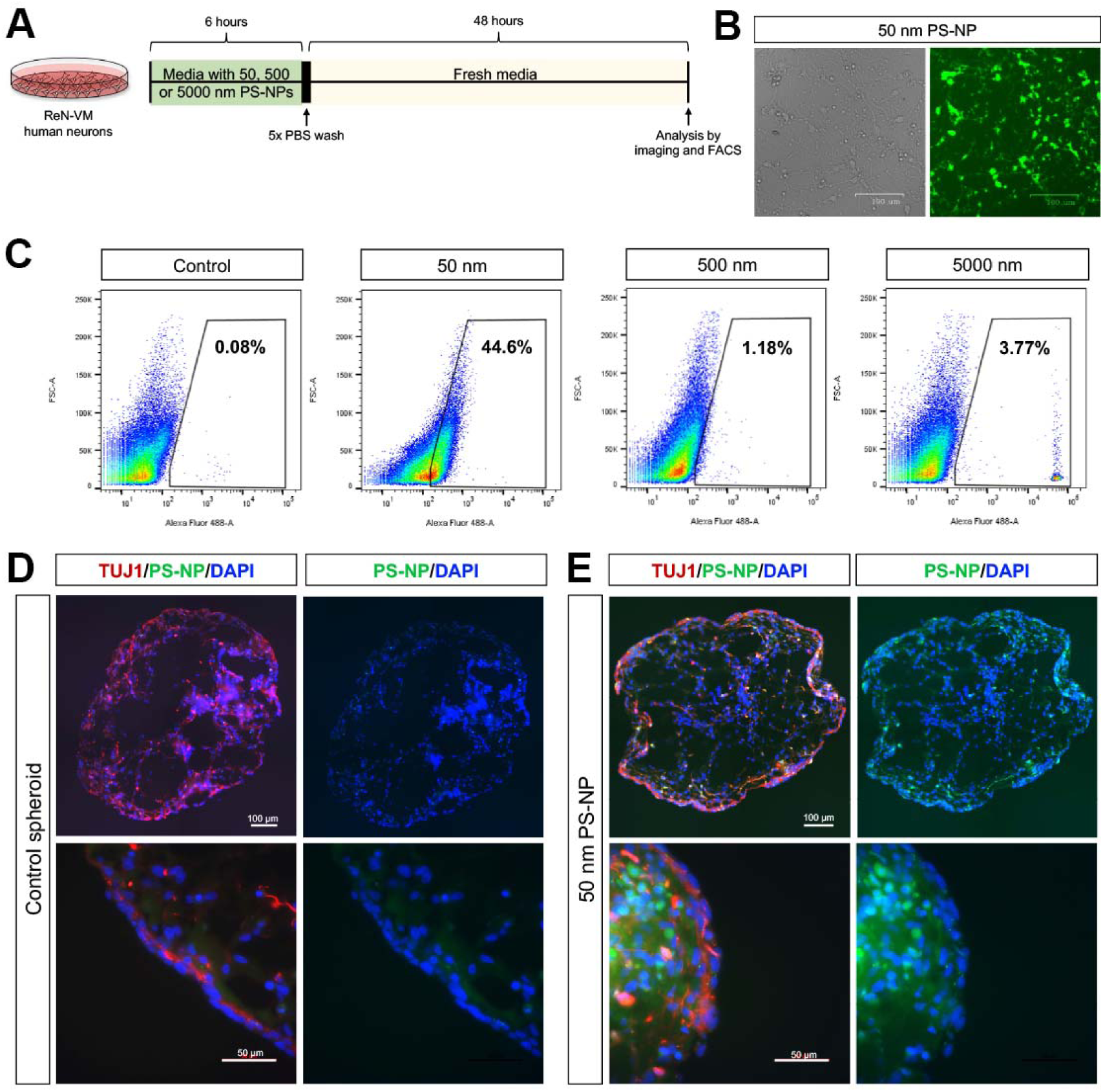
50-nm polystyrene nanoparticles are readily internalized by human neurons in vitro. (A) Treatment paradigm for polystyrene uptake studies using ReN-VM human neurons. (B) Fluorescent microscopy image of ReN-VM neurons treated with 50-nm PS-NPs after 48 hours of media washout. (C) Flow cytometry-based quantification of ReN-VM neurons that internalized the fluorescent PS-NPs of respective sizes. (D-E) ReN-VM neural stem cells were cultured in suspension as spheroids, and these spheroids were subsequently treated with PBS (control) or 50-nm PS-NPs respectively to measure uptake of PS-NPs in a three-dimensional context.

### Nanoplastics exposure reduced neural stem cell proliferation and induced neurotoxicity

To understand the effects of nanoplastics on human neurodevelopment and neuronal survival, we performed three assays: 1) NSC proliferation assay; 2) neurite length assay and 3) neuronal survival assay. In the NSC proliferation assay, ReN-VM NSCs were exposed to 50 nm PS-NP spheres at a concentration of 0.001% for 3 days and pulsed with EdU for 3 hours. Negative controls were incubated with 0.001% PBS instead. Cultures were then analyzed for incorporation of EdU, where it was determined that 50nm PS-NP treatment resulted in a slight but significant reduction of EdU incorporation (**Figures 2A, B**), indicating that exposure to 50 nm PS-NPs has a slight impact on cell proliferation.

**Figure 2:**
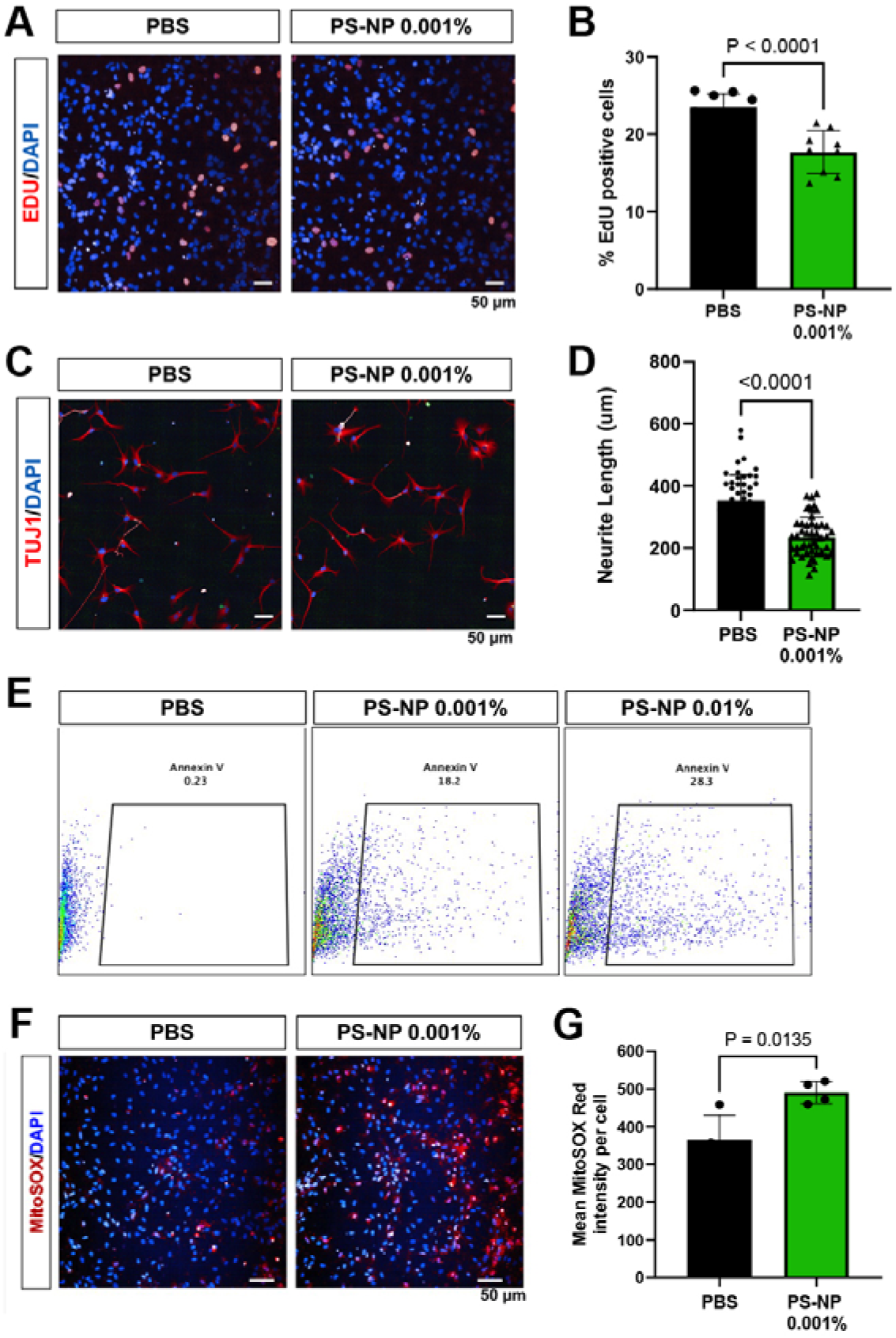
Exposure of nanoplastic reduce neuronal proliferation, neurite length and viability. (A) Fluorescence microscopy image of ReN-VM neurons showing EdU incorporation in red and cellular nuclei counterstained with DAPI in blue. (B) Quantification of EdU positive ReN-VM neurons treated with 50-nm PS-NPs or PBS (control); n = 9, p < 0.0001. (C) Immunofluorescent staining of ReN-VM neurons with neurites stained with TUJ1 in red and cellular nuclei counterstained with DAPI in blue. (D) Quantification of neurite length using neurite outgrowth plug-in from ImageJ, Neurite Tracer; n = 50, p < 0.0001. (E) ReN-VM NSCs were subjected to 2 concentrations of PS-NPs (0.01% and 0.001% respectively). Quantification of Ren-VM NSC apoptosis was performed by means of Annexin V flow-cytometry assay. (F) Fluorescence imaging of ReN-NM neurons depicting MitoSOX in red, and cellular nuclei counterstained with DAPI in blue. (G) Quantification of MitoSOX positive ReN-VM neurons upon PS-NP treatment.

To measure neurite lengths, ReN-VM NSCs were differentiated into neurons by removal of neurotrophic factors bFGF and EGF for 7 days [18], in the presence or absence of PS-NPs. Cultures were then fixed and stained with the neuronal marker beta-III-tubulin (TUJ1) and mean neurite lengths were measured. In both control and PS-NP-treated conditions, cell projections from neuronal somas were observed in almost all cells. A total of 100 neurons were analyzed where the longest projection from each neuron was traced and measured. We found that mean neurite length in control conditions were approximately 350 μm while neurons exposed to PS-NPs were about 225 μm (*p* < 0.0001) (**Figures 2C, D**). Since neurite length is associated with neurite outgrowth, neurotoxicity and general neuronal health, these results could suggest that the week-long PS-NP treatment significantly impaired healthy neuronal function.

Next, to investigate if nanoplastics have an effect on neuronal survival, ReN-VM neurons were exposed to 0.001% nanoplastics for 24 hours, and then analyzed for apoptosis using Annexin V conjugated with PE. Annexin V labeling was subsequently analyzed by flow cytometry. We found that exposure to 50 nm PS-NPs resulted in significantly higher number of neurons expressing Annexin V on their cell surface, with 18.2% of neurons undergoing apoptosis, compared with 0.23% in the control neurons. We also observed a dose-dependent effect on cell death where neurons exposed to a higher concentration of PS-NPs (0.01%) demonstrated more Annexin V-expressing cells (28.3%) (**Figure 2E**).

Previous studies have shown that PS-NPs induce cell death by generation of reactive oxygen species (ROS) and the activation of oxidative stress [19, 20]. Confirming the results from previous studies, we found that human neurons treated with 0.001% PS-NPs (50 nm) for 3 days had significantly higher levels of MitoSOX staining, providing further evidence that mitochondrial ROS is elevated in neurons exposed to PS-NPs (**Figure 2F, G**).

### Polystyrene nanoplastics interact with TDP43 and other proteins implicated in neurodegeneration

Protein misfolding and ER stress activation are molecular hallmarks of several neurodegenerative diseases [21], where misfolding and aggregation of intrinsically disordered proteins such as alpha synuclein, Tau and TDP43 drive neuronal dysfunctional and death. Since nanoplastics possess the ability to bind to proteins and alter their secondary structures [22], we reasoned that PS-NPs may interact with some of these disease-associated proteins and catalyze their aggregation. To investigate the physical interaction between PS-NPs and proteins commonly associated with neurodegenerative diseases, we performed a pulldown assay (**Figure 3A**) where neuronal cell lysates derived from BJ-iPS cells were incubated with 50nm PS-NPs for 30 minutes, followed by centrifugation to pellet the PS-NP spheres and its interacting proteins. The pellet is then washed with RIPA buffer and PBS to remove non-specific interactors. Eluate from the washed pellet was then analyzed by LC-MS/MS as well as SDS-PAGE and Western blot. As a positive control, we used silica nanoparticles since these were previously described to bind to neurodegeneration-associated proteins [23, 24].

**Figure 3:**
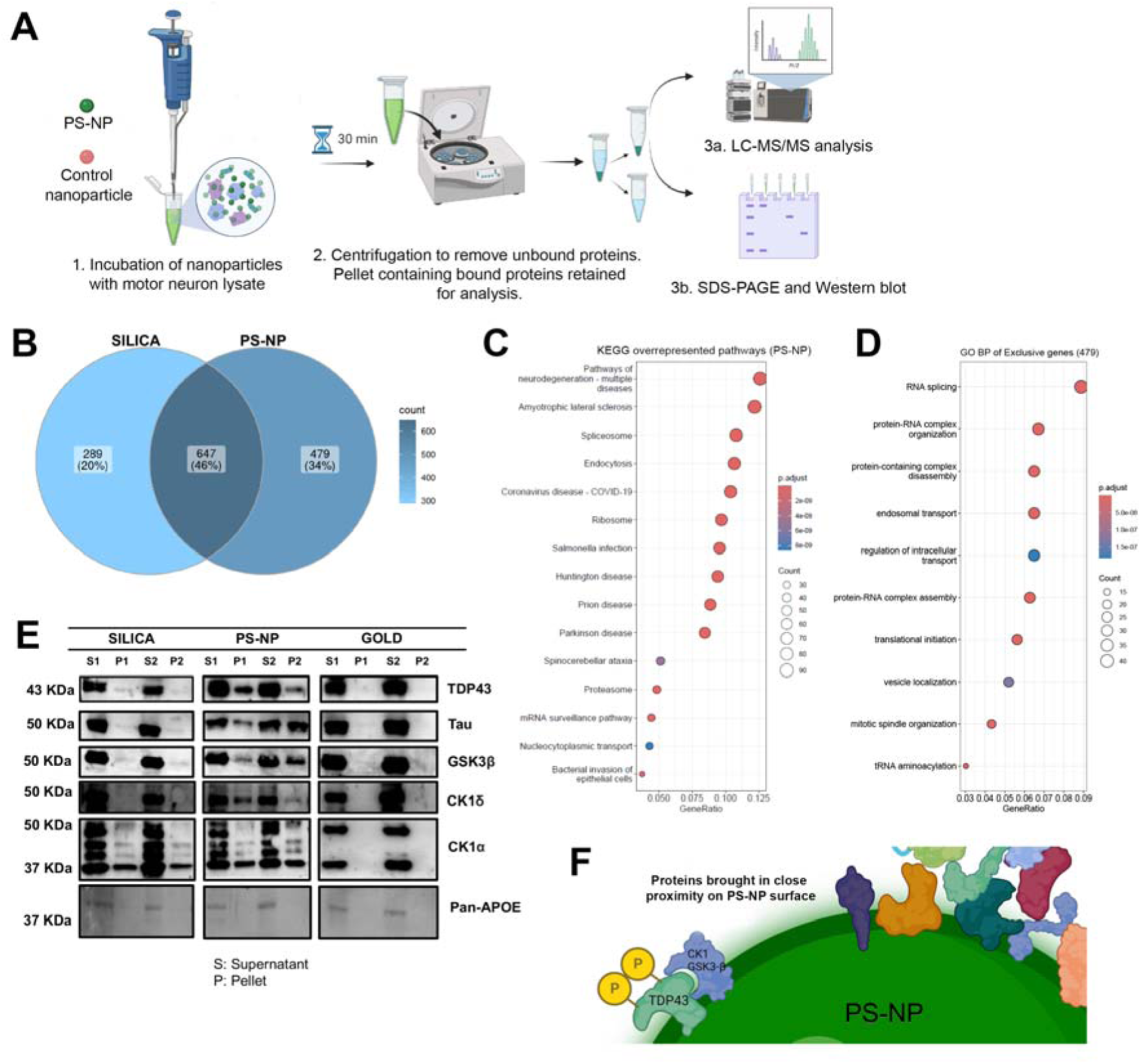
PS-NPs associate with TDP43 and other proteins implicated in neurodegeneration. (A) Schematic illustrating methodology for the nanoparticle pulldown assay used for identification of PS-NP binding partners. (B) Venn diagram analysis showing the overlap and unique proteins found to be interacting with silica (936) and PS-NPs (1126) using mass spectrometry. (C) Overrepresentation analysis of 1126 PS-NP binding partners categorized into pathways using KEGG database, p-value cut-off = 0.01, q-value cut-off = 0.05. (D) Gene ontology of top 10 biological processes identified based on 479 exclusive PS-NP bound proteins, p-value cut-off = 0.01, q-value cut-off = 0.05, with Benjamini-Hochberg correction for adjusted p-values. (E) Western blot validation of the mass spectrometry data, using inert gold nanoparticles as a negative control. (F) Working hypothesis where PS-NPs possibly behave as a physical catalyst that brings TDP43 and its kinases into close proximity, thereby promoting TDP43 phosphorylation.

To ensure reproducibility in the PS-NP pulldown data, two batches of pulldowns using either silica nanoparticles or PS-NPs were each performed, and the eluates were then analyzed by LC-MS/MS mass spectrometry. Our proteomics data revealed robust correlation between the two independent batches, where the Pearson’s correlation coefficient (r) value is greater than 0.6 (**Supplementary** Figure 1A) and the extent of overlap between both batches is larger than 50% (**Supplementary** Figure 1B). We reasoned that the intersect between both batches would represent high confidence interactions between silica nanoparticles or PS-NPs with their respective targets. This revealed that 936 proteins are bound by silica nanoparticles and 1126 proteins are bound by PS-NPs (FDR < 0.01) (**Figure 3B**). Silica nanoparticles have been previously shown to promote alpha synuclein aggregation and pathological features of Parkinson’s disease [24]. Our data now provides evidence of alpha synuclein interacting and co-precipitating with silica nanoparticles (**Supplementary Table 1**), validating that our pulldown approach is capable of identifying the protein interactors of nanoparticles, thereby elucidating biologically-relevant events.

Next, we analyzed the 1126 proteins that are bound by PS-NPs. Functional annotation by KEGG Pathway analysis revealed that proteins involved in ribosome, spliceosome, Amyotrophic Lateral Sclerosis (ALS), and pathways of neurodegeneration were significantly associated with PS-NPs (**Figure 3C**). Amongst this list of proteins, we also found that TDP43 and Tau are PS-NP interactors (**Supplementary Table 2**). Next, Gene Ontology analysis was performed on the 479 unique PS-NP-bound proteins. This revealed over-represented terms such as RNA splicing, ribonucleoprotein (RNP) complex biogenesis, organization, and assembly, as well as endosomal transport, protein localization and vesicle localization (**Figure 3D**), implying that PS-NPs were associated with these proteins and plausibly sequestering them away from their native roles.

To further validate that PS-NPs are bound to TDP43 and Tau, we performed Western blot on pulldown samples, using silica nanoparticles and 50-nm gold nanospheres as additional controls since the latter is presumably inert. We found that silica associates weakly with TDP43 but not Tau, while PS-NPs strongly associate with both TDP43 and Tau (**Figure 3E**). Meanwhile, a “negative control” protein such as ApoE that is not known to interact with PS-NPs or silica nanoparticles based on our mass spectrometry data, is not detected to be pulled down with PS-NPs, silica or gold nanoparticles (**Figure 3E**).

The over-represented Gene Ontology terms highlighting plausible errors in RNA metabolism and RNP function, coupled with the interactions of PS-NPs with TDP43 and Tau, suggest that PS-NPs could contribute to ALS, a disease characterized by motor neuron loss in the brain and spinal cord, where aberrant RNA metabolism is a major contributor to disease pathogenesis [25–27].Up to 90% of patients with sporadic ALS exhibit TDP43 pathology [28–30]. TDP43 is an RNA-binding protein that shuttles between the nucleus and cytosol, but remains mainly nuclear-localized in healthy cells and aberrantly accumulates in the cytoplasm under pathological conditions. The pathological hallmarks of TDP43 pathology therefore include cytoplasmic mislocalization, deposition of hyperphosphorylated TDP43 into inclusion bodies and eventual protein aggregation [30–34]. As TDP43 aggregation is associated with its phosphorylation on Serine 409/410, we first investigated if PS-NP exposure would result in elevated levels of phosphorylated TDP43 on Serine 409/410 (phospho-TDP43) in human neurons. To do so, we treated human ReN-VM neurons with 0.001% PS-NPs for 3 days, and probed for the levels of phospho-TDP43 compared to control samples treated with PBS (**Supplementary** Figure 2). This revealed elevated levels of phospho-TDP43 upon PS-NP exposure, implying that PS-NPs promoted the TDP43 pathology in human neurons.

### Polystyrene nanoplastics interact with kinases that phosphorylate TDP43

Next, we sought to investigate the relationship between PS-NPs and hyperphosphorylation of TDP43. A number of kinases have been identified to phosphorylate TDP43, and we postulate that PS-NPs could behave as a catalytic scaffold that brings TDP43 into close proximity with its kinases, possibly promoting its phosphorylation. To address this, we first analyzed the list of proteins that were found to interact with PS-NPs (**Supplementary Table 2**) and found that 40 of these proteins have kinase activities, including several serine/threonine protein kinases (**Supplementary Table 3**). Of these, we found that PS-NPs associate with GSK-3β, a proline-directed kinase that has been shown to phosphorylate Tau and TDP43 [35–37], as well as casein-kinase 1 alpha (CK-1α) and casein-kinase 2 (CK-2), which are known to phosphorylate TDP43 [33, 38]. We further validated this association using immunoblotting, and found that CK1α, CK1δ and GSK3β associate strongly with PS-NPs but not gold nanoparticles (**Figure 3E**). Therefore, this provides supporting evidence that PS-NPs bind to TDP43 and its kinases. Such interactions could contribute to conditions that favor TDP43 hyperphosphorylation, thereby inducing a neurodegenerative phenotype.(**Figure 3F**).

### Exposure to nanoplastics induce neurodegenerative phenotypes in human neurons

TDP43 hyperphosphorylation is a neuropathological hallmark of ALS. Expanding on our earlier findings that PS-NP exposure results in elevated phospho-TDP43 levels in human neurons (**Supplementary** Figure 2), we sought to investigate if PS-NPs would contribute to ALS-like neurodegeneration in physiologically-relevant models. Towards this end, we differentiated wild-type human iPSCs (BJ-iPS) into motor neurons using a 28-day protocol that is well-established in the lab [39], and exposed these cultures to 0.001% PS-NPs or PBS (control) for 3 or 7 days (**Figure 4A**). These neurons were then analyzed for various motor neuron parameters vis-à-vis an isogenic iPSC line with heterozygous TDP43^G298S^ mutation (BJ-TDP43) that we have previously validated to show well-defined ALS phenotypes [39].

**Figure 4:**
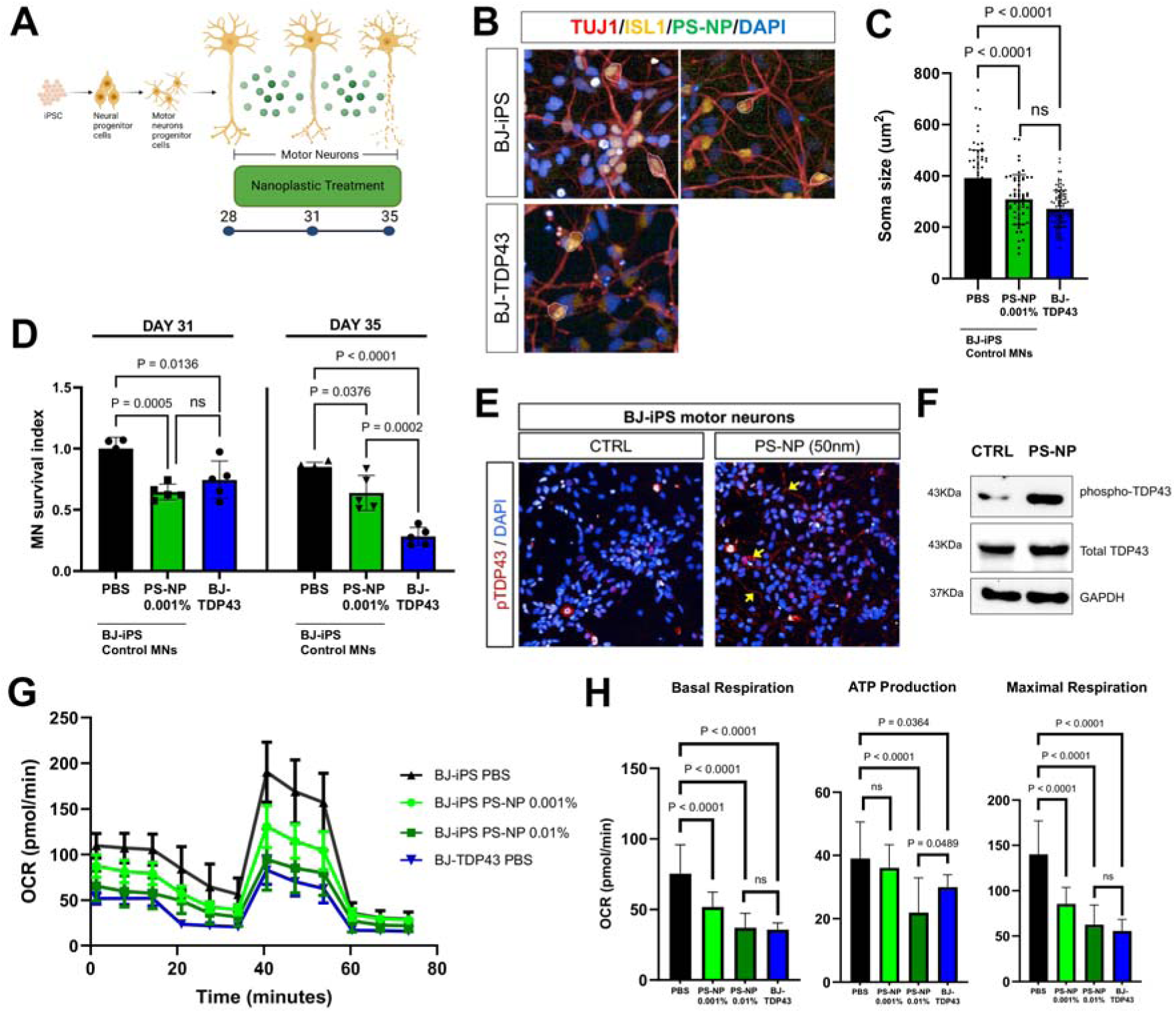
Exposure to PS-NPs induce neurodegenerative phenotypes in human neurons. (A) Schematic illustrating the PS-NP treatment paradigm using human iPSC-derived motor neurons. (B) Immunofluorescence of iPSC-derived motor neurons. Red: TUJ1, Yellow: ISL1, Green: PS-NPs and Blue: DAPI. (C) Quantification of motor neuron soma size treated with 50-nm PS-NPs or PBS (control) for 3 days. (D) Quantification of motor neuron survival treated with 50-nm PS-NPs or PBS (control) for 3 and 7 days respectively. (E) Immunofluorescence images of motor neurons stained for phosphorylated TDP43 (pTDP43). Red: pTDP43, Blue: DAPI. (F) Western blot analysis on probing for phosphorylated TDP-43 and total TDP43. (G) Oxygen consumption rate of wild-type and ALS neurons measured over time. The MitoStress assay was used to measure bioenergetics parameters, by adding oligomycin, FCCP, and a combination of rotenone and antimycin A (AA). **(H)** Quantification of basal respiration, ATP production, and maximal respiration.

First, ISL1+ motor neuron morphology was analyzed 3 days after exposure to 0.001% PS-NPs, at day 31. ALS motor neurons reportedly exhibit a reduction in soma sizes as disease progresses *in vitro* [39–41] and *in vivo* [42]. Likewise, we see a similar trend where the isogenic BJ-TDP43 motor neurons are consistently smaller than the wild-type motor neurons (**Figures 4B, C**). BJ-iPS motor neurons exposed to 0.001% PS-NPs for 3 days also exhibited significantly smaller soma sizes, decreasing from a mean of 392 μm^2^ to 308 µm^2^ (p < 0.001), equivalent to the size of BJ-TDP43 motor neurons at 271 µm^2^. Another disease phenotype associated with ALS is the acceleration of motor neuron death. To determine if motor neurons exposed to PS-NPs were less viable, we measured number of motor neurons at day 28 (baseline), day 31 (3 days of PS-NP exposure) and day 35 (7 days of PS-NP exposure) and compared this to baseline measurements of BJ-iPS motor neurons at day 28. We found that control BJ-iPS motor neurons that were untreated had minimal motor neuron death at day 31 and day 35 while BJ-TDP43 motor neurons had approximately 15.0% motor neuron death at day 31, and 71.7% death by day 35. Meanwhile, BJ-iPS motor neurons exposed to 0.001% PS-NPs exhibit 35.5% and 25.7% motor neuron loss on day 31 and 35 respectively, strongly suggesting that PS-NPs were contributing to motor neuron death (**Figure 4D**).

Using human ReN-VM neurons, we found that PS-NP exposure resulted in hyperphosphorylation of TDP43. Expanding on these earlier findings, we similarly examined the levels of phospho-TDP43 in BJ-iPS neurons exposed to PS-NPs for 3 days and found not just higher phospho-TDP43 levels but also presence of mislocalized TDP43 in the cytoplasm and neurites (**Figures 4E, F**), demonstrating an *in vitro* TDP43 pathology [43]. We had also previously established that mitochondrial dysfunction is a metabolic signature of ALS motor neurons [39]. Here, we found that exposure of healthy motor neurons to PS-NPs resulted in decreased mitochondrial respiration in a dose-dependent manner (**Figure 4G-H**). Taken together, this cumulative evidence strongly suggest that PS-NP exposure induces ALS-like phenotypes in human motor neurons.

### Exposure to nanoplastics induce neurodegenerative phenotypes in mice

Recent studies have provided compelling evidence that PS-NPs can cross the blood-brain barrier in rodents and accumulate in the brain, leading to various neurotoxic effects that potentiate neurodegeneration [15, 16]. While our *in vitro* data also supports the neurotoxic properties of PS-NPs, it is unclear if PS-NPs would induce ALS-like pathology in animal models. Towards this aim, we followed the PS-NP treatment paradigm from previously published papers [15, 16] where 38-week-old C57BL/6 mice were treated with 125 mg/kg/weekday of PS-NPs over the course of 8 weeks, and the spinal cords were harvested for histological and molecular analyses. During these 8 weeks, we found that PS-NP exposure did not induce significant changes to the body weights of mice (**Figure 5A**). Mice exposed to PS-NPs also did not display overt behavioral changes during the experimental time frame (data not shown). However, immunohistological analysis on mice spinal cords revealed that the ChAT+ motor neurons in the ventral horns of mice exposed to PS-NPs have significantly smaller soma sizes (mean of 661± 292 µm^2^ in control to 527± 296 µm^2^ in PS-NP group; p < 0.0001) (**Figures 5B, C**). Additionally, mice treated with PS-NPs also displayed fewer ChAT+ motor neurons per hemisection (**Figure 5D**). These findings strongly suggest that PS-NPs contribute to motor neuron degeneration in the spinal cord as fewer and smaller motor neurons are features of ALS in mice during the symptomatic and end stages [39–42].

**Figure 5:**
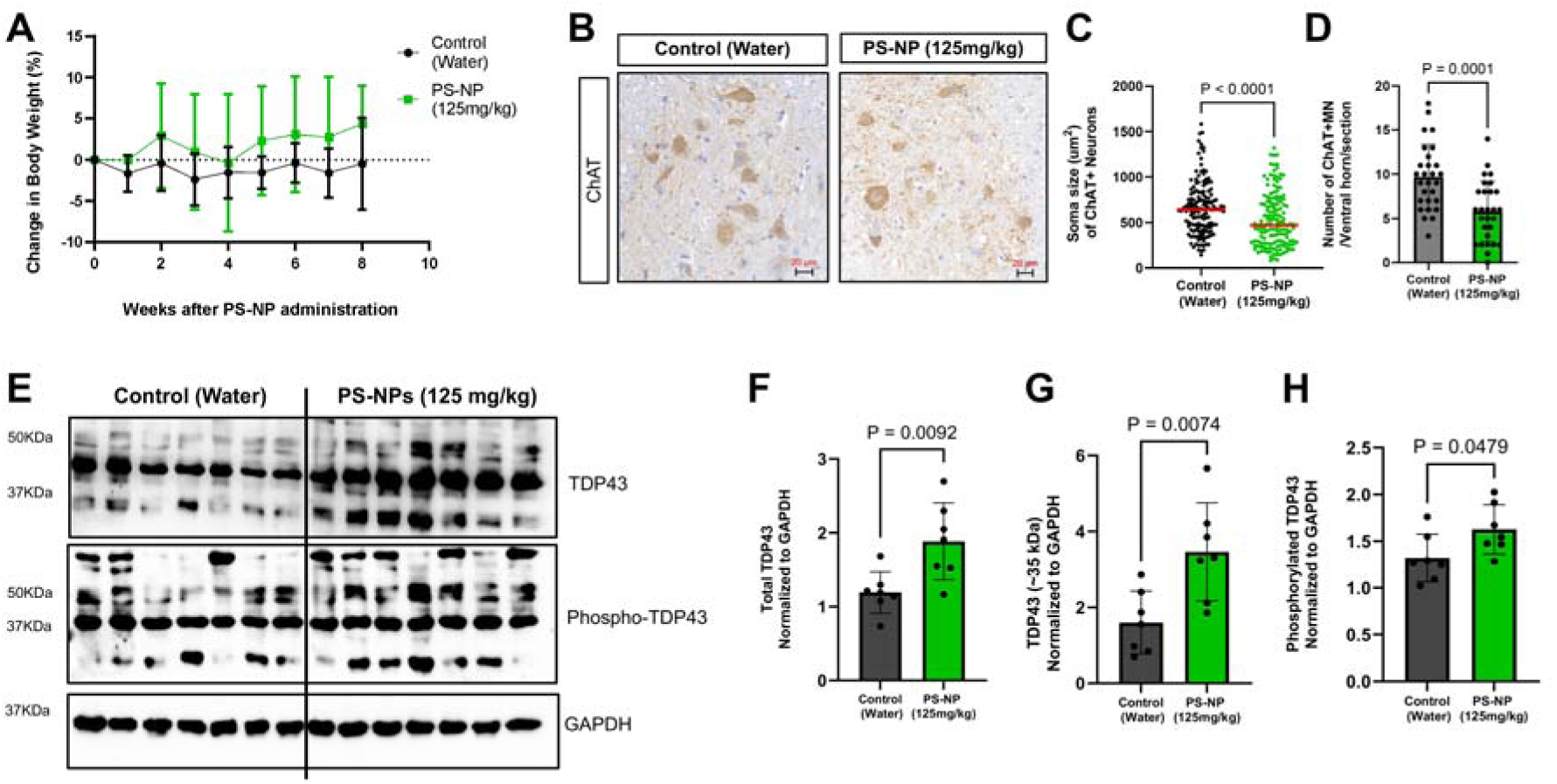
Mice exposed to PS-NPs (125 mg/kg/weekday) for 8 weeks demonstrate TDP43 pathologies. (A) Changes in mice weights over 8 weeks of PS-NP or control treatment, n = 8 each. (B) Representative immunohistochemistry images of the spinal cord ventral horn showing motor neurons labeled with ChAT. (C) Quantification of ventral horn motor neuron soma sizes in control and PS-NP groups. (D) Quantification of number of ChAT+ neurons per ventral horn per spinal cord section, n = 25 sections. (E) Western blot analysis of mice spinal cords probing for total TDP43, phospho-TDP43 and loading control GAPDH. (F-H) Quantification of various TDP43 isoforms.

Importantly, our *in vitro* data suggests that PS-NP exposure resulted in hyperphosphorylation of TDP43 (**Figure 4**), and we questioned if mice fed with PS-NPs would demonstrate TDP43 pathology. To investigate this, we first analyzed the various isoforms of TDP43 in the mice spinal cords by Western blot. Using a specific antibody against the C-terminus of TDP43, we detected higher levels of full-length TDP43 (43 kDa) as well as higher levels of a truncated form of TDP43 at 35 kDa in the PS-NP group (**Figures 5E-H**). Previous studies have shown that the 35 kDa C-truncated fragment of TDP43 causes cytoplasmic mislocalization and is linked to ALS pathology [44–46]. To detect phosphorylated TDP43, we analyzed the spinal cord lysates using a specific antibody that detects TDP43 phosphorylated on Serine 409/410 and likewise detected higher levels of phospho-TDP43 at 35 kDa (**Figures 5E, 5H**). These data collectively suggest that ingestion of PS-NPs contributed to TDP43 pathology and motor neuron degeneration.

## DISCUSSION

Nanoplastics are ubiquitous in our environment and especially in the food and water we consume, thus presenting a significant concern for public health. Amongst all plastic pollutants, polystyrene presents a major risk because it is one of the most common plastics used in food packaging. Studies in smaller mammals have shown that polystyrene nanoplastics (PS-NPs) penetrate the blood brain barrier and contribute to neuronal decline. Therefore, one of the goals of this study was to investigate the cellular and molecular events that PS-NPs trigger when they accumulate within human neurons so as to understand the exact mechanisms in which they potentiate neurodegeneration. Utilizing bona fide human neurons derived from induced pluripotent stem cells (iPSCs), we have identified critical interactions between PS-NPs and neuronal proteins, which led to proposed mechanisms through which PS-NPs influence neuronal pathology.

Our primary findings indicate that PS-NPs are readily assimilated by human neurons and bind to a myriad of proteins, which include proteins associated with neurodegenerative diseases such as Tau and TDP43. Intriguingly, we found that PS-NPs also interact with multiple kinases, including those that phosphorylate TDP43 such as CK1 and GSK3β. This led us to infer that PS-NPs may behave as a physical catalyst which brings the kinase and its substrate in close proximity, resulting in the hyperphosphorylation phenotype that we observed when human neurons were exposed to PS-NPs. This pathological change can result in the mislocalization of phospho-TDP43 to neuronal neurites, closely mirroring the cellular phenotypes observed in ALS. Overall, our *in vitro* data reveals a direct evidence between environmental PS-NPs pollution and TDP43 pathology, which has not been described before.

TDP43 pathology is a hallmark of ALS, an age-onset and progressive neurodegenerative disease where 90% of patients have the sporadic form of the disease. As such, we further explored the effects of PS-NPs on motor neurons, the primary cell type affected in ALS. In cultured human neurons, PS-NP exposure reduced neuronal viability and neuronal health parameters, resulting in impaired mitochondrial function and smaller cell bodies similar to phenotypes seen in ALS motor neuron cultures. Importantly, TDP43 hyperphosphorylation and mislocalization were also observed in neurons exposed to PS-NPs, providing evidence that PS-NPs contribute to TDP43 pathology, a critical mechanism in ALS pathophysiology. To verify that these *in vitro* results are physiologically relevant, wild-type “middle-aged” mice were fed PS-NPs over 8 weeks to model chronic PS-NP exposure, and analyses of their spinal cords revealed smaller and fewer motor neurons coupled with higher levels of cleaved and phosphorylated TDP43. Taken together, these sets of experiments support the notion that PS-NPs potentiate neurodegeneration by inducing TDP43 proteinopathy. While the mice do not exhibit motor dysfunction at the end of the 8-week PS-NP exposure period, it is possible that this exposure window is too short. It is plausible that increasing the PS-NP exposure time will lead to more pronounced ALS-like phenotypes, which is supported by the increased levels of truncated and hyperphosphorylated as well as immunohistological findings revealing fewer and smaller motor neurons at 8 weeks.

Our findings have several important implications. First, this study highlights the potential neurological risks posed by the ubiquitous presence of PS-NPs in our environment. Our work now provides direct evidence that motor neuron function and viability are impaired by PS-NPs, leading to ALS-like phenotypes. Second, it also expands upon previously published work by elucidating the PS-NP-protein interactome and a mechanism in which PS-NPs lead to neurodegeneration, where we proposed that PS-NPs behave as a catalyst that promote TDP43 phosphorylation. In doing so, our study has revealed a potential prevention and therapeutic target for neurodegenerative diseases induced by PS-NPs. It has been shown previously that inhibiting GSK3β and CK1 suppresses TDP43-mediated neurotoxicity [47–49]. Notably, GSK3β inhibition also reduces truncated TDP43 isoforms and ameliorates TDP43-induced toxicity, highlighting GSK3β and CK1 as likely preventive and therapeutic targets for PS-NP-induced neurotoxicity.

## Supporting information

https://drive.google.com/file/d/1Ktl-te45zux_MeJVcmIqPFKfBN-nzGBF/view?usp=sharing

## ACKNOWLEDGEMENTS

We are grateful to Dr Qinsong Lin (National University of Singapore) for his inputs with regards to the mass spectrometry experiments. We thank Advanced Molecular Pathology Lab at A*STAR for cryosectioning and histology, as well as the IMCB Flow Core Facility for flow cytometry-related work. We also thank IMCB for providing institutional and scientific equipment support. This work is supported by the following grants: National Research Foundation Fellowship (NRF-NRFF2018-003) awarded to Ng SY and A*STAR Career Development Award (212D800071) awarded to Winanto.

## Declaration of generative AI and AI-assisted technologies in the writing process

During the preparation of this work the author(s) used ChatGPT to assist with text revision and improve clarity. After using this tool/service, the author(s) reviewed and edited the content as needed and take(s) full responsibility for the content of the publication.

