## Supplementary material for "Polystyrene nanoplastics promote neurodegeneration by catalyzing TDP43 hyperphosphorylation": https://drive.google.com/file/d/1Ktl-te45zux_MeJVcmIqPFKfBN-nzGBF/view?usp=sharing

**This file includes:**

Supplementary Text
Figs. S1 to S2
Tables S1 to S3

**Supp. Fig. S1.**

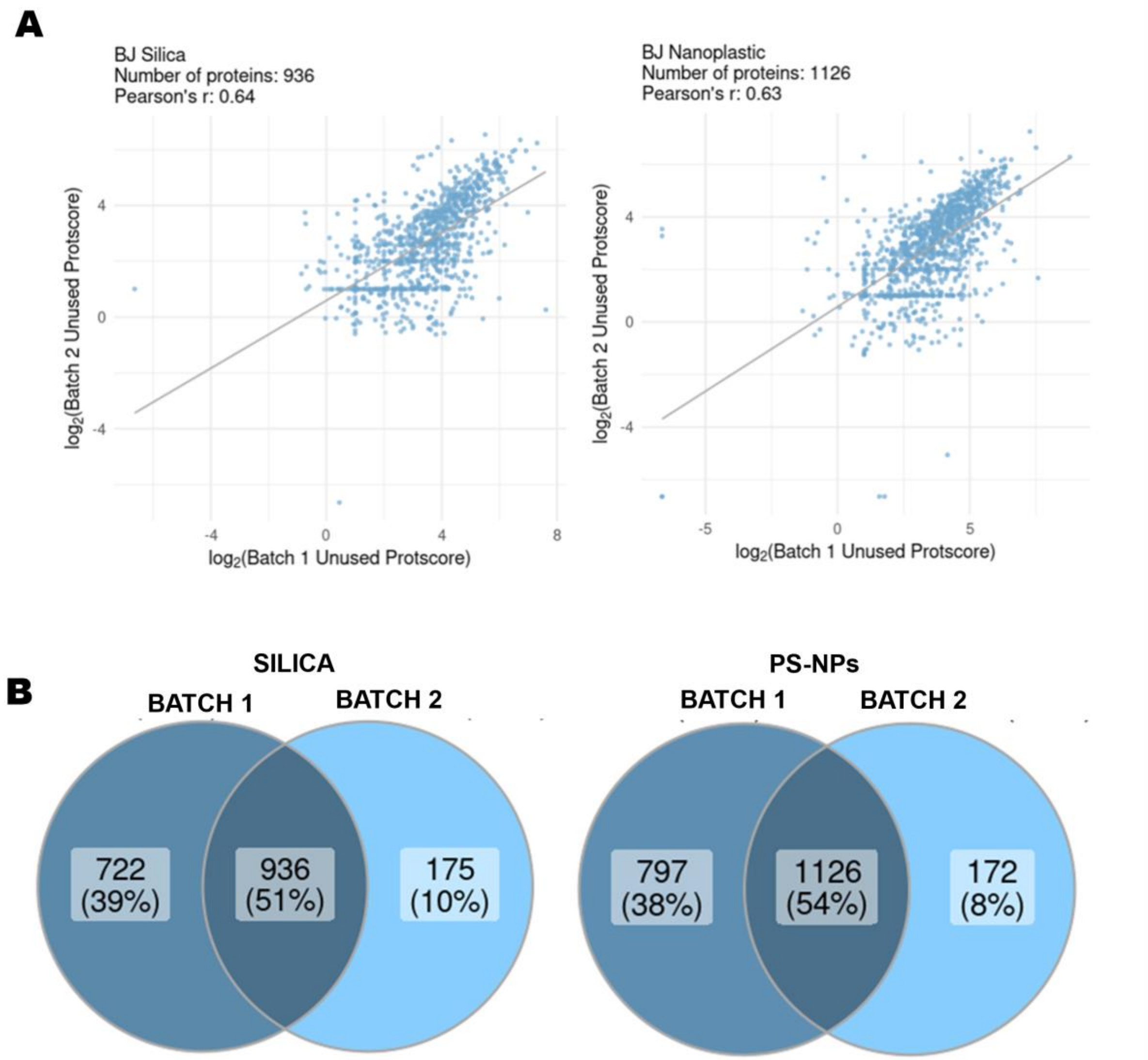


**Fig. S1. Reproducibility assessment of pulldown Assay.** (A) Scatter plots measuring the correlation between two batches of pulldown assays for Silica and PS-NPs respectively. Pearson correlation coefficient values reflecting the reproducibility of two replicates. Silica R= 0.64 and PS-NPs = 0.63. (B) Venn diagram showing the total number of proteins identified by ProteinPilot In pulldown assays for both experimental replicates. For Silica, a total number of 1658 were detected in Batch 1 and 1111 proteins were identified in Batch 2, of which 936 proteins were detected in both batches. For PS-NPs, a total number of 1923 and 1298 were detected in the two replicates, of which 1126 proteins were common between the two batches.

**Supp. Fig. S2.**

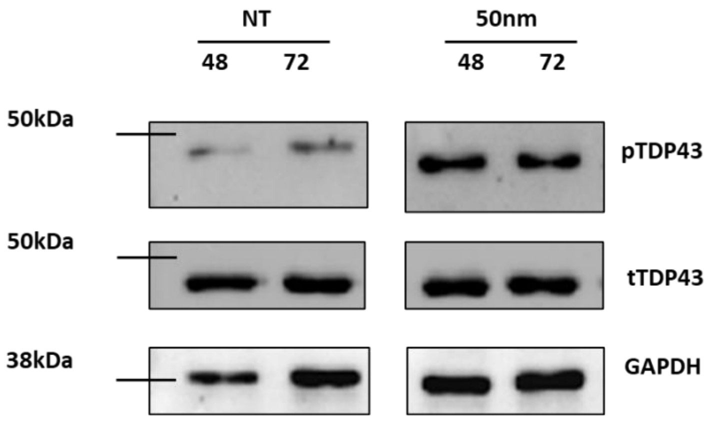


**Fig. S2. Human ReN-VM neurons exposed to PS-NPs show increased phosphorylation of TDP43**. Western blot analysis of the levels of phosphorylated TDP43 (pTDP43) and total TDP43 (tTDP43) expression after PS-NP treatment. Ren-VM neurons were subjected to 50 nm PS-NPs or PBS for 48 and 72 hours before measuring pTDP43 levels.

**Supp. Table S1.**

**Supplementary Table S1: List of 936 proteins bound by silica, detected by LC-MS/MS**

| Accession | Protein Name | Species |
|---|---|---|
| sp\|Q14204\|DYHC1_HUMAN | Cytoplasmic dynein 1 heavy chain 1 | HUMAN |
| sp\|P35580\|MYH10_HUMAN | Myosin-10 | HUMAN |
| sp\|Q00610\|CLH1_HUMAN | Clathrin heavy chain 1 | HUMAN |
| sp\|Q13813\|SPTN1_HUMAN | Spectrin alpha chain, non-erythrocytic 1 | HUMAN |
| sp\|P46821\|MAP1B_HUMAN | Microtubule-associated protein 1B | HUMAN |
| sp\|P52272\|HNRPM_HUMAN | Heterogeneous nuclear ribonucleoprotein M | HUMAN |
| sp\|P07437\|TBB5_HUMAN | Tubulin beta chain | HUMAN |
| sp\|Q71U36\|TBA1A_HUMAN | Tubulin alpha-1A chain | HUMAN |
| sp\|P13639\|EF2_HUMAN | Elongation factor 2 | HUMAN |
| sp\|P43243\|MATR3_HUMAN | Matrin-3 | HUMAN |
| sp\|P08238\|HS90B_HUMAN | Heat shock protein HSP 90-beta | HUMAN |
| sp\|Q13263\|TIF1B_HUMAN | Transcription intermediary factor 1-beta | HUMAN |
| sp\|Q08211\|DHX9_HUMAN | ATP-dependent RNA helicase A | HUMAN |
| sp\|O75643\|U520_HUMAN | U5 small nuclear ribonucleoprotein 200 kDa helicase | HUMAN |
| sp\|Q92945\|FUBP2_HUMAN | Far upstream element-binding protein 2 | HUMAN |
| sp\|P14618\|KPYM_HUMAN | Pyruvate kinase PKM | HUMAN |
| sp\|P11142\|HSP7C_HUMAN | Heat shock cognate 71 kDa protein | HUMAN |
| sp\|P35908\|K22E_HUMAN | Keratin, type II cytoskeletal 2 epidermal | HUMAN |
| sp\|P12956\|XRCC6_HUMAN | X-ray repair cross-complementing protein 6 | HUMAN |
| sp\|P68104\|EF1A1_HUMAN | Elongation factor 1-alpha 1 | HUMAN |
| sp\|P13645\|K1C10_HUMAN | Keratin, type I cytoskeletal 10 | HUMAN |
| sp\|P78527\|PRKDC_HUMAN | DNA-dependent protein kinase catalytic subunit | HUMAN |
| sp\|Q07954\|LRP1_HUMAN | Prolow-density lipoprotein receptor-related protein 1 | HUMAN |
| sp\|P22626\|ROA2_HUMAN | Heterogeneous nuclear ribonucleoproteins A2/B1 | HUMAN |
| sp\|P17844\|DDX5_HUMAN | Probable ATP-dependent RNA helicase DDX5 | HUMAN |
| sp\|P61978\|HNRPK_HUMAN | Heterogeneous nuclear ribonucleoprotein K | HUMAN |
| sp\|P63261\|ACTG_HUMAN | Actin, cytoplasmic 2 | HUMAN |
| sp\|Q96AE4\|FUBP1_HUMAN | Far upstream element-binding protein 1 | HUMAN |
| sp\|Q12906\|ILF3_HUMAN | Interleukin enhancer-binding factor 3 | HUMAN |
| sp\|P23246\|SFPQ_HUMAN | Splicing factor, proline- and glutamine-rich | HUMAN |
| sp\|P13637\|AT1A3_HUMAN | Sodium/potassium-transporting ATPase subunit alpha-3 | HUMAN |
| sp\|P78344\|IF4G2_HUMAN | Eukaryotic translation initiation factor 4 gamma 2 | HUMAN |
| sp\|P04264\|K2C1_HUMAN | Keratin, type II cytoskeletal 1 | HUMAN |
| sp\|P46459\|NSF_HUMAN | Vesicle-fusing ATPase | HUMAN |

| | | |
|---|---|---|
| sp\|P11717\|MPRI_HUMAN | Cation-independent mannose-6-phosphate receptor | HUMAN |
| sp\|P13010\|XRCC5_HUMAN | X-ray repair cross-complementing protein 5 | HUMAN |
| sp\|Q00839\|HNRPU_HUMAN | Heterogeneous nuclear ribonucleoprotein U | HUMAN |
| sp\|P11940\|PABP1_HUMAN | Polyadenylate-binding protein 1 | HUMAN |
| sp\|P21333\|FLNA_HUMAN | Filamin-A | HUMAN |
| sp\|P29401\|TKT_HUMAN | Transketolase | HUMAN |
| sp\|P22314\|UBA1_HUMAN | Ubiquitin-like modifier-activating enzyme 1 | HUMAN |
| sp\|Q9NZI8\|IF2B1_HUMAN | Insulin-like growth factor 2 mRNA-binding protein 1 | HUMAN |
| sp\|O43390\|HNRPR_HUMAN | Heterogeneous nuclear ribonucleoprotein R | HUMAN |
| sp\|P63010\|AP2B1_HUMAN | AP-2 complex subunit beta | HUMAN |
| sp\|Q15459\|SF3A1_HUMAN | Splicing factor 3A subunit 1 | HUMAN |
| sp\|P55072\|TERA_HUMAN | Transitional endoplasmic reticulum ATPase | HUMAN |
| sp\|P50395\|GDIB_HUMAN | Rab GDP dissociation inhibitor beta | HUMAN |
| sp\|P31948\|STIP1_HUMAN | Stress-induced-phosphoprotein 1 | HUMAN |
| sp\|P31943\|HNRH1_HUMAN | Heterogeneous nuclear ribonucleoprotein H | HUMAN |
| sp\|P27824\|CALX_HUMAN | Calnexin | HUMAN |
| sp\|P26038\|MOES_HUMAN | Moesin | HUMAN |
| sp\|P04075\|ALDOA_HUMAN | Fructose-bisphosphate aldolase A | HUMAN |
| sp\|P30101\|PDIA3_HUMAN | Protein disulfide-isomerase A3 | HUMAN |
| sp\|Q15393\|SF3B3_HUMAN | Splicing factor 3B subunit 3 | HUMAN |
| sp\|Q92499\|DDX1_HUMAN | ATP-dependent RNA helicase DDX1 | HUMAN |
| sp\|P11137\|MTAP2_HUMAN | Microtubule-associated protein 2 | HUMAN |
| sp\|Q15029\|U5S1_HUMAN | 116 kDa U5 small nuclear ribonucleoprotein component | HUMAN |
| sp\|Q16555\|DPYL2_HUMAN | Dihydropyrimidinase-related protein 2 | HUMAN |
| sp\|Q15233\|NONO_HUMAN | Non-POU domain-containing octamer-binding protein | HUMAN |
| sp\|P08670\|VIME_HUMAN | Vimentin | HUMAN |
| sp\|P78371\|TCPB_HUMAN | T-complex protein 1 subunit beta | HUMAN |
| sp\|Q16643\|DREB_HUMAN | Drebrin | HUMAN |
| sp\|P50990\|TCPQ_HUMAN | T-complex protein 1 subunit theta | HUMAN |
| sp\|Q9P2K5\|MYEF2_HUMAN | Myelin expression factor 2 | HUMAN |
| sp\|P11021\|BIP_HUMAN | Endoplasmic reticulum chaperone BiP | HUMAN |
| sp\|P19338\|NUCL_HUMAN | Nucleolin | HUMAN |
| sp\|P78347\|GTF2I_HUMAN | General transcription factor II-I | HUMAN |
| sp\|Q01082\|SPTB2_HUMAN | Spectrin beta chain, non-erythrocytic 1 | HUMAN |
| sp\|P06733\|ENOA_HUMAN | Alpha-enolase | HUMAN |
| sp\|P14625\|ENPL_HUMAN | Endoplasmin | HUMAN |
| sp\|P13591\|NCAM1_HUMAN | Neural cell adhesion molecule 1 | HUMAN |
| sp\|Q9Y3I0\|RTCB_HUMAN | RNA-splicing ligase RtcB homolog | HUMAN |
| sp\|P26599\|PTBP1_HUMAN | Polypyrimidine tract-binding protein 1 | HUMAN |

| | | |
|---|---|---|
| sp\|Q8N163\|CCAR2_HUMAN | Cell cycle and apoptosis regulator protein 2 | HUMAN |
| sp\|O00571\|DDX3X_HUMAN | ATP-dependent RNA helicase DDX3X | HUMAN |
| sp\|P62258\|1433E_HUMAN | 14-3-3 protein epsilon | HUMAN |
| sp\|P14866\|HNRPL_HUMAN | Heterogeneous nuclear ribonucleoprotein L | HUMAN |
| sp\|P38919\|IF4A3_HUMAN | Eukaryotic initiation factor 4A-III | HUMAN |
| sp\|Q12926\|ELAV2_HUMAN | ELAV-like protein 2 | HUMAN |
| sp\|Q9Y6M1\|IF2B2_HUMAN | Insulin-like growth factor 2 mRNA-binding protein 2 | HUMAN |
| sp\|Q99536\|VAT1_HUMAN | Synaptic vesicle membrane protein VAT-1 homolog | HUMAN |
| sp\|P00558\|PGK1_HUMAN | Phosphoglycerate kinase 1 | HUMAN |
| sp\|P12277\|KCRB_HUMAN | Creatine kinase B-type | HUMAN |
| sp\|Q14157\|UBP2L_HUMAN | Ubiquitin-associated protein 2-like | HUMAN |
| sp\|P35527\|K1C9_HUMAN | Keratin, type I cytoskeletal 9 | HUMAN |
| sp\|Q13435\|SF3B2_HUMAN | Splicing factor 3B subunit 2 | HUMAN |
| sp\|Q9BPU6\|DPYL5_HUMAN | Dihydropyrimidinase-related protein 5 | HUMAN |
| sp\|Q9UHX1\|PUF60_HUMAN | Poly(U)-binding-splicing factor PUF60 | HUMAN |
| sp\|P50454\|SERPH_HUMAN | Serpin H1 | HUMAN |
| sp\|Q9UQ80\|PA2G4_HUMAN | Proliferation-associated protein 2G4 | HUMAN |
| sp\|P09651\|ROA1_HUMAN | Heterogeneous nuclear ribonucleoprotein A1 | HUMAN |
| sp\|P07900\|HS90A_HUMAN | Heat shock protein HSP 90-alpha | HUMAN |
| sp\|Q9NQC3\|RTN4_HUMAN | Reticulon-4 | HUMAN |
| sp\|Q14195\|DPYL3_HUMAN | Dihydropyrimidinase-related protein 3 | HUMAN |
| sp\|P23396\|RS3_HUMAN | 40S ribosomal protein S3 | HUMAN |
| sp\|Q9Y230\|RUVB2_HUMAN | RuvB-like 2 | HUMAN |
| sp\|Q14980\|NUMA1_HUMAN | Nuclear mitotic apparatus protein 1 | HUMAN |
| sp\|P04406\|G3P_HUMAN | Glyceraldehyde-3-phosphate dehydrogenase | HUMAN |
| sp\|Q12905\|ILF2_HUMAN | Interleukin enhancer-binding factor 2 | HUMAN |
| sp\|Q8NC51\|PAIRB_HUMAN | Plasminogen activator inhibitor 1 RNA-binding protein | HUMAN |
| sp\|P0DMV9\|HS71B_HUMAN | Heat shock 70 kDa protein 1B | HUMAN |
| sp\|Q92841\|DDX17_HUMAN | Probable ATP-dependent RNA helicase DDX17 | HUMAN |
| sp\|P07910\|HNRPC_HUMAN | Heterogeneous nuclear ribonucleoproteins C1/C2 | HUMAN |
| sp\|O00425\|IF2B3_HUMAN | Insulin-like growth factor 2 mRNA-binding protein 3 | HUMAN |
| sp\|P49368\|TCPG_HUMAN | T-complex protein 1 subunit gamma | HUMAN |
| sp\|P38159\|RBMX_HUMAN | RNA-binding motif protein, X chromosome | HUMAN |
| sp\|Q14697\|GANAB_HUMAN | Neutral alpha-glucosidase AB | HUMAN |
| sp\|O14531\|DPYL4_HUMAN | Dihydropyrimidinase-related protein 4 | HUMAN |
| sp\|Q13838\|DX39B_HUMAN | Spliceosome RNA helicase DDX39B | HUMAN |
| sp\|P61011\|SRP54_HUMAN | Signal recognition particle 54 kDa protein | HUMAN |

| | | |
|---|---|---|
| sp\|Q15365\|PCBP1_HUMAN | Poly(rC)-binding protein 1 | HUMAN |
| sp\|P40227\|TCPZ_HUMAN | T-complex protein 1 subunit zeta | HUMAN |
| sp\|P18124\|RL7_HUMAN | 60S ribosomal protein L7 | HUMAN |
| sp\|Q9BWF3\|RBM4_HUMAN | RNA-binding protein 4 | HUMAN |
| sp\|P53621\|COPA_HUMAN | Coatomer subunit alpha | HUMAN |
| sp\|P35606\|COPB2_HUMAN | Coatomer subunit beta' | HUMAN |
| sp\|P22234\|PUR6_HUMAN | Bifunctional phosphoribosylaminoimidazole carboxylase/phosphoribosylaminoimidazole succinocarboxamide synthetase | HUMAN |
| sp\|Q14194\|DPYL1_HUMAN | Dihydropyrimidinase-related protein 1 | HUMAN |
| sp\|Q15293\|RCN1_HUMAN | Reticulocalbin-1 | HUMAN |
| sp\|P48643\|TCPE_HUMAN | T-complex protein 1 subunit epsilon | HUMAN |
| sp\|P36578\|RL4_HUMAN | 60S ribosomal protein L4 | HUMAN |
| sp\|P09874\|PARP1_HUMAN | Poly [ADP-ribose] polymerase 1 | HUMAN |
| sp\|P62701\|RS4X_HUMAN | 40S ribosomal protein S4, X isoform | HUMAN |
| sp\|Q13509\|TBB3_HUMAN | Tubulin beta-3 chain | HUMAN |
| sp\|P63244\|RACK1_HUMAN | Receptor of activated protein C kinase 1 | HUMAN |
| sp\|Q9Y265\|RUVB1_HUMAN | RuvB-like 1 | HUMAN |
| sp\|Q99615\|DNJC7_HUMAN | DnaJ homolog subfamily C member 7 | HUMAN |
| sp\|P10809\|CH60_HUMAN | 60 kDa heat shock protein, mitochondrial | HUMAN |
| sp\|Q9UMS4\|PRP19_HUMAN | Pre-mRNA-processing factor 19 | HUMAN |
| sp\|Q99832\|TCPH_HUMAN | T-complex protein 1 subunit eta | HUMAN |
| sp\|P62424\|RL7A_HUMAN | 60S ribosomal protein L7a | HUMAN |
| sp\|Q14677\|EPN4_HUMAN | Clathrin interactor 1 | HUMAN |
| sp\|Q02878\|RL6_HUMAN | 60S ribosomal protein L6 | HUMAN |
| sp\|O43602\|DCX_HUMAN | Neuronal migration protein doublecortin | HUMAN |
| sp\|O76070\|SYUG_HUMAN | Gamma-synuclein | HUMAN |
| sp\|P61247\|RS3A_HUMAN | 40S ribosomal protein S3a | HUMAN |
| sp\|P08865\|RSSA_HUMAN | 40S ribosomal protein SA | HUMAN |
| sp\|P15880\|RS2_HUMAN | 40S ribosomal protein S2 | HUMAN |
| sp\|P35222\|CTNB1_HUMAN | Catenin beta-1 | HUMAN |
| sp\|Q1KMD3\|HNRL2_HUMAN | Heterogeneous nuclear ribonucleoprotein U-like protein 2 | HUMAN |
| sp\|Q13561\|DCTN2_HUMAN | Dynactin subunit 2 | HUMAN |
| sp\|Q9NTZ6\|RBM12_HUMAN | RNA-binding protein 12 | HUMAN |
| sp\|P52597\|HNRPF_HUMAN | Heterogeneous nuclear ribonucleoprotein F | HUMAN |
| sp\|P39023\|RL3_HUMAN | 60S ribosomal protein L3 | HUMAN |
| sp\|P05388\|RLA0_HUMAN | 60S acidic ribosomal protein P0 | HUMAN |
| sp\|P35221\|CTNA1_HUMAN | Catenin alpha-1 | HUMAN |
| sp\|P62820\|RAB1A_HUMAN | Ras-related protein Rab-1A | HUMAN |
| sp\|P06744\|G6PI_HUMAN | Glucose-6-phosphate isomerase | HUMAN |
| sp\|P17677\|NEUM_HUMAN | Neuromodulin | HUMAN |
| sp\|Q92900\|RENT1_HUMAN | Regulator of nonsense transcripts 1 | HUMAN |

| | | |
|---|---|---|
| sp\|P07195\|LDHB_HUMAN | L-lactate dehydrogenase B chain | HUMAN |
| sp\|Q07955\|SRSF1_HUMAN | Serine/arginine-rich splicing factor 1 | HUMAN |
| sp\|P26641\|EF1G_HUMAN | Elongation factor 1-gamma | HUMAN |
| sp\|P60842\|IF4A1_HUMAN | Eukaryotic initiation factor 4A-I | HUMAN |
| sp\|Q15637\|SF01_HUMAN | Splicing factor 1 | HUMAN |
| sp\|P60174\|TPIS_HUMAN | Triosephosphate isomerase | HUMAN |
| sp\|P17987\|TCPA_HUMAN | T-complex protein 1 subunit alpha | HUMAN |
| sp\|Q8N6H7\|ARFG2_HUMAN | ADP-ribosylation factor GTPase-activating protein 2 | HUMAN |
| sp\|Q13148\|TADBP_HUMAN | TAR DNA-binding protein 43 | HUMAN |
| sp\|Q14103\|HNRPD_HUMAN | Heterogeneous nuclear ribonucleoprotein D0 | HUMAN |
| sp\|P67809\|YBOX1_HUMAN | Y-box-binding protein 1 | HUMAN |
| sp\|Q96CW1\|AP2M1_HUMAN | AP-2 complex subunit mu | HUMAN |
| sp\|P13647\|K2C5_HUMAN | Keratin, type II cytoskeletal 5 | HUMAN |
| sp\|O43347\|MSI1H_HUMAN | RNA-binding protein Musashi homolog 1 | HUMAN |
| sp\|P39019\|RS19_HUMAN | 40S ribosomal protein S19 | HUMAN |
| sp\|Q9UNW9\|NOVA2_HUMAN | RNA-binding protein Nova-2 | HUMAN |
| sp\|O60282\|KIF5C_HUMAN | Kinesin heavy chain isoform 5C | HUMAN |
| sp\|P25685\|DNJB1_HUMAN | DnaJ homolog subfamily B member 1 | HUMAN |
| sp\|O43852\|CALU_HUMAN | Calumenin | HUMAN |
| sp\|P16949\|STMN1_HUMAN | Stathmin | HUMAN |
| sp\|P06748\|NPM_HUMAN | Nucleophosmin | HUMAN |
| sp\|P09471\|GNAO_HUMAN | Guanine nucleotide-binding protein G(o) subunit alpha | HUMAN |
| sp\|Q16531\|DDB1_HUMAN | DNA damage-binding protein 1 | HUMAN |
| sp\|Q16658\|FSCN1_HUMAN | Fascin | HUMAN |
| sp\|P08621\|RU17_HUMAN | U1 small nuclear ribonucleoprotein 70 kDa | HUMAN |
| sp\|Q14498\|RBM39_HUMAN | RNA-binding protein 39 | HUMAN |
| sp\|P51610\|HCFC1_HUMAN | Host cell factor 1 | HUMAN |
| sp\|O43809\|CPSF5_HUMAN | Cleavage and polyadenylation specificity factor subunit 5 | HUMAN |
| sp\|P63104\|1433Z_HUMAN | 14-3-3 protein zeta/delta | HUMAN |
| sp\|P46777\|RL5_HUMAN | 60S ribosomal protein L5 | HUMAN |
| sp\|P37268\|FDFT_HUMAN | Squalene synthase | HUMAN |
| sp\|P62249\|RS16_HUMAN | 40S ribosomal protein S16 | HUMAN |
| sp\|P51991\|ROA3_HUMAN | Heterogeneous nuclear ribonucleoprotein A3 | HUMAN |
| sp\|P29762\|RABP1_HUMAN | Cellular retinoic acid-binding protein 1 | HUMAN |
| sp\|Q8WXF1\|PSPC1_HUMAN | Paraspeckle component 1 | HUMAN |
| sp\|Q15417\|CNN3_HUMAN | Calponin-3 | HUMAN |
| sp\|Q9UKA9\|PTBP2_HUMAN | Polypyrimidine tract-binding protein 2 | HUMAN |
| sp\|Q9NQG5\|RPR1B_HUMAN | Regulation of nuclear pre-mRNA domain-containing protein 1B | HUMAN |
| sp\|Q13177\|PAK2_HUMAN | Serine/threonine-protein kinase PAK 2 | HUMAN |

| | | |
|---|---|---|
| sp\|Q5SSJ5\|HP1B3_HUMAN | Heterochromatin protein 1-binding protein 3 | HUMAN |
| sp\|P46781\|RS9_HUMAN | 40S ribosomal protein S9 | HUMAN |
| sp\|Q9UMX0\|UBQL1_HUMAN | Ubiquilin-1 | HUMAN |
| sp\|P31942\|HNRH3_HUMAN | Heterogeneous nuclear ribonucleoprotein H3 | HUMAN |
| sp\|P05455\|LA_HUMAN | Lupus La protein | HUMAN |
| sp\|P62937\|PPIA_HUMAN | Peptidyl-prolyl cis-trans isomerase A | HUMAN |
| sp\|P23284\|PPIB_HUMAN | Peptidyl-prolyl cis-trans isomerase B | HUMAN |
| sp\|Q96PK6\|RBM14_HUMAN | RNA-binding protein 14 | HUMAN |
| sp\|P02786\|TFR1_HUMAN | Transferrin receptor protein 1 | HUMAN |
| sp\|P61106\|RAB14_HUMAN | Ras-related protein Rab-14 | HUMAN |
| sp\|P80723\|BASP1_HUMAN | Brain acid soluble protein 1 | HUMAN |
| sp\|P27816\|MAP4_HUMAN | Microtubule-associated protein 4 | HUMAN |
| sp\|P48444\|COPD_HUMAN | Coatomer subunit delta | HUMAN |
| sp\|P61981\|1433G_HUMAN | 14-3-3 protein gamma | HUMAN |
| sp\|Q7Z739\|YTHD3_HUMAN | YTH domain-containing family protein 3 | HUMAN |
| sp\|P31689\|DNJA1_HUMAN | DnaJ homolog subfamily A member 1 | HUMAN |
| sp\|Q9BUJ2\|HNRL1_HUMAN | Heterogeneous nuclear ribonucleoprotein U-like protein 1 | HUMAN |
| sp\|Q15907\|RB11B_HUMAN | Ras-related protein Rab-11B | HUMAN |
| sp\|Q9BXS5\|AP1M1_HUMAN | AP-1 complex subunit mu-1 | HUMAN |
| sp\|Q16181\|SEPT7_HUMAN | Septin-7 | HUMAN |
| sp\|Q13409\|DC1I2_HUMAN | Cytoplasmic dynein 1 intermediate chain 2 | HUMAN |
| sp\|P26196\|DDX6_HUMAN | Probable ATP-dependent RNA helicase DDX6 | HUMAN |
| sp\|O00231\|PSD11_HUMAN | 26S proteasome non-ATPase regulatory subunit 11 | HUMAN |
| sp\|P46782\|RS5_HUMAN | 40S ribosomal protein S5 | HUMAN |
| sp\|P48681\|NEST_HUMAN | Nestin | HUMAN |
| sp\|Q9UN86\|G3BP2_HUMAN | Ras GTPase-activating protein-binding protein 2 | HUMAN |
| sp\|Q5JWF2\|GNAS1_HUMAN | Guanine nucleotide-binding protein G(s) subunit alpha isoforms XLas | HUMAN |
| sp\|Q07666\|KHDR1_HUMAN | KH domain-containing, RNA-binding, signal transduction-associated protein 1 | HUMAN |
| sp\|O60506\|HNRPQ_HUMAN | Heterogeneous nuclear ribonucleoprotein Q | HUMAN |
| sp\|Q9NPH2\|INO1_HUMAN | Inositol-3-phosphate synthase 1 | HUMAN |
| sp\|P61019\|RAB2A_HUMAN | Ras-related protein Rab-2A | HUMAN |
| sp\|P09455\|RET1_HUMAN | Retinol-binding protein 1 | HUMAN |
| sp\|Q15019\|SEPT2_HUMAN | Septin-2 | HUMAN |
| sp\|P50991\|TCPD_HUMAN | T-complex protein 1 subunit delta | HUMAN |
| sp\|P62269\|RS18_HUMAN | 40S ribosomal protein S18 | HUMAN |
| sp\|P62873\|GBB1_HUMAN | Guanine nucleotide-binding protein G(I)/G(S)/G(T) subunit beta-1 | HUMAN |
| sp\|P62136\|PP1A_HUMAN | Serine/threonine-protein phosphatase PP1-alpha catalytic subunit | HUMAN |

| | | |
|---|---|---|
| sp\|P27797\|CALR_HUMAN | Calreticulin | HUMAN |
| sp\|O75976\|CBPD_HUMAN | Carboxypeptidase D | HUMAN |
| sp\|Q15084\|PDIA6_HUMAN | Protein disulfide-isomerase A6 | HUMAN |
| sp\|P14314\|GLU2B_HUMAN | Glucosidase 2 subunit beta | HUMAN |
| sp\|O43242\|PSMD3_HUMAN | 26S proteasome non-ATPase regulatory subunit 3 | HUMAN |
| sp\|Q9NR31\|SAR1A_HUMAN | GTP-binding protein SAR1a | HUMAN |
| sp\|Q9UI15\|TAGL3_HUMAN | Transgelin-3 | HUMAN |
| sp\|Q15717\|ELAV1_HUMAN | ELAV-like protein 1 | HUMAN |
| sp\|P61026\|RAB10_HUMAN | Ras-related protein Rab-10 | HUMAN |
| sp\|Q9Y224\|RTRAF_HUMAN | RNA transcription, translation and transport factor protein | HUMAN |
| sp\|P46783\|RS10_HUMAN | 40S ribosomal protein S10 | HUMAN |
| sp\|P21579\|SYT1_HUMAN | Synaptotagmin-1 | HUMAN |
| sp\|O60664\|PLIN3_HUMAN | Perilipin-3 | HUMAN |
| sp\|P18085\|ARF4_HUMAN | ADP-ribosylation factor 4 | HUMAN |
| sp\|P18669\|PGAM1_HUMAN | Phosphoglycerate mutase 1 | HUMAN |
| sp\|P35637\|FUS_HUMAN | RNA-binding protein FUS | HUMAN |
| sp\|P51149\|RAB7A_HUMAN | Ras-related protein Rab-7a | HUMAN |
| sp\|Q92879\|CELF1_HUMAN | CUGBP Elav-like family member 1 | HUMAN |
| sp\|Q01844\|EWS_HUMAN | RNA-binding protein EWS | HUMAN |
| sp\|O75874\|IDHC_HUMAN | Isocitrate dehydrogenase [NADP] cytoplasmic | HUMAN |
| sp\|P26368\|U2AF2_HUMAN | Splicing factor U2AF 65 kDa subunit | HUMAN |
| sp\|O43684\|BUB3_HUMAN | Mitotic checkpoint protein BUB3 | HUMAN |
| sp\|O60716\|CTND1_HUMAN | Catenin delta-1 | HUMAN |
| sp\|P26373\|RL13_HUMAN | 60S ribosomal protein L13 | HUMAN |
| sp\|Q8N684\|CPSF7_HUMAN | Cleavage and polyadenylation specificity factor subunit 7 | HUMAN |
| sp\|P30086\|PEBP1_HUMAN | Phosphatidylethanolamine-binding protein 1 | HUMAN |
| sp\|Q16630\|CPSF6_HUMAN | Cleavage and polyadenylation specificity factor subunit 6 | HUMAN |
| sp\|Q96A49\|SYAP1_HUMAN | Synapse-associated protein 1 | HUMAN |
| sp\|Q06830\|PRDX1_HUMAN | Peroxiredoxin-1 | HUMAN |
| sp\|P54727\|RD23B_HUMAN | UV excision repair protein RAD23 homolog B | HUMAN |
| sp\|Q96PU8\|QKI_HUMAN | KH domain-containing RNA-binding protein QKI | HUMAN |
| sp\|P09429\|HMGB1_HUMAN | High mobility group protein B1 | HUMAN |
| sp\|Q15436\|SC23A_HUMAN | Protein transport protein Sec23A | HUMAN |
| sp\|Q15181\|IPYR_HUMAN | Inorganic pyrophosphatase | HUMAN |
| sp\|Q9UKM9\|RALY_HUMAN | RNA-binding protein Raly | HUMAN |
| sp\|P20020\|AT2B1_HUMAN | Plasma membrane calcium-transporting ATPase 1 | HUMAN |
| sp\|P50148\|GNAQ_HUMAN | Guanine nucleotide-binding protein G(q) subunit alpha | HUMAN |

| | | |
|---|---|---|
| sp\|P07197\|NFM_HUMAN | Neurofilament medium polypeptide | HUMAN |
| sp\|Q12874\|SF3A3_HUMAN | Splicing factor 3A subunit 3 | HUMAN |
| sp\|P55209\|NP1L1_HUMAN | Nucleosome assembly protein 1-like 1 | HUMAN |
| sp\|P31483\|TIA1_HUMAN | Cytotoxic granule associated RNA binding protein TIA1 | HUMAN |
| sp\|P29966\|MARCS_HUMAN | Myristoylated alanine-rich C-kinase substrate | HUMAN |
| sp\|Q9UHD8\|SEPT9_HUMAN | Septin-9 | HUMAN |
| sp\|O60884\|DNJA2_HUMAN | DnaJ homolog subfamily A member 2 | HUMAN |
| sp\|P22087\|FBRL_HUMAN | rRNA 2'-O-methyltransferase fibrillarin | HUMAN |
| sp\|P40429\|RL13A_HUMAN | 60S ribosomal protein L13a | HUMAN |
| sp\|P31150\|GDIA_HUMAN | Rab GDP dissociation inhibitor alpha | HUMAN |
| sp\|P35998\|PRS7_HUMAN | 26S proteasome regulatory subunit 7 | HUMAN |
| sp\|P07355\|ANXA2_HUMAN | Annexin A2 | HUMAN |
| sp\|Q9NVA2\|SEP11_HUMAN | Septin-11 | HUMAN |
| sp\|Q9BVA1\|TBB2B_HUMAN | Tubulin beta-2B chain | HUMAN |
| sp\|Q13492\|PICAL_HUMAN | Phosphatidylinositol-binding clathrin assembly protein | HUMAN |
| sp\|P54920\|SNAA_HUMAN | Alpha-soluble NSF attachment protein | HUMAN |
| sp\|Q15366\|PCBP2_HUMAN | Poly(rC)-binding protein 2 | HUMAN |
| sp\|P63241\|IF5A1_HUMAN | Eukaryotic translation initiation factor 5A-1 | HUMAN |
| sp\|P07237\|PDIA1_HUMAN | Protein disulfide-isomerase | HUMAN |
| sp\|P60660\|MYL6_HUMAN | Myosin light polypeptide 6 | HUMAN |
| sp\|Q9BY77\|PDIP3_HUMAN | Polymerase delta-interacting protein 3 | HUMAN |
| sp\|P49321\|NASP_HUMAN | Nuclear autoantigenic sperm protein | HUMAN |
| sp\|P40616\|ARL1_HUMAN | ADP-ribosylation factor-like protein 1 | HUMAN |
| sp\|Q13200\|PSMD2_HUMAN | 26S proteasome non-ATPase regulatory subunit 2 | HUMAN |
| sp\|Q9Y2W2\|WBP11_HUMAN | WW domain-binding protein 11 | HUMAN |
| sp\|O15240\|VGF_HUMAN | Neurosecretory protein VGF | HUMAN |
| sp\|P62081\|RS7_HUMAN | 40S ribosomal protein S7 | HUMAN |
| sp\|P39687\|AN32A_HUMAN | Acidic leucine-rich nuclear phosphoprotein 32 family member A | HUMAN |
| sp\|Q9UKV3\|ACINU_HUMAN | Apoptotic chromatin condensation inducer in the nucleus | HUMAN |
| sp\|P04899\|GNAI2_HUMAN | Guanine nucleotide-binding protein G(i) subunit alpha-2 | HUMAN |
| sp\|P20340\|RAB6A_HUMAN | Ras-related protein Rab-6A | HUMAN |
| sp\|Q9BZZ5\|API5_HUMAN | Apoptosis inhibitor 5 | HUMAN |
| sp\|Q13247\|SRSF6_HUMAN | Serine/arginine-rich splicing factor 6 | HUMAN |
| sp\|P32119\|PRDX2_HUMAN | Peroxiredoxin-2 | HUMAN |
| sp\|Q9Y383\|LC7L2_HUMAN | Putative RNA-binding protein Luc7-like 2 | HUMAN |
| sp\|Q5SW79\|CE170_HUMAN | Centrosomal protein of 170 kDa | HUMAN |
| sp\|O96019\|ACL6A_HUMAN | Actin-like protein 6A | HUMAN |

| | | |
|---|---|---|
| sp\|Q99729\|ROAA_HUMAN | Heterogeneous nuclear ribonucleoprotein A/B | HUMAN |
| sp\|P23528\|COF1_HUMAN | Cofilin-1 | HUMAN |
| sp\|Q01081\|U2AF1_HUMAN | Splicing factor U2AF 35 kDa subunit | HUMAN |
| sp\|Q8WWY3\|PRP31_HUMAN | U4/U6 small nuclear ribonucleoprotein Prp31 | HUMAN |
| sp\|Q00688\|FKBP3_HUMAN | Peptidyl-prolyl cis-trans isomerase FKBP3 | HUMAN |
| sp\|P08708\|RS17_HUMAN | 40S ribosomal protein S17 | HUMAN |
| sp\|Q14576\|ELAV3_HUMAN | ELAV-like protein 3 | HUMAN |
| sp\|P61020\|RAB5B_HUMAN | Ras-related protein Rab-5B | HUMAN |
| sp\|P62888\|RL30_HUMAN | 60S ribosomal protein L30 | HUMAN |
| sp\|O75122\|CLAP2_HUMAN | CLIP-associating protein 2 | HUMAN |
| sp\|Q07866\|KLC1_HUMAN | Kinesin light chain 1 | HUMAN |
| sp\|Q5SQI0\|ATAT_HUMAN | Alpha-tubulin N-acetyltransferase 1 | HUMAN |
| sp\|Q14257\|RCN2_HUMAN | Reticulocalbin-2 | HUMAN |
| sp\|Q9NVJ2\|ARL8B_HUMAN | ADP-ribosylation factor-like protein 8B | HUMAN |
| sp\|Q7Z7K6\|CENPV_HUMAN | Centromere protein V | HUMAN |
| sp\|P35613\|BASI_HUMAN | Basigin | HUMAN |
| sp\|P62753\|RS6_HUMAN | 40S ribosomal protein S6 | HUMAN |
| sp\|Q15691\|MARE1_HUMAN | Microtubule-associated protein RP/EB family member 1 | HUMAN |
| sp\|P61764\|STXB1_HUMAN | Syntaxin-binding protein 1 | HUMAN |
| sp\|Q14847\|LASP1_HUMAN | LIM and SH3 domain protein 1 | HUMAN |
| sp\|Q86V81\|THOC4_HUMAN | THO complex subunit 4 | HUMAN |
| sp\|O00151\|PDLI1_HUMAN | PDZ and LIM domain protein 1 | HUMAN |
| sp\|P62280\|RS11_HUMAN | 40S ribosomal protein S11 | HUMAN |
| sp\|P50502\|F10A1_HUMAN | Hsc70-interacting protein | HUMAN |
| sp\|P63000\|RAC1_HUMAN | Ras-related C3 botulinum toxin substrate 1 | HUMAN |
| sp\|P13667\|PDIA4_HUMAN | Protein disulfide-isomerase A4 | HUMAN |
| sp\|O75400\|PR40A_HUMAN | Pre-mRNA-processing factor 40 homolog A | HUMAN |
| sp\|Q92896\|GSLG1_HUMAN | Golgi apparatus protein 1 | HUMAN |
| sp\|P09211\|GSTP1_HUMAN | Glutathione S-transferase P | HUMAN |
| sp\|P60900\|PSA6_HUMAN | Proteasome subunit alpha type-6 | HUMAN |
| sp\|Q13151\|ROA0_HUMAN | Heterogeneous nuclear ribonucleoprotein A0 | HUMAN |
| sp\|P62241\|RS8_HUMAN | 40S ribosomal protein S8 | HUMAN |
| sp\|Q9P0L0\|VAPA_HUMAN | Vesicle-associated membrane protein-associated protein A | HUMAN |
| sp\|P62979\|RS27A_HUMAN | Ubiquitin-40S ribosomal protein S27a | HUMAN |
| sp\|Q8NEV1\|CSK23_HUMAN | Casein kinase II subunit alpha 3 | HUMAN |
| sp\|Q14444\|CAPR1_HUMAN | Caprin-1 | HUMAN |
| sp\|Q15008\|PSMD6_HUMAN | 26S proteasome non-ATPase regulatory subunit 6 | HUMAN |
| sp\|Q14019\|COTL1_HUMAN | Coactosin-like protein | HUMAN |
| sp\|P16870\|CBPE_HUMAN | Carboxypeptidase E | HUMAN |

| | | |
|---|---|---|
| sp\|P30153\|2AAA_HUMAN | Serine/threonine-protein phosphatase 2A 65 kDa regulatory subunit A alpha isoform | HUMAN |
| sp\|P46779\|RL28_HUMAN | 60S ribosomal protein L28 | HUMAN |
| sp\|P62750\|RL23A_HUMAN | 60S ribosomal protein L23a | HUMAN |
| sp\|Q13310\|PABP4_HUMAN | Polyadenylate-binding protein 4 | HUMAN |
| sp\|P52209\|6PGD_HUMAN | 6-phosphogluconate dehydrogenase, decarboxylating | HUMAN |
| sp\|Q86Y82\|STX12_HUMAN | Syntaxin-12 | HUMAN |
| sp\|P62805\|H4_HUMAN | Histone H4 | HUMAN |
| sp\|P21281\|VATB2_HUMAN | V-type proton ATPase subunit B, brain isoform | HUMAN |
| sp\|P09496\|CLCA_HUMAN | Clathrin light chain A | HUMAN |
| sp\|Q9UQB8\|BAIP2_HUMAN | Brain-specific angiogenesis inhibitor 1-associated protein 2 | HUMAN |
| sp\|O43765\|SGTA_HUMAN | Small glutamine-rich tetratricopeptide repeat-containing protein alpha | HUMAN |
| sp\|P52907\|CAZA1_HUMAN | F-actin-capping protein subunit alpha-1 | HUMAN |
| sp\|P07384\|CAN1_HUMAN | Calpain-1 catalytic subunit | HUMAN |
| sp\|Q15428\|SF3A2_HUMAN | Splicing factor 3A subunit 2 | HUMAN |
| sp\|Q13283\|G3BP1_HUMAN | Ras GTPase-activating protein-binding protein 1 | HUMAN |
| sp\|Q52LJ0\|FA98B_HUMAN | Protein FAM98B | HUMAN |
| sp\|P04843\|RPN1_HUMAN | Dolichyl-diphosphooligosaccharide--protein glycosyltransferase subunit 1 | HUMAN |
| sp\|P61160\|ARP2_HUMAN | Actin-related protein 2 | HUMAN |
| sp\|P84103\|SRSF3_HUMAN | Serine/arginine-rich splicing factor 3 | HUMAN |
| sp\|P61254\|RL26_HUMAN | 60S ribosomal protein L26 | HUMAN |
| sp\|P62277\|RS13_HUMAN | 40S ribosomal protein S13 | HUMAN |
| sp\|P32969\|RL9_HUMAN | 60S ribosomal protein L9 | HUMAN |
| sp\|P67936\|TPM4_HUMAN | Tropomyosin alpha-4 chain | HUMAN |
| sp\|O00232\|PSD12_HUMAN | 26S proteasome non-ATPase regulatory subunit 12 | HUMAN |
| sp\|Q7L1Q6\|5MP2_HUMAN | eIF5-mimic protein 2 | HUMAN |
| sp\|P23193\|TCEA1_HUMAN | Transcription elongation factor A protein 1 | HUMAN |
| sp\|P18621\|RL17_HUMAN | 60S ribosomal protein L17 | HUMAN |
| sp\|P61163\|ACTZ_HUMAN | Alpha-centractin | HUMAN |
| sp\|P27635\|RL10_HUMAN | 60S ribosomal protein L10 | HUMAN |
| sp\|P05023\|AT1A1_HUMAN | Sodium/potassium-transporting ATPase subunit alpha-1 | HUMAN |
| sp\|P31350\|RIR2_HUMAN | Ribonucleoside-diphosphate reductase subunit M2 | HUMAN |
| sp\|Q99733\|NP1L4_HUMAN | Nucleosome assembly protein 1-like 4 | HUMAN |
| sp\|P38606\|VATA_HUMAN | V-type proton ATPase catalytic subunit A | HUMAN |
| sp\|P62316\|SMD2_HUMAN | Small nuclear ribonucleoprotein Sm D2 | HUMAN |

| | | |
|---|---|---|
| sp\|P27348\|1433T_HUMAN | 14-3-3 protein theta | HUMAN |
| sp\|O00264\|PGRC1_HUMAN | Membrane-associated progesterone receptor component 1 | HUMAN |
| sp\|O00422\|SAP18_HUMAN | Histone deacetylase complex subunit SAP18 | HUMAN |
| sp\|P62826\|RAN_HUMAN | GTP-binding nuclear protein Ran | HUMAN |
| sp\|P25786\|PSA1_HUMAN | Proteasome subunit alpha type-1 | HUMAN |
| sp\|Q92804\|RBP56_HUMAN | TATA-binding protein-associated factor 2N | HUMAN |
| sp\|P07196\|NFL_HUMAN | Neurofilament light polypeptide | HUMAN |
| sp\|P61586\|RHOA_HUMAN | Transforming protein RhoA | HUMAN |
| sp\|Q53GQ0\|DHB12_HUMAN | Very-long-chain 3-oxoacyl-CoA reductase | HUMAN |
| sp\|Q92930\|RAB8B_HUMAN | Ras-related protein Rab-8B | HUMAN |
| sp\|P62195\|PRS8_HUMAN | 26S proteasome regulatory subunit 8 | HUMAN |
| sp\|O43776\|SYNC_HUMAN | Asparagine--tRNA ligase, cytoplasmic | HUMAN |
| sp\|Q93050\|VPP1_HUMAN | V-type proton ATPase 116 kDa subunit a 1 | HUMAN |
| sp\|P0DP25\|CALM3_HUMAN | Calmodulin-3 | HUMAN |
| sp\|Q99459\|CDC5L_HUMAN | Cell division cycle 5-like protein | HUMAN |
| sp\|Q01518\|CAP1_HUMAN | Adenylyl cyclase-associated protein 1 | HUMAN |
| sp\|P09661\|RU2A_HUMAN | U2 small nuclear ribonucleoprotein A' | HUMAN |
| sp\|P27694\|RFA1_HUMAN | Replication protein A 70 kDa DNA-binding subunit | HUMAN |
| sp\|Q92890\|UFD1_HUMAN | Ubiquitin recognition factor in ER-associated degradation protein 1 | HUMAN |
| sp\|P30041\|PRDX6_HUMAN | Peroxiredoxin-6 | HUMAN |
| sp\|P78310\|CXAR_HUMAN | Coxsackievirus and adenovirus receptor | HUMAN |
| sp\|P78406\|RAE1L_HUMAN | mRNA export factor RAE1 | HUMAN |
| sp\|P20645\|MPRD_HUMAN | Cation-dependent mannose-6-phosphate receptor | HUMAN |
| sp\|P04632\|CPNS1_HUMAN | Calpain small subunit 1 | HUMAN |
| sp\|P07737\|PROF1_HUMAN | Profilin-1 | HUMAN |
| sp\|Q9HB71\|CYBP_HUMAN | Calcyclin-binding protein | HUMAN |
| sp\|P62633\|CNBP_HUMAN | CCHC-type zinc finger nucleic acid binding protein | HUMAN |
| sp\|P05026\|AT1B1_HUMAN | Sodium/potassium-transporting ATPase subunit beta-1 | HUMAN |
| sp\|P15531\|NDKA_HUMAN | Nucleoside diphosphate kinase A | HUMAN |
| sp\|P25398\|RS12_HUMAN | 40S ribosomal protein S12 | HUMAN |
| sp\|P19105\|ML12A_HUMAN | Myosin regulatory light chain 12A | HUMAN |
| sp\|P28074\|PSB5_HUMAN | Proteasome subunit beta type-5 | HUMAN |
| sp\|Q7Z5L9\|I2BP2_HUMAN | Interferon regulatory factor 2-binding protein 2 | HUMAN |
| sp\|O14818\|PSA7_HUMAN | Proteasome subunit alpha type-7 | HUMAN |
| sp\|P61088\|UBE2N_HUMAN | Ubiquitin-conjugating enzyme E2 N | HUMAN |
| sp\|Q6Y7W6\|GGYF2_HUMAN | GRB10-interacting GYF protein 2 | HUMAN |
| sp\|P27695\|APEX1_HUMAN | DNA-(apurinic or apyrimidinic site) endonuclease | HUMAN |

| | | |
|---|---|---|
| sp\|Q96E17\|RAB3C_HUMAN | Ras-related protein Rab-3C | HUMAN |
| sp\|Q15424\|SAFB1_HUMAN | Scaffold attachment factor B1 | HUMAN |
| sp\|P51513\|NOVA1_HUMAN | RNA-binding protein Nova-1 | HUMAN |
| sp\|O15498\|YKT6_HUMAN | Synaptobrevin homolog YKT6 | HUMAN |
| sp\|P05198\|IF2A_HUMAN | Eukaryotic translation initiation factor 2 subunit 1 | HUMAN |
| sp\|Q16527\|CSRP2_HUMAN | Cysteine and glycine-rich protein 2 | HUMAN |
| sp\|P28066\|PSA5_HUMAN | Proteasome subunit alpha type-5 | HUMAN |
| sp\|Q9ULK5\|VANG2_HUMAN | Vang-like protein 2 | HUMAN |
| sp\|Q99497\|PARK7_HUMAN | Parkinson disease protein 7 | HUMAN |
| sp\|P07339\|CATD_HUMAN | Cathepsin D | HUMAN |
| sp\|O14828\|SCAM3_HUMAN | Secretory carrier-associated membrane protein 3 | HUMAN |
| sp\|Q7L0J3\|SV2A_HUMAN | Synaptic vesicle glycoprotein 2A | HUMAN |
| sp\|P25788\|PSA3_HUMAN | Proteasome subunit alpha type-3 | HUMAN |
| sp\|P35813\|PPM1A_HUMAN | Protein phosphatase 1A | HUMAN |
| sp\|O95793\|STAU1_HUMAN | Double-stranded RNA-binding protein Staufen homolog 1 | HUMAN |
| sp\|O15400\|STX7_HUMAN | Syntaxin-7 | HUMAN |
| sp\|P01111\|RASN_HUMAN | GTPase NRas | HUMAN |
| sp\|Q9Y5A9\|YTHD2_HUMAN | YTH domain-containing family protein 2 | HUMAN |
| sp\|P04350\|TBB4A_HUMAN | Tubulin beta-4A chain | HUMAN |
| sp\|O14979\|HNRDL_HUMAN | Heterogeneous nuclear ribonucleoprotein D-like | HUMAN |
| sp\|P62841\|RS15_HUMAN | 40S ribosomal protein S15 | HUMAN |
| sp\|Q07020\|RL18_HUMAN | 60S ribosomal protein L18 | HUMAN |
| sp\|P17980\|PRS6A_HUMAN | 26S proteasome regulatory subunit 6A | HUMAN |
| sp\|P62906\|RL10A_HUMAN | 60S ribosomal protein L10a | HUMAN |
| sp\|P46778\|RL21_HUMAN | 60S ribosomal protein L21 | HUMAN |
| sp\|O43143\|DHX15_HUMAN | ATP-dependent RNA helicase DHX15 | HUMAN |
| sp\|P21796\|VDAC1_HUMAN | Voltage-dependent anion-selective channel protein 1 | HUMAN |
| sp\|P82979\|SARNP_HUMAN | SAP domain-containing ribonucleoprotein | HUMAN |
| sp\|Q16629\|SRSF7_HUMAN | Serine/arginine-rich splicing factor 7 | HUMAN |
| sp\|P09497\|CLCB_HUMAN | Clathrin light chain B | HUMAN |
| sp\|Q99439\|CNN2_HUMAN | Calponin-2 | HUMAN |
| sp\|O15075\|DCLK1_HUMAN | Serine/threonine-protein kinase DCLK1 | HUMAN |
| sp\|P61313\|RL15_HUMAN | 60S ribosomal protein L15 | HUMAN |
| sp\|P29373\|RABP2_HUMAN | Cellular retinoic acid-binding protein 2 | HUMAN |
| sp\|P55795\|HNRH2_HUMAN | Heterogeneous nuclear ribonucleoprotein H2 | HUMAN |
| sp\|P43686\|PRS6B_HUMAN | 26S proteasome regulatory subunit 6B | HUMAN |
| sp\|Q96M27\|PRRC1_HUMAN | Protein PRRC1 | HUMAN |
| sp\|P30050\|RL12_HUMAN | 60S ribosomal protein L12 | HUMAN |

| | | |
|---|---|---|
| sp\|O94979\|SC31A_HUMAN | Protein transport protein Sec31A | HUMAN |
| sp\|P67870\|CSK2B_HUMAN | Casein kinase II subunit beta | HUMAN |
| sp\|P15311\|EZRI_HUMAN | Ezrin | HUMAN |
| sp\|P26992\|CNTFR_HUMAN | Ciliary neurotrophic factor receptor subunit alpha | HUMAN |
| sp\|Q7L099\|RUFY3_HUMAN | Protein RUFY3 | HUMAN |
| sp\|Q92522\|H1X_HUMAN | Histone H1.10 | HUMAN |
| sp\|P98175\|RBM10_HUMAN | RNA-binding protein 10 | HUMAN |
| sp\|Q02543\|RL18A_HUMAN | 60S ribosomal protein L18a | HUMAN |
| sp\|P32004\|L1CAM_HUMAN | Neural cell adhesion molecule L1 | HUMAN |
| sp\|Q9H444\|CHM4B_HUMAN | Charged multivesicular body protein 4b | HUMAN |
| sp\|Q99523\|SORT_HUMAN | Sortilin | HUMAN |
| sp\|O76094\|SRP72_HUMAN | Signal recognition particle subunit SRP72 | HUMAN |
| sp\|Q15185\|TEBP_HUMAN | Prostaglandin E synthase 3 | HUMAN |
| sp\|Q04917\|1433F_HUMAN | 14-3-3 protein eta | HUMAN |
| sp\|P47756\|CAPZB_HUMAN | F-actin-capping protein subunit beta | HUMAN |
| sp\|Q01105\|SET_HUMAN | Protein SET | HUMAN |
| sp\|Q6PKG0\|LARP1_HUMAN | La-related protein 1 | HUMAN |
| sp\|P55036\|PSMD4_HUMAN | 26S proteasome non-ATPase regulatory subunit 4 | HUMAN |
| sp\|Q14011\|CIRBP_HUMAN | Cold-inducible RNA-binding protein | HUMAN |
| sp\|P62263\|RS14_HUMAN | 40S ribosomal protein S14 | HUMAN |
| sp\|Q13242\|SRSF9_HUMAN | Serine/arginine-rich splicing factor 9 | HUMAN |
| sp\|P62851\|RS25_HUMAN | 40S ribosomal protein S25 | HUMAN |
| sp\|P11166\|GTR1_HUMAN | Solute carrier family 2, facilitated glucose transporter member 1 | HUMAN |
| sp\|P49207\|RL34_HUMAN | 60S ribosomal protein L34 | HUMAN |
| sp\|P49721\|PSB2_HUMAN | Proteasome subunit beta type-2 | HUMAN |
| sp\|P83731\|RL24_HUMAN | 60S ribosomal protein L24 | HUMAN |
| sp\|P20290\|BTF3_HUMAN | Transcription factor BTF3 | HUMAN |
| sp\|P52594\|AGFG1_HUMAN | Arf-GAP domain and FG repeat-containing protein 1 | HUMAN |
| sp\|P37840\|SYUA_HUMAN | Alpha-synuclein | HUMAN |
| sp\|Q9NYF8\|BCLF1_HUMAN | Bcl-2-associated transcription factor 1 | HUMAN |
| sp\|O15126\|SCAM1_HUMAN | Secretory carrier-associated membrane protein 1 | HUMAN |
| sp\|O60573\|IF4E2_HUMAN | Eukaryotic translation initiation factor 4E type 2 | HUMAN |
| sp\|Q9NP72\|RAB18_HUMAN | Ras-related protein Rab-18 | HUMAN |
| sp\|P08758\|ANXA5_HUMAN | Annexin A5 | HUMAN |
| sp\|P25789\|PSA4_HUMAN | Proteasome subunit alpha type-4 | HUMAN |
| sp\|Q9HDC9\|APMAP_HUMAN | Adipocyte plasma membrane-associated protein | HUMAN |
| sp\|P09012\|SNRPA_HUMAN | U1 small nuclear ribonucleoprotein A | HUMAN |

| | | |
|---|---|---|
| sp\|Q9BWD1\|THIC_HUMAN | Acetyl-CoA acetyltransferase, cytosolic | HUMAN |
| sp\|P62244\|RS15A_HUMAN | 40S ribosomal protein S15a | HUMAN |
| sp\|P53999\|TCP4_HUMAN | Activated RNA polymerase II transcriptional coactivator p15 | HUMAN |
| sp\|P50914\|RL14_HUMAN | 60S ribosomal protein L14 | HUMAN |
| sp\|P16615\|AT2A2_HUMAN | Sarcoplasmic/endoplasmic reticulum calcium ATPase 2 | HUMAN |
| sp\|P45973\|CBX5_HUMAN | Chromobox protein homolog 5 | HUMAN |
| sp\|P62829\|RL23_HUMAN | 60S ribosomal protein L23 | HUMAN |
| sp\|Q9P035\|HACD3_HUMAN | Very-long-chain (3R)-3-hydroxyacyl-CoA dehydratase 3 | HUMAN |
| sp\|P10253\|LYAG_HUMAN | Lysosomal alpha-glucosidase | HUMAN |
| sp\|P61224\|RAP1B_HUMAN | Ras-related protein Rap-1b | HUMAN |
| sp\|O75396\|SC22B_HUMAN | Vesicle-trafficking protein SEC22b | HUMAN |
| sp\|P28070\|PSB4_HUMAN | Proteasome subunit beta type-4 | HUMAN |
| sp\|Q9Y6B6\|SAR1B_HUMAN | GTP-binding protein SAR1b | HUMAN |
| sp\|P84085\|ARF5_HUMAN | ADP-ribosylation factor 5 | HUMAN |
| sp\|Q01469\|FABP5_HUMAN | Fatty acid-binding protein 5 | HUMAN |
| sp\|P49419\|AL7A1_HUMAN | Alpha-aminoadipic semialdehyde dehydrogenase | HUMAN |
| sp\|P54709\|AT1B3_HUMAN | Sodium/potassium-transporting ATPase subunit beta-3 | HUMAN |
| sp\|Q9UL25\|RAB21_HUMAN | Ras-related protein Rab-21 | HUMAN |
| sp\|Q96HC4\|PDLI5_HUMAN | PDZ and LIM domain protein 5 | HUMAN |
| sp\|O95292\|VAPB_HUMAN | Vesicle-associated membrane protein-associated protein B/C | HUMAN |
| sp\|P39656\|OST48_HUMAN | Dolichyl-diphosphooligosaccharide--protein glycosyltransferase 48 kDa subunit | HUMAN |
| sp\|Q13642\|FHL1_HUMAN | Four and a half LIM domains protein 1 | HUMAN |
| sp\|P11216\|PYGB_HUMAN | Glycogen phosphorylase, brain form | HUMAN |
| sp\|P08195\|4F2_HUMAN | 4F2 cell-surface antigen heavy chain | HUMAN |
| sp\|P61353\|RL27_HUMAN | 60S ribosomal protein L27 | HUMAN |
| sp\|P09104\|ENOG_HUMAN | Gamma-enolase | HUMAN |
| sp\|Q8IYB3\|SRRM1_HUMAN | Serine/arginine repetitive matrix protein 1 | HUMAN |
| sp\|P11586\|C1TC_HUMAN | C-1-tetrahydrofolate synthase, cytoplasmic | HUMAN |
| sp\|P16401\|H15_HUMAN | Histone H1.5 | HUMAN |
| sp\|Q92530\|PSMF1_HUMAN | Proteasome inhibitor PI31 subunit | HUMAN |
| sp\|P36543\|VATE1_HUMAN | V-type proton ATPase subunit E 1 | HUMAN |
| sp\|E9PAV3\|NACAM_HUMAN | Nascent polypeptide-associated complex subunit alpha, muscle-specific form | HUMAN |
| sp\|P06454\|PTMA_HUMAN | Prothymosin alpha | HUMAN |
| sp\|P45974\|UBP5_HUMAN | Ubiquitin carboxyl-terminal hydrolase 5 | HUMAN |
| sp\|O43633\|CHM2A_HUMAN | Charged multivesicular body protein 2a | HUMAN |
| sp\|Q96DH6\|MSI2H_HUMAN | RNA-binding protein Musashi homolog 2 | HUMAN |

| | | |
|---|---|---|
| sp\|Q9Y3U8\|RL36_HUMAN | 60S ribosomal protein L36 | HUMAN |
| sp\|P19388\|RPAB1_HUMAN | DNA-directed RNA polymerases I, II, and III subunit RPABC1 | HUMAN |
| sp\|P54619\|AAKG1_HUMAN | 5'-AMP-activated protein kinase subunit gamma-1 | HUMAN |
| sp\|Q16352\|AINX_HUMAN | Alpha-internexin | HUMAN |
| sp\|P37108\|SRP14_HUMAN | Signal recognition particle 14 kDa protein | HUMAN |
| sp\|P35241\|RADI_HUMAN | Radixin | HUMAN |
| sp\|P20339\|RAB5A_HUMAN | Ras-related protein Rab-5A | HUMAN |
| sp\|P60953\|CDC42_HUMAN | Cell division control protein 42 homolog | HUMAN |
| sp\|P57723\|PCBP4_HUMAN | Poly(rC)-binding protein 4 | HUMAN |
| sp\|P62330\|ARF6_HUMAN | ADP-ribosylation factor 6 | HUMAN |
| sp\|Q14108\|SCRB2_HUMAN | Lysosome membrane protein 2 | HUMAN |
| sp\|P62314\|SMD1_HUMAN | Small nuclear ribonucleoprotein Sm D1 | HUMAN |
| sp\|P11233\|RALA_HUMAN | Ras-related protein Ral-A | HUMAN |
| sp\|P00387\|NB5R3_HUMAN | NADH-cytochrome b5 reductase 3 | HUMAN |
| sp\|Q96GY0\|ZC21A_HUMAN | Zinc finger C2HC domain-containing protein 1A | HUMAN |
| sp\|P55884\|EIF3B_HUMAN | Eukaryotic translation initiation factor 3 subunit B | HUMAN |
| sp\|P05556\|ITB1_HUMAN | Integrin beta-1 | HUMAN |
| sp\|P62917\|RL8_HUMAN | 60S ribosomal protein L8 | HUMAN |
| sp\|P62266\|RS23_HUMAN | 40S ribosomal protein S23 | HUMAN |
| sp\|Q01130\|SRSF2_HUMAN | Serine/arginine-rich splicing factor 2 | HUMAN |
| sp\|Q8N6T3\|ARFG1_HUMAN | ADP-ribosylation factor GTPase-activating protein 1 | HUMAN |
| sp\|Q02246\|CNTN2_HUMAN | Contactin-2 | HUMAN |
| sp\|P18077\|RL35A_HUMAN | 60S ribosomal protein L35a | HUMAN |
| sp\|Q8IV08\|PLD3_HUMAN | 5'-3' exonuclease PLD3 | HUMAN |
| sp\|O95865\|DDAH2_HUMAN | N(G),N(G)-dimethylarginine dimethylaminohydrolase 2 | HUMAN |
| sp\|Q96KP4\|CNDP2_HUMAN | Cytosolic non-specific dipeptidase | HUMAN |
| sp\|Q7Z4W1\|DCXR_HUMAN | L-xylulose reductase | HUMAN |
| sp\|Q15056\|IF4H_HUMAN | Eukaryotic translation initiation factor 4H | HUMAN |
| sp\|Q99747\|SNAG_HUMAN | Gamma-soluble NSF attachment protein | HUMAN |
| sp\|Q86X55\|CARM1_HUMAN | Histone-arginine methyltransferase CARM1 | HUMAN |
| sp\|O75367\|H2AY_HUMAN | Core histone macro-H2A.1 | HUMAN |
| sp\|P18031\|PTN1_HUMAN | Tyrosine-protein phosphatase non-receptor type 1 | HUMAN |
| sp\|Q13526\|PIN1_HUMAN | Peptidyl-prolyl cis-trans isomerase NIMA-interacting 1 | HUMAN |
| sp\|Q7RTV0\|PHF5A_HUMAN | PHD finger-like domain-containing protein 5A | HUMAN |
| sp\|P10599\|THIO_HUMAN | Thioredoxin | HUMAN |
| sp\|P06493\|CDK1_HUMAN | Cyclin-dependent kinase 1 | HUMAN |

| | | |
|---|---|---|
| sp\|P35579\|MYH9_HUMAN | Myosin-9 | HUMAN |
| sp\|Q9H115\|SNAB_HUMAN | Beta-soluble NSF attachment protein | HUMAN |
| sp\|Q9Y4L1\|HYOU1_HUMAN | Hypoxia up-regulated protein 1 | HUMAN |
| sp\|P60880\|SNP25_HUMAN | Synaptosomal-associated protein 25 | HUMAN |
| sp\|Q15363\|TMED2_HUMAN | Transmembrane emp24 domain-containing protein 2 | HUMAN |
| sp\|P06576\|ATPB_HUMAN | ATP synthase subunit beta, mitochondrial | HUMAN |
| sp\|Q9BRX8\|PXL2A_HUMAN | Peroxiredoxin-like 2A | HUMAN |
| sp\|O94973\|AP2A2_HUMAN | AP-2 complex subunit alpha-2 | HUMAN |
| sp\|Q8WW12\|PCNP_HUMAN | PEST proteolytic signal-containing nuclear protein | HUMAN |
| sp\|Q9Y371\|SHLB1_HUMAN | Endophilin-B1 | HUMAN |
| sp\|Q15942\|ZYX_HUMAN | Zyxin | HUMAN |
| sp\|P62899\|RL31_HUMAN | 60S ribosomal protein L31 | HUMAN |
| sp\|O43399\|TPD54_HUMAN | Tumor protein D54 | HUMAN |
| sp\|Q9NWB1\|RFOX1_HUMAN | RNA binding protein fox-1 homolog 1 | HUMAN |
| sp\|P62847\|RS24_HUMAN | 40S ribosomal protein S24 | HUMAN |
| sp\|Q96BM9\|ARL8A_HUMAN | ADP-ribosylation factor-like protein 8A | HUMAN |
| sp\|Q8N8S7\|ENAH_HUMAN | Protein enabled homolog | HUMAN |
| sp\|P46776\|RL27A_HUMAN | 60S ribosomal protein L27a | HUMAN |
| sp\|Q9H4M9\|EHD1_HUMAN | EH domain-containing protein 1 | HUMAN |
| sp\|P49755\|TMEDA_HUMAN | Transmembrane emp24 domain-containing protein 10 | HUMAN |
| sp\|P51572\|BAP31_HUMAN | B-cell receptor-associated protein 31 | HUMAN |
| sp\|P09936\|UCHL1_HUMAN | Ubiquitin carboxyl-terminal hydrolase isozyme L1 | HUMAN |
| sp\|P00338\|LDHA_HUMAN | L-lactate dehydrogenase A chain | HUMAN |
| sp\|P49257\|LMAN1_HUMAN | Protein ERGIC-53 | HUMAN |
| sp\|Q9H8Y8\|GORS2_HUMAN | Golgi reassembly-stacking protein 2 | HUMAN |
| sp\|Q8N9N7\|LRC57_HUMAN | Leucine-rich repeat-containing protein 57 | HUMAN |
| sp\|Q9Y266\|NUDC_HUMAN | Nuclear migration protein nudC | HUMAN |
| sp\|P31946\|1433B_HUMAN | 14-3-3 protein beta/alpha | HUMAN |
| sp\|Q99816\|TS101_HUMAN | Tumor susceptibility gene 101 protein | HUMAN |
| sp\|P84077\|ARF1_HUMAN | ADP-ribosylation factor 1 | HUMAN |
| sp\|P61158\|ARP3_HUMAN | Actin-related protein 3 | HUMAN |
| sp\|P09543\|CN37_HUMAN | 2',3'-cyclic-nucleotide 3'-phosphodiesterase | HUMAN |
| sp\|P30040\|ERP29_HUMAN | Endoplasmic reticulum resident protein 29 | HUMAN |
| sp\|P25787\|PSA2_HUMAN | Proteasome subunit alpha type-2 | HUMAN |
| sp\|P25705\|ATPA_HUMAN | ATP synthase subunit alpha, mitochondrial | HUMAN |
| sp\|P63173\|RL38_HUMAN | 60S ribosomal protein L38 | HUMAN |
| sp\|Q9BUF5\|TBB6_HUMAN | Tubulin beta-6 chain | HUMAN |
| sp\|Q15392\|DHC24_HUMAN | Delta(24)-sterol reductase | HUMAN |
| sp\|P05387\|RLA2_HUMAN | 60S acidic ribosomal protein P2 | HUMAN |

| | | |
|---|---|---|
| sp\|P62166\|NCS1_HUMAN | Neuronal calcium sensor 1 | HUMAN |
| sp\|Q13185\|CBX3_HUMAN | Chromobox protein homolog 3 | HUMAN |
| sp\|Q16543\|CDC37_HUMAN | Hsp90 co-chaperone Cdc37 | HUMAN |
| sp\|Q96I24\|FUBP3_HUMAN | Far upstream element-binding protein 3 | HUMAN |
| sp\|P51148\|RAB5C_HUMAN | Ras-related protein Rab-5C | HUMAN |
| sp\|P49458\|SRP09_HUMAN | Signal recognition particle 9 kDa protein | HUMAN |
| sp\|Q9C040\|TRIM2_HUMAN | Tripartite motif-containing protein 2 | HUMAN |
| sp\|P62854\|RS26_HUMAN | 40S ribosomal protein S26 | HUMAN |
| sp\|Q96A72\|MGN2_HUMAN | Protein mago nashi homolog 2 | HUMAN |
| sp\|Q16799\|RTN1_HUMAN | Reticulon-1 | HUMAN |
| sp\|P55327\|TPD52_HUMAN | Tumor protein D52 | HUMAN |
| sp\|P43487\|RANG_HUMAN | Ran-specific GTPase-activating protein | HUMAN |
| sp\|P08579\|RU2B_HUMAN | U2 small nuclear ribonucleoprotein B" | HUMAN |
| sp\|Q9Y5S9\|RBM8A_HUMAN | RNA-binding protein 8A | HUMAN |
| sp\|P53990\|IST1_HUMAN | IST1 homolog | HUMAN |
| sp\|Q96K17\|BT3L4_HUMAN | Transcription factor BTF3 homolog 4 | HUMAN |
| sp\|P60866\|RS20_HUMAN | 40S ribosomal protein S20 | HUMAN |
| sp\|P61513\|RL37A_HUMAN | 60S ribosomal protein L37a | HUMAN |
| sp\|Q9Y3D6\|FIS1_HUMAN | Mitochondrial fission 1 protein | HUMAN |
| sp\|P62995\|TRA2B_HUMAN | Transformer-2 protein homolog beta | HUMAN |
| sp\|P62333\|PRS10_HUMAN | 26S proteasome regulatory subunit 10B | HUMAN |
| sp\|P07305\|H10_HUMAN | Histone H1.0 | HUMAN |
| sp\|Q9Y3Y2\|CHTOP_HUMAN | Chromatin target of PRMT1 protein | HUMAN |
| sp\|P16435\|NCPR_HUMAN | NADPH--cytochrome P450 reductase | HUMAN |
| sp\|O00299\|CLIC1_HUMAN | Chloride intracellular channel protein 1 | HUMAN |
| sp\|Q9NUL3\|STAU2_HUMAN | Double-stranded RNA-binding protein Staufen homolog 2 | HUMAN |
| sp\|P59998\|ARPC4_HUMAN | Actin-related protein 2/3 complex subunit 4 | HUMAN |
| sp\|P20618\|PSB1_HUMAN | Proteasome subunit beta type-1 | HUMAN |
| sp\|P84098\|RL19_HUMAN | 60S ribosomal protein L19 | HUMAN |
| sp\|Q99426\|TBCB_HUMAN | Tubulin-folding cofactor B | HUMAN |
| sp\|O43670\|ZN207_HUMAN | BUB3-interacting and GLEBS motif-containing protein ZNF207 | HUMAN |
| sp\|Q71UM5\|RS27L_HUMAN | 40S ribosomal protein S27-like | HUMAN |
| sp\|P08247\|SYPH_HUMAN | Synaptophysin | HUMAN |
| sp\|P62910\|RL32_HUMAN | 60S ribosomal protein L32 | HUMAN |
| sp\|Q9UNZ2\|NSF1C_HUMAN | NSFL1 cofactor p47 | HUMAN |
| sp\|P53680\|AP2S1_HUMAN | AP-2 complex subunit sigma | HUMAN |
| sp\|Q16186\|ADRM1_HUMAN | Proteasomal ubiquitin receptor ADRM1 | HUMAN |
| sp\|Q92688\|AN32B_HUMAN | Acidic leucine-rich nuclear phosphoprotein 32 family member B | HUMAN |
| sp\|Q07812\|BAX_HUMAN | Apoptosis regulator BAX | HUMAN |
| sp\|O15144\|ARPC2_HUMAN | Actin-related protein 2/3 complex subunit 2 | HUMAN |

| | | |
|---|---|---|
| sp\|P49006\|MRP_HUMAN | MARCKS-related protein | HUMAN |
| sp\|Q71DI3\|H32_HUMAN | Histone H3.2 | HUMAN |
| sp\|P56373\|P2RX3_HUMAN | P2X purinoceptor 3 | HUMAN |
| sp\|P24534\|EF1B_HUMAN | Elongation factor 1-beta | HUMAN |
| sp\|O75569\|PRKRA_HUMAN | Interferon-inducible double-stranded RNA-dependent protein kinase activator A | HUMAN |
| sp\|P62306\|RUXF_HUMAN | Small nuclear ribonucleoprotein F | HUMAN |
| sp\|P98164\|LRP2_HUMAN | Low-density lipoprotein receptor-related protein 2 | HUMAN |
| sp\|O75525\|KHDR3_HUMAN | KH domain-containing, RNA-binding, signal transduction-associated protein 3 | HUMAN |
| sp\|Q9UBC2\|EP15R_HUMAN | Epidermal growth factor receptor substrate 15-like 1 | HUMAN |
| sp\|Q8IWA5\|CTL2_HUMAN | Choline transporter-like protein 2 | HUMAN |
| sp\|P40926\|MDHM_HUMAN | Malate dehydrogenase, mitochondrial | HUMAN |
| sp\|P48556\|PSMD8_HUMAN | 26S proteasome non-ATPase regulatory subunit 8 | HUMAN |
| sp\|P06756\|ITAV_HUMAN | Integrin alpha-V | HUMAN |
| sp\|P37837\|TALDO_HUMAN | Transaldolase | HUMAN |
| sp\|P10636\|TAU_HUMAN | Microtubule-associated protein tau | HUMAN |
| sp\|O43447\|PPIH_HUMAN | Peptidyl-prolyl cis-trans isomerase H | HUMAN |
| sp\|P62913\|RL11_HUMAN | 60S ribosomal protein L11 | HUMAN |
| sp\|Q05519\|SRS11_HUMAN | Serine/arginine-rich splicing factor 11 | HUMAN |
| sp\|Q15286\|RAB35_HUMAN | Ras-related protein Rab-35 | HUMAN |
| sp\|P53985\|MOT1_HUMAN | Monocarboxylate transporter 1 | HUMAN |
| sp\|P49720\|PSB3_HUMAN | Proteasome subunit beta type-3 | HUMAN |
| sp\|P61601\|NCALD_HUMAN | Neurocalcin-delta | HUMAN |
| sp\|P51665\|PSMD7_HUMAN | 26S proteasome non-ATPase regulatory subunit 7 | HUMAN |
| sp\|P29692\|EF1D_HUMAN | Elongation factor 1-delta | HUMAN |
| sp\|Q13243\|SRSF5_HUMAN | Serine/arginine-rich splicing factor 5 | HUMAN |
| sp\|O00487\|PSDE_HUMAN | 26S proteasome non-ATPase regulatory subunit 14 | HUMAN |
| sp\|O14662\|STX16_HUMAN | Syntaxin-16 | HUMAN |
| sp\|O95747\|OXSR1_HUMAN | Serine/threonine-protein kinase OSR1 | HUMAN |
| sp\|Q9UHD9\|UBQL2_HUMAN | Ubiquilin-2 | HUMAN |
| sp\|P55735\|SEC13_HUMAN | Protein SEC13 homolog | HUMAN |
| sp\|Q86U42\|PABP2_HUMAN | Polyadenylate-binding protein 2 | HUMAN |
| sp\|Q9NR46\|SHLB2_HUMAN | Endophilin-B2 | HUMAN |
| sp\|O15347\|HMGB3_HUMAN | High mobility group protein B3 | HUMAN |
| sp\|Q02818\|NUCB1_HUMAN | Nucleobindin-1 | HUMAN |
| sp\|A1L0T0\|HACL2_HUMAN | 2-hydroxyacyl-CoA lyase 2 | HUMAN |
| sp\|O43169\|CYB5B_HUMAN | Cytochrome b5 type B | HUMAN |
| sp\|Q12907\|LMAN2_HUMAN | Vesicular integral-membrane protein VIP36 | HUMAN |

| | | |
|---|---|---|
| sp\|Q14978\|NOLC1_HUMAN | Nucleolar and coiled-body phosphoprotein 1 | HUMAN |
| sp\|P26378\|ELAV4_HUMAN | ELAV-like protein 4 | HUMAN |
| sp\|Q14240\|IF4A2_HUMAN | Eukaryotic initiation factor 4A-II | HUMAN |
| sp\|Q9BYJ9\|YTHD1_HUMAN | YTH domain-containing family protein 1 | HUMAN |
| sp\|Q93045\|STMN2_HUMAN | Stathmin-2 | HUMAN |
| sp\|O15371\|EIF3D_HUMAN | Eukaryotic translation initiation factor 3 subunit D | HUMAN |
| sp\|Q9Y3C6\|PPIL1_HUMAN | Peptidyl-prolyl cis-trans isomerase-like 1 | HUMAN |
| sp\|P63208\|SKP1_HUMAN | S-phase kinase-associated protein 1 | HUMAN |
| sp\|P05204\|HMGN2_HUMAN | Non-histone chromosomal protein HMG-17 | HUMAN |
| sp\|Q6IAA8\|LTOR1_HUMAN | Ragulator complex protein LAMTOR1 | HUMAN |
| sp\|Q15738\|NSDHL_HUMAN | Sterol-4-alpha-carboxylate 3-dehydrogenase, decarboxylating | HUMAN |
| sp\|P62273\|RS29_HUMAN | 40S ribosomal protein S29 | HUMAN |
| sp\|P09234\|RU1C_HUMAN | U1 small nuclear ribonucleoprotein C | HUMAN |
| sp\|Q12800\|TFCP2_HUMAN | Alpha-globin transcription factor CP2 | HUMAN |
| sp\|P62857\|RS28_HUMAN | 40S ribosomal protein S28 | HUMAN |
| sp\|P21291\|CSRP1_HUMAN | Cysteine and glycine-rich protein 1 | HUMAN |
| sp\|P28676\|GRAN_HUMAN | Grancalcin | HUMAN |
| sp\|Q08722\|CD47_HUMAN | Leukocyte surface antigen CD47 | HUMAN |
| sp\|P35268\|RL22_HUMAN | 60S ribosomal protein L22 | HUMAN |
| sp\|P19022\|CADH2_HUMAN | Cadherin-2 | HUMAN |
| sp\|P28072\|PSB6_HUMAN | Proteasome subunit beta type-6 | HUMAN |
| sp\|P05067\|A4_HUMAN | Amyloid-beta precursor protein | HUMAN |
| sp\|Q8WXD2\|SCG3_HUMAN | Secretogranin-3 | HUMAN |
| sp\|O75937\|DNJC8_HUMAN | DnaJ homolog subfamily C member 8 | HUMAN |
| sp\|O43681\|GET3_HUMAN | ATPase GET3 | HUMAN |
| sp\|P11279\|LAMP1_HUMAN | Lysosome-associated membrane glycoprotein 1 | HUMAN |
| sp\|P04844\|RPN2_HUMAN | Dolichyl-diphosphooligosaccharide--protein glycosyltransferase subunit 2 | HUMAN |
| sp\|Q15819\|UB2V2_HUMAN | Ubiquitin-conjugating enzyme E2 variant 2 | HUMAN |
| sp\|O75821\|EIF3G_HUMAN | Eukaryotic translation initiation factor 3 subunit G | HUMAN |
| sp\|P11234\|RALB_HUMAN | Ras-related protein Ral-B | HUMAN |
| sp\|P17600\|SYN1_HUMAN | Synapsin-1 | HUMAN |
| sp\|Q9Y333\|LSM2_HUMAN | U6 snRNA-associated Sm-like protein LSm2 | HUMAN |
| sp\|Q08945\|SSRP1_HUMAN | FACT complex subunit SSRP1 | HUMAN |
| sp\|Q6PUV4\|CPLX2_HUMAN | Complexin-2 | HUMAN |
| sp\|Q15904\|VAS1_HUMAN | V-type proton ATPase subunit S1 | HUMAN |
| sp\|P59768\|GBG2_HUMAN | Guanine nucleotide-binding protein G(I)/G(S)/G(O) subunit gamma-2 | HUMAN |
| sp\|P55769\|NH2L1_HUMAN | NHP2-like protein 1 | HUMAN |

| | | |
|---|---|---|
| sp\|O95721\|SNP29_HUMAN | Synaptosomal-associated protein 29 | HUMAN |
| sp\|Q9NYU2\|UGGG1_HUMAN | UDP-glucose:glycoprotein glucosyltransferase 1 | HUMAN |
| sp\|P57088\|TMM33_HUMAN | Transmembrane protein 33 | HUMAN |
| sp\|Q15102\|PA1B3_HUMAN | Platelet-activating factor acetylhydrolase IB subunit alpha1 | HUMAN |
| sp\|Q53GG5\|PDLI3_HUMAN | PDZ and LIM domain protein 3 | HUMAN |
| sp\|Q13257\|MD2L1_HUMAN | Mitotic spindle assembly checkpoint protein MAD2A | HUMAN |
| sp\|Q9NPD3\|EXOS4_HUMAN | Exosome complex component RRP41 | HUMAN |
| sp\|O14994\|SYN3_HUMAN | Synapsin-3 | HUMAN |
| sp\|P61421\|VA0D1_HUMAN | V-type proton ATPase subunit d 1 | HUMAN |
| sp\|P62879\|GBB2_HUMAN | Guanine nucleotide-binding protein G(I)/G(S)/G(T) subunit beta-2 | HUMAN |
| sp\|Q9HD42\|CHM1A_HUMAN | Charged multivesicular body protein 1a | HUMAN |
| sp\|P50897\|PPT1_HUMAN | Palmitoyl-protein thioesterase 1 | HUMAN |
| sp\|O15511\|ARPC5_HUMAN | Actin-related protein 2/3 complex subunit 5 | HUMAN |
| sp\|O75915\|PRAF3_HUMAN | PRA1 family protein 3 | HUMAN |
| sp\|P51116\|FXR2_HUMAN | RNA-binding protein FXR2 | HUMAN |
| sp\|P52565\|GDIR1_HUMAN | Rho GDP-dissociation inhibitor 1 | HUMAN |
| sp\|Q92542\|NICA_HUMAN | Nicastrin | HUMAN |
| sp\|P62318\|SMD3_HUMAN | Small nuclear ribonucleoprotein Sm D3 | HUMAN |
| sp\|P47914\|RL29_HUMAN | 60S ribosomal protein L29 | HUMAN |
| sp\|P06753\|TPM3_HUMAN | Tropomyosin alpha-3 chain | HUMAN |
| sp\|Q96FW1\|OTUB1_HUMAN | Ubiquitin thioesterase OTUB1 | HUMAN |
| sp\|O00560\|SDCB1_HUMAN | Syntenin-1 | HUMAN |
| sp\|P13473\|LAMP2_HUMAN | Lysosome-associated membrane glycoprotein 2 | HUMAN |
| sp\|Q14254\|FLOT2_HUMAN | Flotillin-2 | HUMAN |
| sp\|Q6UXD5\|SE6L2_HUMAN | Seizure 6-like protein 2 | HUMAN |
| sp\|Q9BVK6\|TMED9_HUMAN | Transmembrane emp24 domain-containing protein 9 | HUMAN |
| sp\|Q15427\|SF3B4_HUMAN | Splicing factor 3B subunit 4 | HUMAN |
| sp\|Q9UEU0\|VTI1B_HUMAN | Vesicle transport through interaction with t-SNAREs homolog 1B | HUMAN |
| sp\|P60891\|PRPS1_HUMAN | Ribose-phosphate pyrophosphokinase 1 | HUMAN |
| sp\|Q99961\|SH3G1_HUMAN | Endophilin-A2 | HUMAN |
| sp\|O14579\|COPE_HUMAN | Coatomer subunit epsilon | HUMAN |
| sp\|Q9NRW1\|RAB6B_HUMAN | Ras-related protein Rab-6B | HUMAN |
| sp\|Q9H307\|PININ_HUMAN | Pinin | HUMAN |
| sp\|Q13595\|TRA2A_HUMAN | Transformer-2 protein homolog alpha | HUMAN |
| sp\|P62745\|RHOB_HUMAN | Rho-related GTP-binding protein RhoB | HUMAN |
| sp\|P61006\|RAB8A_HUMAN | Ras-related protein Rab-8A | HUMAN |
| sp\|O75475\|PSIP1_HUMAN | PC4 and SFRS1-interacting protein | HUMAN |

| | | |
|---|---|---|
| sp\|O60749\|SNX2_HUMAN | Sorting nexin-2 | HUMAN |
| sp\|Q07065\|CKAP4_HUMAN | Cytoskeleton-associated protein 4 | HUMAN |
| sp\|Q9UHV9\|PFD2_HUMAN | Prefoldin subunit 2 | HUMAN |
| sp\|Q00765\|REEP5_HUMAN | Receptor expression-enhancing protein 5 | HUMAN |
| sp\|P14174\|MIF_HUMAN | Macrophage migration inhibitory factor | HUMAN |
| sp\|Q92820\|GGH_HUMAN | Gamma-glutamyl hydrolase | HUMAN |
| sp\|P11117\|PPAL_HUMAN | Lysosomal acid phosphatase | HUMAN |
| sp\|O15258\|RER1_HUMAN | Protein RER1 | HUMAN |
| sp\|Q8WUH6\|TM263_HUMAN | Transmembrane protein 263 | HUMAN |
| sp\|P27449\|VATL_HUMAN | V-type proton ATPase 16 kDa proteolipid subunit c | HUMAN |
| sp\|Q9Y3B3\|TMED7_HUMAN | Transmembrane emp24 domain-containing protein 7 | HUMAN |
| sp\|P68363\|TBA1B_HUMAN | Tubulin alpha-1B chain | HUMAN |
| sp\|Q9H0U4\|RAB1B_HUMAN | Ras-related protein Rab-1B | HUMAN |
| sp\|Q16778\|H2B2E_HUMAN | Histone H2B type 2-E | HUMAN |
| sp\|P08754\|GNAI3_HUMAN | Guanine nucleotide-binding protein G(i) subunit alpha-3 | HUMAN |
| sp\|P39748\|FEN1_HUMAN | Flap endonuclease 1 | HUMAN |
| sp\|P15121\|ALDR_HUMAN | Aldo-keto reductase family 1 member B1 | HUMAN |
| sp\|O00154\|BACH_HUMAN | Cytosolic acyl coenzyme A thioester hydrolase | HUMAN |
| sp\|Q9HCJ6\|VAT1L_HUMAN | Synaptic vesicle membrane protein VAT-1 homolog-like | HUMAN |
| sp\|P62304\|RUXE_HUMAN | Small nuclear ribonucleoprotein E | HUMAN |
| sp\|P35080\|PROF2_HUMAN | Profilin-2 | HUMAN |
| sp\|Q8WXF7\|ATLA1_HUMAN | Atlastin-1 | HUMAN |
| sp\|P63172\|DYLT1_HUMAN | Dynein light chain Tctex-type 1 | HUMAN |
| sp\|Q9BY43\|CHM4A_HUMAN | Charged multivesicular body protein 4a | HUMAN |
| sp\|Q9BWW4\|SSBP3_HUMAN | Single-stranded DNA-binding protein 3 | HUMAN |
| sp\|Q92581\|SL9A6_HUMAN | Sodium/hydrogen exchanger 6 | HUMAN |
| sp\|Q16563\|SYPL1_HUMAN | Synaptophysin-like protein 1 | HUMAN |
| sp\|Q15041\|AR6P1_HUMAN | ADP-ribosylation factor-like protein 6-interacting protein 1 | HUMAN |
| sp\|P68402\|PA1B2_HUMAN | Platelet-activating factor acetylhydrolase IB subunit alpha2 | HUMAN |
| sp\|P14209\|CD99_HUMAN | CD99 antigen | HUMAN |
| sp\|P42766\|RL35_HUMAN | 60S ribosomal protein L35 | HUMAN |
| sp\|P52294\|IMA5_HUMAN | Importin subunit alpha-5 | HUMAN |
| sp\|P61081\|UBC12_HUMAN | NEDD8-conjugating enzyme Ubc12 | HUMAN |
| sp\|Q6FI81\|CPIN1_HUMAN | Anamorsin | HUMAN |
| sp\|Q9P0T7\|TMEM9_HUMAN | Proton-transporting V-type ATPase complex assembly regulator TMEM9 | HUMAN |
| sp\|P98179\|RBM3_HUMAN | RNA-binding protein 3 | HUMAN |
| sp\|Q9UH03\|SEPT3_HUMAN | Neuronal-specific septin-3 | HUMAN |

| | | |
|---|---|---|
| sp\|Q92572\|AP3S1_HUMAN | AP-3 complex subunit sigma-1 | HUMAN |
| sp\|O75781\|PALM_HUMAN | Paralemmin-1 | HUMAN |
| sp\|Q9UHG3\|PCYOX_HUMAN | Prenylcysteine oxidase 1 | HUMAN |
| sp\|P63096\|GNAI1_HUMAN | Guanine nucleotide-binding protein G(i) subunit alpha-1 | HUMAN |
| sp\|Q5ZPR3\|CD276_HUMAN | CD276 antigen | HUMAN |
| sp\|P49411\|EFTU_HUMAN | Elongation factor Tu, mitochondrial | HUMAN |
| sp\|Q9H3Z4\|DNJC5_HUMAN | DnaJ homolog subfamily C member 5 | HUMAN |
| sp\|Q5IJ48\|CRUM2_HUMAN | Protein crumbs homolog 2 | HUMAN |
| sp\|Q9Y2V2\|CHSP1_HUMAN | Calcium-regulated heat-stable protein 1 | HUMAN |
| sp\|Q04837\|SSBP_HUMAN | Single-stranded DNA-binding protein, mitochondrial | HUMAN |
| sp\|P20962\|PTMS_HUMAN | Parathymosin | HUMAN |
| sp\|Q99805\|TM9S2_HUMAN | Transmembrane 9 superfamily member 2 | HUMAN |
| sp\|P48449\|LSS_HUMAN | Lanosterol synthase | HUMAN |
| sp\|Q92544\|TM9S4_HUMAN | Transmembrane 9 superfamily member 4 | HUMAN |
| sp\|P61604\|CH10_HUMAN | 10 kDa heat shock protein, mitochondrial | HUMAN |
| sp\|Q9BW30\|TPPP3_HUMAN | Tubulin polymerization-promoting protein family member 3 | HUMAN |
| sp\|Q96EP5\|DAZP1_HUMAN | DAZ-associated protein 1 | HUMAN |
| sp\|O94772\|LY6H_HUMAN | Lymphocyte antigen 6H | HUMAN |
| sp\|Q99436\|PSB7_HUMAN | Proteasome subunit beta type-7 | HUMAN |
| sp\|P45880\|VDAC2_HUMAN | Voltage-dependent anion-selective channel protein 2 | HUMAN |
| sp\|Q92747\|ARC1A_HUMAN | Actin-related protein 2/3 complex subunit 1A | HUMAN |
| sp\|P08138\|TNR16_HUMAN | Tumor necrosis factor receptor superfamily member 16 | HUMAN |
| sp\|P20336\|RAB3A_HUMAN | Ras-related protein Rab-3A | HUMAN |
| sp\|P30044\|PRDX5_HUMAN | Peroxiredoxin-5, mitochondrial | HUMAN |
| sp\|P41217\|OX2G_HUMAN | OX-2 membrane glycoprotein | HUMAN |
| sp\|Q9H3N1\|TMX1_HUMAN | Thioredoxin-related transmembrane protein 1 | HUMAN |
| sp\|Q96G97\|BSCL2_HUMAN | Seipin | HUMAN |
| sp\|Q8N5K1\|CISD2_HUMAN | CDGSH iron-sulfur domain-containing protein 2 | HUMAN |
| sp\|P51674\|GPM6A_HUMAN | Neuronal membrane glycoprotein M6-a | HUMAN |
| sp\|Q9UNK0\|STX8_HUMAN | Syntaxin-8 | HUMAN |
| sp\|P13987\|CD59_HUMAN | CD59 glycoprotein | HUMAN |
| sp\|Q12824\|SNF5_HUMAN | SWI/SNF-related matrix-associated actin-dependent regulator of chromatin subfamily B member 1 | HUMAN |
| sp\|O75935\|DCTN3_HUMAN | Dynactin subunit 3 | HUMAN |
| sp\|Q71UI9\|H2AV_HUMAN | Histone H2A.V | HUMAN |
| sp\|Q99614\|TTC1_HUMAN | Tetratricopeptide repeat protein 1 | HUMAN |
| sp\|Q9NWB6\|ARGL1_HUMAN | Arginine and glutamate-rich protein 1 | HUMAN |

| | | |
|---|---|---|
| sp\|P12236\|ADT3_HUMAN | ADP/ATP translocase 3 | HUMAN |
| sp\|P61266\|STX1B_HUMAN | Syntaxin-1B | HUMAN |
| sp\|Q969X5\|ERGI1_HUMAN | Endoplasmic reticulum-Golgi intermediate compartment protein 1 | HUMAN |
| sp\|P33947\|ERD22_HUMAN | ER lumen protein-retaining receptor 2 | HUMAN |
| sp\|P41208\|CETN2_HUMAN | Centrin-2 | HUMAN |
| sp\|Q8TCT9\|HM13_HUMAN | Minor histocompatibility antigen H13 | HUMAN |
| sp\|Q9NPA8\|ENY2_HUMAN | Transcription and mRNA export factor ENY2 | HUMAN |
| sp\|Q7Z7H5\|TMED4_HUMAN | Transmembrane emp24 domain-containing protein 4 | HUMAN |
| sp\|Q9UDY4\|DNJB4_HUMAN | DnaJ homolog subfamily B member 4 | HUMAN |
| sp\|Q8ND24\|RN214_HUMAN | RING finger protein 214 | HUMAN |
| sp\|P68036\|UB2L3_HUMAN | Ubiquitin-conjugating enzyme E2 L3 | HUMAN |
| sp\|P63218\|GBG5_HUMAN | Guanine nucleotide-binding protein G(I)/G(S)/G(O) subunit gamma-5 | HUMAN |
| sp\|Q8WUD1\|RAB2B_HUMAN | Ras-related protein Rab-2B | HUMAN |
| sp\|Q14320\|FA50A_HUMAN | Protein FAM50A | HUMAN |
| sp\|Q9UHN6\|CEIP2_HUMAN | Cell surface hyaluronidase | HUMAN |
| sp\|Q8TAC9\|SCAM5_HUMAN | Secretory carrier-associated membrane protein 5 | HUMAN |
| sp\|Q9UBM7\|DHCR7_HUMAN | 7-dehydrocholesterol reductase | HUMAN |
| sp\|O15145\|ARPC3_HUMAN | Actin-related protein 2/3 complex subunit 3 | HUMAN |
| sp\|Q9Y2Q0\|AT8A1_HUMAN | Phospholipid-transporting ATPase IA | HUMAN |
| sp\|P47755\|CAZA2_HUMAN | F-actin-capping protein subunit alpha-2 | HUMAN |
| sp\|Q9UL26\|RB22A_HUMAN | Ras-related protein Rab-22A | HUMAN |
| sp\|Q13449\|LSAMP_HUMAN | Limbic system-associated membrane protein | HUMAN |
| sp\|P83916\|CBX1_HUMAN | Chromobox protein homolog 1 | HUMAN |
| sp\|P07858\|CATB_HUMAN | Cathepsin B | HUMAN |
| sp\|Q8N0U8\|VKORL_HUMAN | Vitamin K epoxide reductase complex subunit 1-like protein 1 | HUMAN |
| sp\|Q9NUP9\|LIN7C_HUMAN | Protein lin-7 homolog C | HUMAN |
| sp\|P14923\|PLAK_HUMAN | Junction plakoglobin | HUMAN |
| sp\|O60641\|AP180_HUMAN | Clathrin coat assembly protein AP180 | HUMAN |
| sp\|Q9Y296\|TPPC4_HUMAN | Trafficking protein particle complex subunit 4 | HUMAN |
| sp\|P04792\|HSPB1_HUMAN | Heat shock protein beta-1 | HUMAN |
| sp\|Q5BJF2\|SGMR2_HUMAN | Sigma intracellular receptor 2 | HUMAN |
| sp\|O95197\|RTN3_HUMAN | Reticulon-3 | HUMAN |
| sp\|Q9NV96\|CC50A_HUMAN | Cell cycle control protein 50A | HUMAN |
| sp\|P62861\|RS30_HUMAN | FAU ubiquitin-like and ribosomal protein S30 | HUMAN |
| sp\|Q01085\|TIAR_HUMAN | Nucleolysin TIAR | HUMAN |
| sp\|Q9UNF1\|MAGD2_HUMAN | Melanoma-associated antigen D2 | HUMAN |
| sp\|Q96DA2\|RB39B_HUMAN | Ras-related protein Rab-39B | HUMAN |
| sp\|P37802\|TAGL2_HUMAN | Transgelin-2 | HUMAN |

| | | |
|---|---|---|
| sp\|P63167\|DYL1_HUMAN | Dynein light chain 1, cytoplasmic | HUMAN |
| sp\|P20338\|RAB4A_HUMAN | Ras-related protein Rab-4A | HUMAN |
| sp\|Q12931\|TRAP1_HUMAN | Heat shock protein 75 kDa, mitochondrial | HUMAN |
| sp\|Q8WZA9\|IRGQ_HUMAN | Immunity-related GTPase family Q protein | HUMAN |
| sp\|O00592\|PODXL_HUMAN | Podocalyxin | HUMAN |
| sp\|P06396\|GELS_HUMAN | Gelsolin | HUMAN |
| sp\|P68371\|TBB4B_HUMAN | Tubulin beta-4B chain | HUMAN |
| sp\|Q13885\|TBB2A_HUMAN | Tubulin beta-2A chain | HUMAN |
| sp\|P17858\|PFKAL_HUMAN | ATP-dependent 6-phosphofructokinase, liver type | HUMAN |
| sp\|Q15836\|VAMP3_HUMAN | Vesicle-associated membrane protein 3 | HUMAN |
| sp\|Q9H169\|STMN4_HUMAN | Stathmin-4 | HUMAN |
| sp\|P61073\|CXCR4_HUMAN | C-X-C chemokine receptor type 4 | HUMAN |
| sp\|Q8TC12\|RDH11_HUMAN | Retinol dehydrogenase 11 | HUMAN |
| sp\|Q9BZF1\|OSBL8_HUMAN | Oxysterol-binding protein-related protein 8 | HUMAN |
| sp\|Q5T1Q4\|S35F1_HUMAN | Solute carrier family 35 member F1 | HUMAN |
| sp\|Q08380\|LG3BP_HUMAN | Galectin-3-binding protein | HUMAN |
| sp\|O95674\|CDS2_HUMAN | Phosphatidate cytidylyltransferase 2 | HUMAN |
| sp\|O95297\|MPZL1_HUMAN | Myelin protein zero-like protein 1 | HUMAN |
| sp\|Q9UI12\|VATH_HUMAN | V-type proton ATPase subunit H | HUMAN |
| sp\|Q9NZ01\|TECR_HUMAN | Very-long-chain enoyl-CoA reductase | HUMAN |
| sp\|Q9NRX5\|SERC1_HUMAN | Serine incorporator 1 | HUMAN |
| sp\|Q9H0Q3\|FXYD6_HUMAN | FXYD domain-containing ion transport regulator 6 | HUMAN |
| sp\|Q99873\|ANM1_HUMAN | Protein arginine N-methyltransferase 1 | HUMAN |
| sp\|Q99623\|PHB2_HUMAN | Prohibitin-2 | HUMAN |
| sp\|Q969E2\|SCAM4_HUMAN | Secretory carrier-associated membrane protein 4 | HUMAN |
| sp\|Q92823\|NRCAM_HUMAN | Neuronal cell adhesion molecule | HUMAN |
| sp\|Q92520\|FAM3C_HUMAN | Protein FAM3C | HUMAN |
| sp\|Q5BJH7\|YIF1B_HUMAN | Protein YIF1B | HUMAN |
| sp\|Q13867\|BLMH_HUMAN | Bleomycin hydrolase | HUMAN |
| sp\|Q00325\|MPCP_HUMAN | Phosphate carrier protein, mitochondrial | HUMAN |
| sp\|P61278\|SMS_HUMAN | Somatostatin | HUMAN |
| sp\|P61009\|SPCS3_HUMAN | Signal peptidase complex subunit 3 | HUMAN |
| sp\|P54289\|CA2D1_HUMAN | Voltage-dependent calcium channel subunit alpha-2/delta-1 | HUMAN |
| sp\|P42785\|PCP_HUMAN | Lysosomal Pro-X carboxypeptidase | HUMAN |
| sp\|P05386\|RLA1_HUMAN | 60S acidic ribosomal protein P1 | HUMAN |
| sp\|P04216\|THY1_HUMAN | Thy-1 membrane glycoprotein | HUMAN |
| sp\|O95864\|FADS2_HUMAN | Acyl-CoA 6-desaturase | HUMAN |
| sp\|O76062\|ERG24_HUMAN | Delta(14)-sterol reductase TM7SF2 | HUMAN |
| sp\|O43761\|SNG3_HUMAN | Synaptogyrin-3 | HUMAN |

| | | |
|---|---|---|
| sp\|P52948\|NUP98_HUMAN | Nuclear pore complex protein Nup98-Nup96 | HUMAN |
| sp\|O60231\|DHX16_HUMAN | Pre-mRNA-splicing factor ATP-dependent RNA helicase DHX16 | HUMAN |
| sp\|Q9Y639\|NPTN_HUMAN | Neuroplastin | HUMAN |
| sp\|Q9UHA4\|LTOR3_HUMAN | Ragulator complex protein LAMTOR3 | HUMAN |
| sp\|O95772\|STR3N_HUMAN | STARD3 N-terminal-like protein | HUMAN |
| sp\|Q15404\|RSU1_HUMAN | Ras suppressor protein 1 | HUMAN |
| sp\|P08962\|CD63_HUMAN | CD63 antigen | HUMAN |
| sp\|Q92692\|NECT2_HUMAN | Nectin-2 | HUMAN |
| sp\|Q9H0E2\|TOLIP_HUMAN | Toll-interacting protein | HUMAN |
| sp\|Q92733\|PRCC_HUMAN | Proline-rich protein PRCC | HUMAN |
| sp\|Q9Y2S6\|TMA7_HUMAN | Translation machinery-associated protein 7 | HUMAN |
| sp\|P43307\|SSRA_HUMAN | Translocon-associated protein subunit alpha | HUMAN |
| sp\|Q9NQ48\|LZTL1_HUMAN | Leucine zipper transcription factor-like protein 1 | HUMAN |
| sp\|Q9BPW8\|NIPS1_HUMAN | Protein NipSnap homolog 1 | HUMAN |
| sp\|Q9H1E3\|NUCKS_HUMAN | Nuclear ubiquitous casein and cyclin-dependent kinase substrate 1 | HUMAN |
| sp\|P22392\|NDKB_HUMAN | Nucleoside diphosphate kinase B | HUMAN |
| sp\|P56377\|AP1S2_HUMAN | AP-1 complex subunit sigma-2 | HUMAN |
| sp\|P63272\|SPT4H_HUMAN | Transcription elongation factor SPT4 | HUMAN |
| sp\|P52943\|CRIP2_HUMAN | Cysteine-rich protein 2 | HUMAN |
| sp\|Q9BTV4\|TMM43_HUMAN | Transmembrane protein 43 | HUMAN |
| sp\|Q15125\|EBP_HUMAN | 3-beta-hydroxysteroid-Delta(8),Delta(7)-isomerase | HUMAN |
| sp\|Q02952\|AKA12_HUMAN | A-kinase anchor protein 12 | HUMAN |
| sp\|Q8NC96\|NECP1_HUMAN | Adaptin ear-binding coat-associated protein 1 | HUMAN |
| sp\|O75347\|TBCA_HUMAN | Tubulin-specific chaperone A | HUMAN |
| sp\|Q15382\|RHEB_HUMAN | GTP-binding protein Rheb | HUMAN |
| sp\|Q9NP90\|RAB9B_HUMAN | Ras-related protein Rab-9B | HUMAN |
| sp\|Q8WUX9\|CHMP7_HUMAN | Charged multivesicular body protein 7 | HUMAN |
| sp\|Q92734\|TFG_HUMAN | Protein TFG | HUMAN |
| sp\|P51571\|SSRD_HUMAN | Translocon-associated protein subunit delta | HUMAN |
| sp\|Q16572\|VACHT_HUMAN | Vesicular acetylcholine transporter | HUMAN |
| sp\|Q8WVC6\|DCAKD_HUMAN | Dephospho-CoA kinase domain-containing protein | HUMAN |
| sp\|Q9HD45\|TM9S3_HUMAN | Transmembrane 9 superfamily member 3 | HUMAN |
| sp\|Q8TDB8\|GTR14_HUMAN | Solute carrier family 2, facilitated glucose transporter member 14 | HUMAN |
| sp\|Q8IUE6\|H2A2B_HUMAN | Histone H2A type 2-B | HUMAN |
| sp\|Q13123\|RED_HUMAN | Protein Red | HUMAN |
| sp\|Q16851\|UGPA_HUMAN | UTP--glucose-1-phosphate uridylyltransferase | HUMAN |
| sp\|Q9BS26\|ERP44_HUMAN | Endoplasmic reticulum resident protein 44 | HUMAN |

| | | |
|---|---|---|
| sp\|P13521\|SCG2_HUMAN | Secretogranin-2 | HUMAN |
| sp\|P09622\|DLDH_HUMAN | Dihydrolipoyl dehydrogenase, mitochondrial | HUMAN |

**Supp. Table S2.**

**Supplementary Table S2: List of 936 proteins bound by PS-NP, detected by LC-MS/MS**

| Accession | Protein Name | Species |
|---|---|---|
| **sp\|Q14204\|DYHC1_HUMAN** | Cytoplasmic dynein 1 heavy chain 1 | HUMAN |
| **sp\|P78527\|PRKDC_HUMAN** | DNA-dependent protein kinase catalytic subunit | HUMAN |

| | | |
|---|---|---|
| **sp\|Q13813\|SPTN1_HUMAN** | Spectrin alpha chain, non-erythrocytic 1 | HUMAN |
| **sp\|P35580\|MYH10_HUMAN** | Myosin-10 | HUMAN |
| **sp\|P46821\|MAP1B_HUMAN** | Microtubule-associated protein 1B | HUMAN |
| **sp\|Q14152\|EIF3A_HUMAN** | Eukaryotic translation initiation factor 3 subunit A | HUMAN |
| **sp\|Q01082\|SPTB2_HUMAN** | Spectrin beta chain, non-erythrocytic 1 | HUMAN |
| **sp\|Q00610\|CLH1_HUMAN** | Clathrin heavy chain 1 | HUMAN |
| **sp\|O75533\|SF3B1_HUMAN** | Splicing factor 3B subunit 1 | HUMAN |
| **sp\|P33176\|KINH_HUMAN** | Kinesin-1 heavy chain | HUMAN |
| **sp\|P08238\|HS90B_HUMAN** | Heat shock protein HSP 90-beta | HUMAN |
| **sp\|O75643\|U520_HUMAN** | U5 small nuclear ribonucleoprotein 200 kDa helicase | HUMAN |
| **sp\|Q04637\|IF4G1_HUMAN** | Eukaryotic translation initiation factor 4 gamma 1 | HUMAN |
| **sp\|P11137\|MTAP2_HUMAN** | Microtubule-associated protein 2 | HUMAN |
| **sp\|P55060\|XPO2_HUMAN** | Exportin-2 | HUMAN |
| **sp\|P07437\|TBB5_HUMAN** | Tubulin beta chain | HUMAN |
| **sp\|P12270\|TPR_HUMAN** | Nucleoprotein TPR | HUMAN |
| **sp\|Q15393\|SF3B3_HUMAN** | Splicing factor 3B subunit 3 | HUMAN |
| **sp\|P78347\|GTF2I_HUMAN** | General transcription factor II-I | HUMAN |
| **sp\|P31948\|STIP1_HUMAN** | Stress-induced-phosphoprotein 1 | HUMAN |
| **sp\|P52272\|HNRPM_HUMAN** | Heterogeneous nuclear ribonucleoprotein M | HUMAN |
| **sp\|Q6P2Q9\|PRP8_HUMAN** | Pre-mRNA-processing-splicing factor 8 | HUMAN |
| **sp\|P13639\|EF2_HUMAN** | Elongation factor 2 | HUMAN |
| **sp\|Q08211\|DHX9_HUMAN** | ATP-dependent RNA helicase A | HUMAN |
| **sp\|P68104\|EF1A1_HUMAN** | Elongation factor 1-alpha 1 | HUMAN |
| **sp\|Q14839\|CHD4_HUMAN** | Chromodomain-helicase-DNA-binding protein 4 | HUMAN |
| **sp\|P11586\|C1TC_HUMAN** | C-1-tetrahydrofolate synthase, cytoplasmic | HUMAN |
| **sp\|P07814\|SYEP_HUMAN** | Bifunctional glutamate/proline--tRNA ligase | HUMAN |
| **sp\|Q00839\|HNRPU_HUMAN** | Heterogeneous nuclear ribonucleoprotein U | HUMAN |
| **sp\|Q13263\|TIF1B_HUMAN** | Transcription intermediary factor 1-beta | HUMAN |
| **sp\|Q71U36\|TBA1A_HUMAN** | Tubulin alpha-1A chain | HUMAN |
| **sp\|Q15459\|SF3A1_HUMAN** | Splicing factor 3A subunit 1 | HUMAN |
| **sp\|P21333\|FLNA_HUMAN** | Filamin-A | HUMAN |
| **sp\|Q92945\|FUBP2_HUMAN** | Far upstream element-binding protein 2 | HUMAN |
| **sp\|Q13435\|SF3B2_HUMAN** | Splicing factor 3B subunit 2 | HUMAN |
| **sp\|Q92499\|DDX1_HUMAN** | ATP-dependent RNA helicase DDX1 | HUMAN |
| **sp\|P50395\|GDIB_HUMAN** | Rab GDP dissociation inhibitor beta | HUMAN |
| **sp\|P43243\|MATR3_HUMAN** | Matrin-3 | HUMAN |
| **sp\|P78344\|IF4G2_HUMAN** | Eukaryotic translation initiation factor 4 gamma 2 | HUMAN |
| **sp\|P26038\|MOES_HUMAN** | Moesin | HUMAN |
| **sp\|Q92841\|DDX17_HUMAN** | Probable ATP-dependent RNA helicase DDX17 | HUMAN |
| **sp\|P11142\|HSP7C_HUMAN** | Heat shock cognate 71 kDa protein | HUMAN |
| **sp\|Q9P2J5\|SYLC_HUMAN** | Leucine--tRNA ligase, cytoplasmic | HUMAN |
| **sp\|Q14194\|DPYL1_HUMAN** | Dihydropyrimidinase-related protein 1 | HUMAN |

| | | |
|---|---|---|
| **sp\|Q99613\|EIF3C_HUMAN** | Eukaryotic translation initiation factor 3 subunit C | HUMAN |
| **sp\|P11940\|PABP1_HUMAN** | Polyadenylate-binding protein 1 | HUMAN |
| **sp\|Q93009\|UBP7_HUMAN** | Ubiquitin carboxyl-terminal hydrolase 7 | HUMAN |
| **sp\|O60282\|KIF5C_HUMAN** | Kinesin heavy chain isoform 5C | HUMAN |
| **sp\|Q8TAQ2\|SMRC2_HUMAN** | SWI/SNF complex subunit SMARCC2 | HUMAN |
| **sp\|Q9BXP5\|SRRT_HUMAN** | Serrate RNA effector molecule homolog | HUMAN |
| **sp\|Q12906\|ILF3_HUMAN** | Interleukin enhancer-binding factor 3 | HUMAN |
| **sp\|Q96AE4\|FUBP1_HUMAN** | Far upstream element-binding protein 1 | HUMAN |
| **sp\|O43390\|HNRPR_HUMAN** | Heterogeneous nuclear ribonucleoprotein R | HUMAN |
| **sp\|P55884\|EIF3B_HUMAN** | Eukaryotic translation initiation factor 3 subunit B | HUMAN |
| **sp\|Q07866\|KLC1_HUMAN** | Kinesin light chain 1 | HUMAN |
| **sp\|P61978\|HNRPK_HUMAN** | Heterogeneous nuclear ribonucleoprotein K | HUMAN |
| **sp\|P14618\|KPYM_HUMAN** | Pyruvate kinase PKM | HUMAN |
| **sp\|P04264\|K2C1_HUMAN** | Keratin, type II cytoskeletal 1 | HUMAN |
| **sp\|P09874\|PARP1_HUMAN** | Poly [ADP-ribose] polymerase 1 | HUMAN |
| **sp\|P13010\|XRCC5_HUMAN** | X-ray repair cross-complementing protein 5 | HUMAN |
| **sp\|Q16531\|DDB1_HUMAN** | DNA damage-binding protein 1 | HUMAN |
| **sp\|P50570\|DYN2_HUMAN** | Dynamin-2 | HUMAN |
| **sp\|P14868\|SYDC_HUMAN** | Aspartate--tRNA ligase, cytoplasmic | HUMAN |
| **sp\|Q8N163\|CCAR2_HUMAN** | Cell cycle and apoptosis regulator protein 2 | HUMAN |
| **sp\|O94979\|SC31A_HUMAN** | Protein transport protein Sec31A | HUMAN |
| **sp\|P50990\|TCPQ_HUMAN** | T-complex protein 1 subunit theta | HUMAN |
| **sp\|P22626\|ROA2_HUMAN** | Heterogeneous nuclear ribonucleoproteins A2/B1 | HUMAN |
| **sp\|O14776\|TCRG1_HUMAN** | Transcription elongation regulator 1 | HUMAN |
| **sp\|P12956\|XRCC6_HUMAN** | X-ray repair cross-complementing protein 6 | HUMAN |
| **sp\|Q14203\|DCTN1_HUMAN** | Dynactin subunit 1 | HUMAN |
| **sp\|P63261\|ACTG_HUMAN** | Actin, cytoplasmic 2 | HUMAN |
| **sp\|P29401\|TKT_HUMAN** | Transketolase | HUMAN |
| **sp\|Q00341\|VIGLN_HUMAN** | Vigilin | HUMAN |
| **sp\|P25205\|MCM3_HUMAN** | DNA replication licensing factor MCM3 | HUMAN |
| **sp\|Q9Y262\|EIF3L_HUMAN** | Eukaryotic translation initiation factor 3 subunit L | HUMAN |
| **sp\|P47897\|SYQ_HUMAN** | Glutamine--tRNA ligase | HUMAN |
| **sp\|Q14008\|CKAP5_HUMAN** | Cytoskeleton-associated protein 5 | HUMAN |
| **sp\|Q7Z6Z7\|HUWE1_HUMAN** | E3 ubiquitin-protein ligase HUWE1 | HUMAN |
| **sp\|P23588\|IF4B_HUMAN** | Eukaryotic translation initiation factor 4B | HUMAN |
| **sp\|P78371\|TCPB_HUMAN** | T-complex protein 1 subunit beta | HUMAN |
| **sp\|P22314\|UBA1_HUMAN** | Ubiquitin-like modifier-activating enzyme 1 | HUMAN |
| **sp\|P63010\|AP2B1_HUMAN** | AP-2 complex subunit beta | HUMAN |
| **sp\|Q15029\|U5S1_HUMAN** | 116 kDa U5 small nuclear ribonucleoprotein component | HUMAN |
| **sp\|P33992\|MCM5_HUMAN** | DNA replication licensing factor MCM5 | HUMAN |
| **sp\|Q9NZI8\|IF2B1_HUMAN** | Insulin-like growth factor 2 mRNA-binding protein 1 | HUMAN |

| | | |
|---|---|---|
| **sp\|Q9UHD8\|SEPT9_HUMAN** | Septin-9 | HUMAN |
| **sp\|Q9BPU6\|DPYL5_HUMAN** | Dihydropyrimidinase-related protein 5 | HUMAN |
| **sp\|P23921\|RIR1_HUMAN** | Ribonucleoside-diphosphate reductase large subunit | HUMAN |
| **sp\|O60341\|KDM1A_HUMAN** | Lysine-specific histone demethylase 1A | HUMAN |
| **sp\|P49368\|TCPG_HUMAN** | T-complex protein 1 subunit gamma | HUMAN |
| **sp\|Q9H4M9\|EHD1_HUMAN** | EH domain-containing protein 1 | HUMAN |
| **sp\|P41252\|SYIC_HUMAN** | Isoleucine--tRNA ligase, cytoplasmic | HUMAN |
| **sp\|Q13409\|DC1I2_HUMAN** | Cytoplasmic dynein 1 intermediate chain 2 | HUMAN |
| **sp\|Q16555\|DPYL2_HUMAN** | Dihydropyrimidinase-related protein 2 | HUMAN |
| **sp\|Q9Y230\|RUVB2_HUMAN** | RuvB-like 2 | HUMAN |
| **sp\|P13645\|K1C10_HUMAN** | Keratin, type I cytoskeletal 10 | HUMAN |
| **sp\|O00571\|DDX3X_HUMAN** | ATP-dependent RNA helicase DDX3X | HUMAN |
| **sp\|Q9UBT2\|SAE2_HUMAN** | SUMO-activating enzyme subunit 2 | HUMAN |
| **sp\|Q7Z4S6\|KI21A_HUMAN** | Kinesin-like protein KIF21A | HUMAN |
| **sp\|Q9Y3I0\|RTCB_HUMAN** | RNA-splicing ligase RtcB homolog | HUMAN |
| **sp\|Q7L014\|DDX46_HUMAN** | Probable ATP-dependent RNA helicase DDX46 | HUMAN |
| **sp\|Q8NC51\|PAIRB_HUMAN** | Plasminogen activator inhibitor 1 RNA-binding protein | HUMAN |
| **sp\|P07900\|HS90A_HUMAN** | Heat shock protein HSP 90-alpha | HUMAN |
| **sp\|P50991\|TCPD_HUMAN** | T-complex protein 1 subunit delta | HUMAN |
| **sp\|P08670\|VIME_HUMAN** | Vimentin | HUMAN |
| **sp\|Q1KMD3\|HNRL2_HUMAN** | Heterogeneous nuclear ribonucleoprotein U-like protein 2 | HUMAN |
| **sp\|Q5SW79\|CE170_HUMAN** | Centrosomal protein of 170 kDa | HUMAN |
| **sp\|P19338\|NUCL_HUMAN** | Nucleolin | HUMAN |
| **sp\|Q9UHX1\|PUF60_HUMAN** | Poly(U)-binding-splicing factor PUF60 | HUMAN |
| **sp\|P35606\|COPB2_HUMAN** | Coatomer subunit beta' | HUMAN |
| **sp\|P11216\|PYGB_HUMAN** | Glycogen phosphorylase, brain form | HUMAN |
| **sp\|Q7L1Q6\|5MP2_HUMAN** | eIF5-mimic protein 2 | HUMAN |
| **sp\|P14866\|HNRPL_HUMAN** | Heterogeneous nuclear ribonucleoprotein L | HUMAN |
| **sp\|Q9HAV4\|XPO5_HUMAN** | Exportin-5 | HUMAN |
| **sp\|Q9NVA2\|SEP11_HUMAN** | Septin-11 | HUMAN |
| **sp\|Q9H9A6\|LRC40_HUMAN** | Leucine-rich repeat-containing protein 40 | HUMAN |
| **sp\|Q16643\|DREB_HUMAN** | Drebrin | HUMAN |
| **sp\|P31943\|HNRH1_HUMAN** | Heterogeneous nuclear ribonucleoprotein H | HUMAN |
| **sp\|Q9NTZ6\|RBM12_HUMAN** | RNA-binding protein 12 | HUMAN |
| **sp\|Q9BZZ5\|API5_HUMAN** | Apoptosis inhibitor 5 | HUMAN |
| **sp\|Q9Y265\|RUVB1_HUMAN** | RuvB-like 1 | HUMAN |
| **sp\|Q8TAT6\|NPL4_HUMAN** | Nuclear protein localization protein 4 homolog | HUMAN |
| **sp\|O14531\|DPYL4_HUMAN** | Dihydropyrimidinase-related protein 4 | HUMAN |
| **sp\|Q12874\|SF3A3_HUMAN** | Splicing factor 3A subunit 3 | HUMAN |
| **sp\|P48643\|TCPE_HUMAN** | T-complex protein 1 subunit epsilon | HUMAN |

| | | |
|---|---|---|
| **sp\|P63244\|RACK1_HUMAN** | Receptor of activated protein C kinase 1 | HUMAN |
| **sp\|Q14566\|MCM6_HUMAN** | DNA replication licensing factor MCM6 | HUMAN |
| **sp\|P06733\|ENOA_HUMAN** | Alpha-enolase | HUMAN |
| **sp\|P54136\|SYRC_HUMAN** | Arginine--tRNA ligase, cytoplasmic | HUMAN |
| **sp\|P05198\|IF2A_HUMAN** | Eukaryotic translation initiation factor 2 subunit 1 | HUMAN |
| **sp\|Q02790\|FKBP4_HUMAN** | Peptidyl-prolyl cis-trans isomerase FKBP4 | HUMAN |
| **sp\|Q15046\|SYK_HUMAN** | Lysine--tRNA ligase | HUMAN |
| **sp\|O43295\|SRGP3_HUMAN** | SLIT-ROB Rho GTPase-activating protein 3 | HUMAN |
| **sp\|P56192\|SYMC_HUMAN** | Methionine--tRNA ligase, cytoplasmic | HUMAN |
| **sp\|P30153\|2AAA_HUMAN** | Serine/threonine-protein phosphatase 2A 65 kDa regulatory subunit A alpha isoform | HUMAN |
| **sp\|P40227\|TCPZ_HUMAN** | T-complex protein 1 subunit zeta | HUMAN |
| **sp\|O43602\|DCX_HUMAN** | Neuronal migration protein doublecortin | HUMAN |
| **sp\|P23246\|SFPQ_HUMAN** | Splicing factor, proline- and glutamine-rich | HUMAN |
| **sp\|P62258\|1433E_HUMAN** | 14-3-3 protein epsilon | HUMAN |
| **sp\|Q14157\|UBP2L_HUMAN** | Ubiquitin-associated protein 2-like | HUMAN |
| **sp\|P23396\|RS3_HUMAN** | 40S ribosomal protein S3 | HUMAN |
| **sp\|Q8IX12\|CCAR1_HUMAN** | Cell division cycle and apoptosis regulator protein 1 | HUMAN |
| **sp\|Q96KP4\|CNDP2_HUMAN** | Cytosolic non-specific dipeptidase | HUMAN |
| **sp\|Q8N8S7\|ENAH_HUMAN** | Protein enabled homolog | HUMAN |
| **sp\|O95782\|AP2A1_HUMAN** | AP-2 complex subunit alpha-1 | HUMAN |
| **sp\|P26599\|PTBP1_HUMAN** | Polypyrimidine tract-binding protein 1 | HUMAN |
| **sp\|Q13200\|PSMD2_HUMAN** | 26S proteasome non-ATPase regulatory subunit 2 | HUMAN |
| **sp\|P30101\|PDIA3_HUMAN** | Protein disulfide-isomerase A3 | HUMAN |
| **sp\|Q15019\|SEPT2_HUMAN** | Septin-2 | HUMAN |
| **sp\|O43719\|HTSF1_HUMAN** | HIV Tat-specific factor 1 | HUMAN |
| **sp\|Q14974\|IMB1_HUMAN** | Importin subunit beta-1 | HUMAN |
| **sp\|P60228\|EIF3E_HUMAN** | Eukaryotic translation initiation factor 3 subunit E | HUMAN |
| **sp\|Q12905\|ILF2_HUMAN** | Interleukin enhancer-binding factor 2 | HUMAN |
| **sp\|P26641\|EF1G_HUMAN** | Elongation factor 1-gamma | HUMAN |
| **sp\|P17858\|PFKAL_HUMAN** | ATP-dependent 6-phosphofructokinase, liver type | HUMAN |
| **sp\|Q9UQB8\|BAIP2_HUMAN** | Brain-specific angiogenesis inhibitor 1-associated protein 2 | HUMAN |
| **sp\|Q14195\|DPYL3_HUMAN** | Dihydropyrimidinase-related protein 3 | HUMAN |
| **sp\|P17844\|DDX5_HUMAN** | Probable ATP-dependent RNA helicase DDX5 | HUMAN |
| **sp\|P38919\|IF4A3_HUMAN** | Eukaryotic initiation factor 4A-III | HUMAN |
| **sp\|Q96QK1\|VPS35_HUMAN** | Vacuolar protein sorting-associated protein 35 | HUMAN |
| **sp\|P17987\|TCPA_HUMAN** | T-complex protein 1 subunit alpha | HUMAN |
| **sp\|Q13561\|DCTN2_HUMAN** | Dynactin subunit 2 | HUMAN |
| **sp\|Q12926\|ELAV2_HUMAN** | ELAV-like protein 2 | HUMAN |
| **sp\|P50454\|SERPH_HUMAN** | Serpin H1 | HUMAN |
| **sp\|P10809\|CH60_HUMAN** | 60 kDa heat shock protein, mitochondrial | HUMAN |

| | | |
|---|---|---|
| **sp\|P0DMV9\|HS71B_HUMAN** | Heat shock 70 kDa protein 1B | HUMAN |
| **sp\|P33991\|MCM4_HUMAN** | DNA replication licensing factor MCM4 | HUMAN |
| **sp\|P05455\|LA_HUMAN** | Lupus La protein | HUMAN |
| **sp\|Q99832\|TCPH_HUMAN** | T-complex protein 1 subunit eta | HUMAN |
| **sp\|O43175\|SERA_HUMAN** | D-3-phosphoglycerate dehydrogenase | HUMAN |
| **sp\|P35612\|ADDB_HUMAN** | Beta-adducin | HUMAN |
| **sp\|P09651\|ROA1_HUMAN** | Heterogeneous nuclear ribonucleoprotein A1 | HUMAN |
| **sp\|Q86XP3\|DDX42_HUMAN** | ATP-dependent RNA helicase DDX42 | HUMAN |
| **sp\|Q9UN86\|G3BP2_HUMAN** | Ras GTPase-activating protein-binding protein 2 | HUMAN |
| **sp\|Q9UQ80\|PA2G4_HUMAN** | Proliferation-associated protein 2G4 | HUMAN |
| **sp\|Q13177\|PAK2_HUMAN** | Serine/threonine-protein kinase PAK 2 | HUMAN |
| **sp\|O00231\|PSD11_HUMAN** | 26S proteasome non-ATPase regulatory subunit 11 | HUMAN |
| **sp\|P35998\|PRS7_HUMAN** | 26S proteasome regulatory subunit 7 | HUMAN |
| **sp\|O95819\|M4K4_HUMAN** | Mitogen-activated protein kinase kinase kinase kinase 4 | HUMAN |
| **sp\|Q15811\|ITSN1_HUMAN** | Intersectin-1 | HUMAN |
| **sp\|Q16181\|SEPT7_HUMAN** | Septin-7 | HUMAN |
| **sp\|Q9Y266\|NUDC_HUMAN** | Nuclear migration protein nudC | HUMAN |
| **sp\|P00558\|PGK1_HUMAN** | Phosphoglycerate kinase 1 | HUMAN |
| **sp\|P51610\|HCFC1_HUMAN** | Host cell factor 1 | HUMAN |
| **sp\|P04075\|ALDOA_HUMAN** | Fructose-bisphosphate aldolase A | HUMAN |
| **sp\|P04406\|G3P_HUMAN** | Glyceraldehyde-3-phosphate dehydrogenase | HUMAN |
| **sp\|P09429\|HMGB1_HUMAN** | High mobility group protein B1 | HUMAN |
| **sp\|Q9UMS4\|PRP19_HUMAN** | Pre-mRNA-processing factor 19 | HUMAN |
| **sp\|Q13509\|TBB3_HUMAN** | Tubulin beta-3 chain | HUMAN |
| **sp\|P23193\|TCEA1_HUMAN** | Transcription elongation factor A protein 1 | HUMAN |
| **sp\|O75821\|EIF3G_HUMAN** | Eukaryotic translation initiation factor 3 subunit G | HUMAN |
| **sp\|Q14103\|HNRPD_HUMAN** | Heterogeneous nuclear ribonucleoprotein D0 | HUMAN |
| **sp\|P61163\|ACTZ_HUMAN** | Alpha-centractin | HUMAN |
| **sp\|O60749\|SNX2_HUMAN** | Sorting nexin-2 | HUMAN |
| **sp\|P48681\|NEST_HUMAN** | Nestin | HUMAN |
| **sp\|Q9Y6M1\|IF2B2_HUMAN** | Insulin-like growth factor 2 mRNA-binding protein 2 | HUMAN |
| **sp\|P43246\|MSH2_HUMAN** | DNA mismatch repair protein Msh2 | HUMAN |
| **sp\|Q7Z460\|CLAP1_HUMAN** | CLIP-associating protein 1 | HUMAN |
| **sp\|Q8N1F7\|NUP93_HUMAN** | Nuclear pore complex protein Nup93 | HUMAN |
| **sp\|P49591\|SYSC_HUMAN** | Serine--tRNA ligase, cytoplasmic | HUMAN |
| **sp\|P43034\|LIS1_HUMAN** | Platelet-activating factor acetylhydrolase IB subunit beta | HUMAN |
| **sp\|Q99536\|VAT1_HUMAN** | Synaptic vesicle membrane protein VAT-1 homolog | HUMAN |
| **sp\|Q9ULV4\|COR1C_HUMAN** | Coronin-1C | HUMAN |
| **sp\|P28482\|MK01_HUMAN** | Mitogen-activated protein kinase 1 | HUMAN |
| **sp\|Q13347\|EIF3I_HUMAN** | Eukaryotic translation initiation factor 3 subunit I | HUMAN |

| | | |
|---|---|---|
| **sp\|P12277\|KCRB_HUMAN** | Creatine kinase B-type | HUMAN |
| **sp\|Q15637\|SF01_HUMAN** | Splicing factor 1 | HUMAN |
| **sp\|Q92900\|RENT1_HUMAN** | Regulator of nonsense transcripts 1 | HUMAN |
| **sp\|P16949\|STMN1_HUMAN** | Stathmin | HUMAN |
| **sp\|Q15365\|PCBP1_HUMAN** | Poly(rC)-binding protein 1 | HUMAN |
| **sp\|P61011\|SRP54_HUMAN** | Signal recognition particle 54 kDa protein | HUMAN |
| **sp\|P36507\|MP2K2_HUMAN** | Dual specificity mitogen-activated protein kinase kinase 2 | HUMAN |
| **sp\|Q14694\|UBP10_HUMAN** | Ubiquitin carboxyl-terminal hydrolase 10 | HUMAN |
| **sp\|P26196\|DDX6_HUMAN** | Probable ATP-dependent RNA helicase DDX6 | HUMAN |
| **sp\|O15371\|EIF3D_HUMAN** | Eukaryotic translation initiation factor 3 subunit D | HUMAN |
| **sp\|O15372\|EIF3H_HUMAN** | Eukaryotic translation initiation factor 3 subunit H | HUMAN |
| **sp\|Q9UNW9\|NOVA2_HUMAN** | RNA-binding protein Nova-2 | HUMAN |
| **sp\|P08621\|RU17_HUMAN** | U1 small nuclear ribonucleoprotein 70 kDa | HUMAN |
| **sp\|P31689\|DNJA1_HUMAN** | DnaJ homolog subfamily A member 1 | HUMAN |
| **sp\|P38159\|RBMX_HUMAN** | RNA-binding motif protein, X chromosome | HUMAN |
| **sp\|P07910\|HNRPC_HUMAN** | Heterogeneous nuclear ribonucleoproteins C1/C2 | HUMAN |
| **sp\|P08865\|RSSA_HUMAN** | 40S ribosomal protein SA | HUMAN |
| **sp\|O60506\|HNRPQ_HUMAN** | Heterogeneous nuclear ribonucleoprotein Q | HUMAN |
| **sp\|O00267\|SPT5H_HUMAN** | Transcription elongation factor SPT5 | HUMAN |
| **sp\|Q9Y6G9\|DC1L1_HUMAN** | Cytoplasmic dynein 1 light intermediate chain 1 | HUMAN |
| **sp\|Q96GM5\|SMRD1_HUMAN** | SWI/SNF-related matrix-associated actin-dependent regulator of chromatin subfamily D member 1 | HUMAN |
| **sp\|O00425\|IF2B3_HUMAN** | Insulin-like growth factor 2 mRNA-binding protein 3 | HUMAN |
| **sp\|Q92769\|HDAC2_HUMAN** | Histone deacetylase 2 | HUMAN |
| **sp\|Q13838\|DX39B_HUMAN** | Spliceosome RNA helicase DDX39B | HUMAN |
| **sp\|P30419\|NMT1_HUMAN** | Glycylpeptide N-tetradecanoyltransferase 1 | HUMAN |
| **sp\|Q8N1G4\|LRC47_HUMAN** | Leucine-rich repeat-containing protein 47 | HUMAN |
| **sp\|Q15233\|NONO_HUMAN** | Non-POU domain-containing octamer-binding protein | HUMAN |
| **sp\|Q13283\|G3BP1_HUMAN** | Ras GTPase-activating protein-binding protein 1 | HUMAN |
| **sp\|Q9UBC2\|EP15R_HUMAN** | Epidermal growth factor receptor substrate 15-like 1 | HUMAN |
| **sp\|P22234\|PUR6_HUMAN** | Bifunctional phosphoribosylaminoimidazole carboxylase/phosphoribosylaminoimidazole succinocarboxamide synthetase | HUMAN |
| **sp\|O43491\|E41L2_HUMAN** | Band 4.1-like protein 2 | HUMAN |
| **sp\|P61247\|RS3A_HUMAN** | 40S ribosomal protein S3a | HUMAN |
| **sp\|P62136\|PP1A_HUMAN** | Serine/threonine-protein phosphatase PP1-alpha catalytic subunit | HUMAN |
| **sp\|P49736\|MCM2_HUMAN** | DNA replication licensing factor MCM2 | HUMAN |
| **sp\|O14974\|MYPT1_HUMAN** | Protein phosphatase 1 regulatory subunit 12A | HUMAN |

| | | |
|---|---|---|
| **sp\|P62701\|RS4X_HUMAN** | 40S ribosomal protein S4, X isoform | HUMAN |
| **sp\|Q14315\|FLNC_HUMAN** | Filamin-C | HUMAN |
| **sp\|P60842\|IF4A1_HUMAN** | Eukaryotic initiation factor 4A-I | HUMAN |
| **sp\|Q9Y6E2\|5MP1_HUMAN** | eIF5-mimic protein 1 | HUMAN |
| **sp\|P06493\|CDK1_HUMAN** | Cyclin-dependent kinase 1 | HUMAN |
| **sp\|Q9H0B6\|KLC2_HUMAN** | Kinesin light chain 2 | HUMAN |
| **sp\|Q12888\|TP53B_HUMAN** | TP53-binding protein 1 | HUMAN |
| **sp\|P05388\|RLA0_HUMAN** | 60S acidic ribosomal protein P0 | HUMAN |
| **sp\|Q14247\|SRC8_HUMAN** | Src substrate cortactin | HUMAN |
| **sp\|Q15334\|L2GL1_HUMAN** | Lethal(2) giant larvae protein homolog 1 | HUMAN |
| **sp\|O15075\|DCLK1_HUMAN** | Serine/threonine-protein kinase DCLK1 | HUMAN |
| **sp\|Q09028\|RBBP4_HUMAN** | Histone-binding protein RBBP4 | HUMAN |
| **sp\|O43143\|DHX15_HUMAN** | ATP-dependent RNA helicase DHX15 | HUMAN |
| **sp\|O43237\|DC1L2_HUMAN** | Cytoplasmic dynein 1 light intermediate chain 2 | HUMAN |
| **sp\|Q9HCJ6\|VAT1L_HUMAN** | Synaptic vesicle membrane protein VAT-1 homolog-like | HUMAN |
| **sp\|P51991\|ROA3_HUMAN** | Heterogeneous nuclear ribonucleoprotein A3 | HUMAN |
| **sp\|O60884\|DNJA2_HUMAN** | DnaJ homolog subfamily A member 2 | HUMAN |
| **sp\|Q15293\|RCN1_HUMAN** | Reticulocalbin-1 | HUMAN |
| **sp\|P46459\|NSF_HUMAN** | Vesicle-fusing ATPase | HUMAN |
| **sp\|O00170\|AIP_HUMAN** | AH receptor-interacting protein | HUMAN |
| **sp\|Q14498\|RBM39_HUMAN** | RNA-binding protein 39 | HUMAN |
| **sp\|O95747\|OXSR1_HUMAN** | Serine/threonine-protein kinase OSR1 | HUMAN |
| **sp\|Q7Z4V5\|HDGR2_HUMAN** | Hepatoma-derived growth factor-related protein 2 | HUMAN |
| **sp\|P68400\|CSK21_HUMAN** | Casein kinase II subunit alpha | HUMAN |
| **sp\|P07195\|LDHB_HUMAN** | L-lactate dehydrogenase B chain | HUMAN |
| **sp\|Q7L2H7\|EIF3M_HUMAN** | Eukaryotic translation initiation factor 3 subunit M | HUMAN |
| **sp\|P60174\|TPIS_HUMAN** | Triosephosphate isomerase | HUMAN |
| **sp\|Q9UBB4\|ATX10_HUMAN** | Ataxin-10 | HUMAN |
| **sp\|Q00688\|FKBP3_HUMAN** | Peptidyl-prolyl cis-trans isomerase FKBP3 | HUMAN |
| **sp\|Q15020\|SART3_HUMAN** | Squamous cell carcinoma antigen recognized by T-cells 3 | HUMAN |
| **sp\|P62937\|PPIA_HUMAN** | Peptidyl-prolyl cis-trans isomerase A | HUMAN |
| **sp\|P46777\|RL5_HUMAN** | 60S ribosomal protein L5 | HUMAN |
| **sp\|P41091\|IF2G_HUMAN** | Eukaryotic translation initiation factor 2 subunit 3 | HUMAN |
| **sp\|Q13310\|PABP4_HUMAN** | Polyadenylate-binding protein 4 | HUMAN |
| **sp\|P62820\|RAB1A_HUMAN** | Ras-related protein Rab-1A | HUMAN |
| **sp\|P35579\|MYH9_HUMAN** | Myosin-9 | HUMAN |
| **sp\|Q9UKV3\|ACINU_HUMAN** | Apoptotic chromatin condensation inducer in the nucleus | HUMAN |
| **sp\|P55010\|IF5_HUMAN** | Eukaryotic translation initiation factor 5 | HUMAN |
| **sp\|P53618\|COPB_HUMAN** | Coatomer subunit beta | HUMAN |

| | | |
|---|---|---|
| **sp\|Q16630\|CPSF6_HUMAN** | Cleavage and polyadenylation specificity factor subunit 6 | HUMAN |
| **sp\|Q9UMX0\|UBQL1_HUMAN** | Ubiquilin-1 | HUMAN |
| **sp\|Q7KZF4\|SND1_HUMAN** | Staphylococcal nuclease domain-containing protein 1 | HUMAN |
| **sp\|Q8WVV9\|HNRLL_HUMAN** | Heterogeneous nuclear ribonucleoprotein L-like | HUMAN |
| **sp\|O43347\|MSI1H_HUMAN** | RNA-binding protein Musashi homolog 1 | HUMAN |
| **sp\|Q8WWM7\|ATX2L_HUMAN** | Ataxin-2-like protein | HUMAN |
| **sp\|Q96P16\|RPR1A_HUMAN** | Regulation of nuclear pre-mRNA domain-containing protein 1A | HUMAN |
| **sp\|O43242\|PSMD3_HUMAN** | 26S proteasome non-ATPase regulatory subunit 3 | HUMAN |
| **sp\|Q07955\|SRSF1_HUMAN** | Serine/arginine-rich splicing factor 1 | HUMAN |
| **sp\|O60264\|SMCA5_HUMAN** | SWI/SNF-related matrix-associated actin-dependent regulator of chromatin subfamily A member 5 | HUMAN |
| **sp\|O15020\|SPTN2_HUMAN** | Spectrin beta chain, non-erythrocytic 2 | HUMAN |
| **sp\|O00303\|EIF3F_HUMAN** | Eukaryotic translation initiation factor 3 subunit F | HUMAN |
| **sp\|Q9NSD9\|SYFB_HUMAN** | Phenylalanine--tRNA ligase beta subunit | HUMAN |
| **sp\|Q8WX92\|NELFB_HUMAN** | Negative elongation factor B | HUMAN |
| **sp\|P53621\|COPA_HUMAN** | Coatomer subunit alpha | HUMAN |
| **sp\|Q99460\|PSMD1_HUMAN** | 26S proteasome non-ATPase regulatory subunit 1 | HUMAN |
| **sp\|P51178\|PLCD1_HUMAN** | 1-phosphatidylinositol 4,5-bisphosphate phosphodiesterase delta-1 | HUMAN |
| **sp\|Q13557\|KCC2D_HUMAN** | Calcium/calmodulin-dependent protein kinase type II subunit delta | HUMAN |
| **sp\|Q52LJ0\|FA98B_HUMAN** | Protein FAM98B | HUMAN |
| **sp\|Q7L099\|RUFY3_HUMAN** | Protein RUFY3 | HUMAN |
| **sp\|Q9UKA9\|PTBP2_HUMAN** | Polypyrimidine tract-binding protein 2 | HUMAN |
| **sp\|P09661\|RU2A_HUMAN** | U2 small nuclear ribonucleoprotein A' | HUMAN |
| **sp\|P39019\|RS19_HUMAN** | 40S ribosomal protein S19 | HUMAN |
| **sp\|P35611\|ADDA_HUMAN** | Alpha-adducin | HUMAN |
| **sp\|O43809\|CPSF5_HUMAN** | Cleavage and polyadenylation specificity factor subunit 5 | HUMAN |
| **sp\|P13861\|KAP2_HUMAN** | cAMP-dependent protein kinase type II-alpha regulatory subunit | HUMAN |
| **sp\|P31942\|HNRH3_HUMAN** | Heterogeneous nuclear ribonucleoprotein H3 | HUMAN |
| **sp\|O14497\|ARI1A_HUMAN** | AT-rich interactive domain-containing protein 1A | HUMAN |
| **sp\|P35269\|T2FA_HUMAN** | General transcription factor IIF subunit 1 | HUMAN |
| **sp\|P62195\|PRS8_HUMAN** | 26S proteasome regulatory subunit 8 | HUMAN |
| **sp\|Q9UNM6\|PSD13_HUMAN** | 26S proteasome non-ATPase regulatory subunit 13 | HUMAN |
| **sp\|Q14444\|CAPR1_HUMAN** | Caprin-1 | HUMAN |
| **sp\|Q9Y2A7\|NCKP1_HUMAN** | Nck-associated protein 1 | HUMAN |
| **sp\|Q8WWY3\|PRP31_HUMAN** | U4/U6 small nuclear ribonucleoprotein Prp31 | HUMAN |
| **sp\|Q9P2K5\|MYEF2_HUMAN** | Myelin expression factor 2 | HUMAN |

| | | |
|---|---|---|
| **sp\|O96019\|ACL6A_HUMAN** | Actin-like protein 6A | HUMAN |
| **sp\|O60664\|PLIN3_HUMAN** | Perilipin-3 | HUMAN |
| **sp\|P22059\|OSBP1_HUMAN** | Oxysterol-binding protein 1 | HUMAN |
| **sp\|Q9UNH7\|SNX6_HUMAN** | Sorting nexin-6 | HUMAN |
| **sp\|O00429\|DNM1L_HUMAN** | Dynamin-1-like protein | HUMAN |
| **sp\|P48444\|COPD_HUMAN** | Coatomer subunit delta | HUMAN |
| **sp\|Q15417\|CNN3_HUMAN** | Calponin-3 | HUMAN |
| **sp\|P27695\|APEX1_HUMAN** | DNA-(apurinic or apyrimidinic site) endonuclease | HUMAN |
| **sp\|Q9Y4L1\|HYOU1_HUMAN** | Hypoxia up-regulated protein 1 | HUMAN |
| **sp\|Q15907\|RB11B_HUMAN** | Ras-related protein Rab-11B | HUMAN |
| **sp\|O94888\|UBXN7_HUMAN** | UBX domain-containing protein 7 | HUMAN |
| **sp\|Q9Y224\|RTRAF_HUMAN** | RNA transcription, translation and transport factor protein | HUMAN |
| **sp\|P67809\|YBOX1_HUMAN** | Y-box-binding protein 1 | HUMAN |
| **sp\|P23284\|PPIB_HUMAN** | Peptidyl-prolyl cis-trans isomerase B | HUMAN |
| **sp\|O14964\|HGS_HUMAN** | Hepatocyte growth factor-regulated tyrosine kinase substrate | HUMAN |
| **sp\|P18085\|ARF4_HUMAN** | ADP-ribosylation factor 4 | HUMAN |
| **sp\|P55072\|TERA_HUMAN** | Transitional endoplasmic reticulum ATPase | HUMAN |
| **sp\|P63104\|1433Z_HUMAN** | 14-3-3 protein zeta/delta | HUMAN |
| **sp\|P15880\|RS2_HUMAN** | 40S ribosomal protein S2 | HUMAN |
| **sp\|P06748\|NPM_HUMAN** | Nucleophosmin | HUMAN |
| **sp\|P17980\|PRS6A_HUMAN** | 26S proteasome regulatory subunit 6A | HUMAN |
| **sp\|P31150\|GDIA_HUMAN** | Rab GDP dissociation inhibitor alpha | HUMAN |
| **sp\|O75122\|CLAP2_HUMAN** | CLIP-associating protein 2 | HUMAN |
| **sp\|Q5SQI0\|ATAT_HUMAN** | Alpha-tubulin N-acetyltransferase 1 | HUMAN |
| **sp\|Q15008\|PSMD6_HUMAN** | 26S proteasome non-ATPase regulatory subunit 6 | HUMAN |
| **sp\|P20290\|BTF3_HUMAN** | Transcription factor BTF3 | HUMAN |
| **sp\|P49959\|MRE11_HUMAN** | Double-strand break repair protein MRE11 | HUMAN |
| **sp\|P11171\|EPB41_HUMAN** | Protein 4.1 | HUMAN |
| **sp\|P46781\|RS9_HUMAN** | 40S ribosomal protein S9 | HUMAN |
| **sp\|O75937\|DNJC8_HUMAN** | DnaJ homolog subfamily C member 8 | HUMAN |
| **sp\|P29762\|RABP1_HUMAN** | Cellular retinoic acid-binding protein 1 | HUMAN |
| **sp\|P23528\|COF1_HUMAN** | Cofilin-1 | HUMAN |
| **sp\|P26640\|SYVC_HUMAN** | Valine--tRNA ligase | HUMAN |
| **sp\|Q08752\|PPID_HUMAN** | Peptidyl-prolyl cis-trans isomerase D | HUMAN |
| **sp\|Q7L576\|CYFP1_HUMAN** | Cytoplasmic FMR1-interacting protein 1 | HUMAN |
| **sp\|Q9Y2W2\|WBP11_HUMAN** | WW domain-binding protein 11 | HUMAN |
| **sp\|O00151\|PDLI1_HUMAN** | PDZ and LIM domain protein 1 | HUMAN |
| **sp\|P39023\|RL3_HUMAN** | 60S ribosomal protein L3 | HUMAN |
| **sp\|Q86X55\|CARM1_HUMAN** | Histone-arginine methyltransferase CARM1 | HUMAN |
| **sp\|P62249\|RS16_HUMAN** | 40S ribosomal protein S16 | HUMAN |
| **sp\|Q9UI15\|TAGL3_HUMAN** | Transgelin-3 | HUMAN |

| | | |
|---|---|---|
| **sp\|P27694\|RFA1_HUMAN** | Replication protein A 70 kDa DNA-binding subunit | HUMAN |
| **sp\|Q53EL6\|PDCD4_HUMAN** | Programmed cell death protein 4 | HUMAN |
| **sp\|Q15717\|ELAV1_HUMAN** | ELAV-like protein 1 | HUMAN |
| **sp\|Q9H2M9\|RBGPR_HUMAN** | Rab3 GTPase-activating protein non-catalytic subunit | HUMAN |
| **sp\|P07339\|CATD_HUMAN** | Cathepsin D | HUMAN |
| **sp\|O60763\|USO1_HUMAN** | General vesicular transport factor p115 | HUMAN |
| **sp\|O00232\|PSD12_HUMAN** | 26S proteasome non-ATPase regulatory subunit 12 | HUMAN |
| **sp\|P11172\|UMPS_HUMAN** | Uridine 5'-monophosphate synthase | HUMAN |
| **sp\|O75688\|PPM1B_HUMAN** | Protein phosphatase 1B | HUMAN |
| **sp\|P43686\|PRS6B_HUMAN** | 26S proteasome regulatory subunit 6B | HUMAN |
| **sp\|P54727\|RD23B_HUMAN** | UV excision repair protein RAD23 homolog B | HUMAN |
| **sp\|P25685\|DNJB1_HUMAN** | DnaJ homolog subfamily B member 1 | HUMAN |
| **sp\|Q15181\|IPYR_HUMAN** | Inorganic pyrophosphatase | HUMAN |
| **sp\|O76070\|SYUG_HUMAN** | Gamma-synuclein | HUMAN |
| **sp\|P52597\|HNRPF_HUMAN** | Heterogeneous nuclear ribonucleoprotein F | HUMAN |
| **sp\|P62191\|PRS4_HUMAN** | 26S proteasome regulatory subunit 4 | HUMAN |
| **sp\|Q86V81\|THOC4_HUMAN** | THO complex subunit 4 | HUMAN |
| **sp\|Q13596\|SNX1_HUMAN** | Sorting nexin-1 | HUMAN |
| **sp\|P45974\|UBP5_HUMAN** | Ubiquitin carboxyl-terminal hydrolase 5 | HUMAN |
| **sp\|Q06830\|PRDX1_HUMAN** | Peroxiredoxin-1 | HUMAN |
| **sp\|P51148\|RAB5C_HUMAN** | Ras-related protein Rab-5C | HUMAN |
| **sp\|P55209\|NP1L1_HUMAN** | Nucleosome assembly protein 1-like 1 | HUMAN |
| **sp\|Q969G3\|SMCE1_HUMAN** | SWI/SNF-related matrix-associated actin-dependent regulator of chromatin subfamily E member 1 | HUMAN |
| **sp\|P63241\|IF5A1_HUMAN** | Eukaryotic translation initiation factor 5A-1 | HUMAN |
| **sp\|Q9Y3A5\|SBDS_HUMAN** | Ribosome maturation protein SBDS | HUMAN |
| **sp\|P61019\|RAB2A_HUMAN** | Ras-related protein Rab-2A | HUMAN |
| **sp\|O14818\|PSA7_HUMAN** | Proteasome subunit alpha type-7 | HUMAN |
| **sp\|Q13148\|TADBP_HUMAN** | TAR DNA-binding protein 43 | HUMAN |
| **sp\|P08579\|RU2B_HUMAN** | U2 small nuclear ribonucleoprotein B'' | HUMAN |
| **sp\|P26368\|U2AF2_HUMAN** | Splicing factor U2AF 65 kDa subunit | HUMAN |
| **sp\|P06744\|G6PI_HUMAN** | Glucose-6-phosphate isomerase | HUMAN |
| **sp\|P62495\|ERF1_HUMAN** | Eukaryotic peptide chain release factor subunit 1 | HUMAN |
| **sp\|P50502\|F10A1_HUMAN** | Hsc70-interacting protein | HUMAN |
| **sp\|Q7Z4W1\|DCXR_HUMAN** | L-xylulose reductase | HUMAN |
| **sp\|Q16658\|FSCN1_HUMAN** | Fascin | HUMAN |
| **sp\|P60660\|MYL6_HUMAN** | Myosin light polypeptide 6 | HUMAN |
| **sp\|P35221\|CTNA1_HUMAN** | Catenin alpha-1 | HUMAN |
| **sp\|O60271\|JIP4_HUMAN** | C-Jun-amino-terminal kinase-interacting protein 4 | HUMAN |
| **sp\|P29692\|EF1D_HUMAN** | Elongation factor 1-delta | HUMAN |
| **sp\|P43487\|RANG_HUMAN** | Ran-specific GTPase-activating protein | HUMAN |

| | | |
|---|---|---|
| **sp\|Q9UH03\|SEPT3_HUMAN** | Neuronal-specific septin-3 | HUMAN |
| **sp\|Q96CW1\|AP2M1_HUMAN** | AP-2 complex subunit mu | HUMAN |
| **sp\|P35637\|FUS_HUMAN** | RNA-binding protein FUS | HUMAN |
| **sp\|P46060\|RAGP1_HUMAN** | Ran GTPase-activating protein 1 | HUMAN |
| **sp\|Q15056\|IF4H_HUMAN** | Eukaryotic translation initiation factor 4H | HUMAN |
| **sp\|Q99439\|CNN2_HUMAN** | Calponin-2 | HUMAN |
| **sp\|Q01813\|PFKAP_HUMAN** | ATP-dependent 6-phosphofructokinase, platelet type | HUMAN |
| **sp\|Q9NPQ8\|RIC8A_HUMAN** | Synembryn-A | HUMAN |
| **sp\|Q99961\|SH3G1_HUMAN** | Endophilin-A2 | HUMAN |
| **sp\|Q96ST2\|IWS1_HUMAN** | Protein IWS1 homolog | HUMAN |
| **sp\|P61981\|1433G_HUMAN** | 14-3-3 protein gamma | HUMAN |
| **sp\|P51149\|RAB7A_HUMAN** | Ras-related protein Rab-7a | HUMAN |
| **sp\|Q9H1B7\|I2BPL_HUMAN** | Probable E3 ubiquitin-protein ligase IRF2BPL | HUMAN |
| **sp\|Q01844\|EWS_HUMAN** | RNA-binding protein EWS | HUMAN |
| **sp\|Q16543\|CDC37_HUMAN** | Hsp90 co-chaperone Cdc37 | HUMAN |
| **sp\|P62269\|RS18_HUMAN** | 40S ribosomal protein S18 | HUMAN |
| **sp\|Q13492\|PICAL_HUMAN** | Phosphatidylinositol-binding clathrin assembly protein | HUMAN |
| **sp\|P62333\|PRS10_HUMAN** | 26S proteasome regulatory subunit 10B | HUMAN |
| **sp\|Q9Y383\|LC7L2_HUMAN** | Putative RNA-binding protein Luc7-like 2 | HUMAN |
| **sp\|P46379\|BAG6_HUMAN** | Large proline-rich protein BAG6 | HUMAN |
| **sp\|P47756\|CAPZB_HUMAN** | F-actin-capping protein subunit beta | HUMAN |
| **sp\|P38606\|VATA_HUMAN** | V-type proton ATPase catalytic subunit A | HUMAN |
| **sp\|Q9NQG5\|RPR1B_HUMAN** | Regulation of nuclear pre-mRNA domain-containing protein 1B | HUMAN |
| **sp\|Q15428\|SF3A2_HUMAN** | Splicing factor 3A subunit 2 | HUMAN |
| **sp\|P13984\|T2FB_HUMAN** | General transcription factor IIF subunit 2 | HUMAN |
| **sp\|P46782\|RS5_HUMAN** | 40S ribosomal protein S5 | HUMAN |
| **sp\|P08708\|RS17_HUMAN** | 40S ribosomal protein S17 | HUMAN |
| **sp\|Q16401\|PSMD5_HUMAN** | 26S proteasome non-ATPase regulatory subunit 5 | HUMAN |
| **sp\|Q14847\|LASP1_HUMAN** | LIM and SH3 domain protein 1 | HUMAN |
| **sp\|P29966\|MARCS_HUMAN** | Myristoylated alanine-rich C-kinase substrate | HUMAN |
| **sp\|P25398\|RS12_HUMAN** | 40S ribosomal protein S12 | HUMAN |
| **sp\|Q99459\|CDC5L_HUMAN** | Cell division cycle 5-like protein | HUMAN |
| **sp\|Q08170\|SRSF4_HUMAN** | Serine/arginine-rich splicing factor 4 | HUMAN |
| **sp\|P14678\|RSMB_HUMAN** | Small nuclear ribonucleoprotein-associated proteins B and B' | HUMAN |
| **sp\|Q13151\|ROA0_HUMAN** | Heterogeneous nuclear ribonucleoprotein A0 | HUMAN |
| **sp\|P62979\|RS27A_HUMAN** | Ubiquitin-40S ribosomal protein S27a | HUMAN |
| **sp\|P18669\|PGAM1_HUMAN** | Phosphoglycerate mutase 1 | HUMAN |
| **sp\|P26583\|HMGB2_HUMAN** | High mobility group protein B2 | HUMAN |
| **sp\|P17612\|KAPCA_HUMAN** | cAMP-dependent protein kinase catalytic subunit alpha | HUMAN |

| | | |
|---|---|---|
| **sp\|Q9H267\|VP33B_HUMAN** | Vacuolar protein sorting-associated protein 33B | HUMAN |
| **sp\|Q5T8P6\|RBM26_HUMAN** | RNA-binding protein 26 | HUMAN |
| **sp\|P31483\|TIA1_HUMAN** | Cytotoxic granule associated RNA binding protein TIA1 | HUMAN |
| **sp\|Q9GZS3\|SKI8_HUMAN** | SKI8 subunit of superkiller complex protein | HUMAN |
| **sp\|Q9BVA1\|TBB2B_HUMAN** | Tubulin beta-2B chain | HUMAN |
| **sp\|Q8WYA6\|CTBL1_HUMAN** | Beta-catenin-like protein 1 | HUMAN |
| **sp\|P62241\|RS8_HUMAN** | 40S ribosomal protein S8 | HUMAN |
| **sp\|P49756\|RBM25_HUMAN** | RNA-binding protein 25 | HUMAN |
| **sp\|P62826\|RAN_HUMAN** | GTP-binding nuclear protein Ran | HUMAN |
| **sp\|Q99816\|TS101_HUMAN** | Tumor susceptibility gene 101 protein | HUMAN |
| **sp\|P21281\|VATB2_HUMAN** | V-type proton ATPase subunit B, brain isoform | HUMAN |
| **sp\|Q92890\|UFD1_HUMAN** | Ubiquitin recognition factor in ER-associated degradation protein 1 | HUMAN |
| **sp\|Q96K17\|BT3L4_HUMAN** | Transcription factor BTF3 homolog 4 | HUMAN |
| **sp\|Q8WX93\|PALLD_HUMAN** | Palladin | HUMAN |
| **sp\|P08237\|PFKAM_HUMAN** | ATP-dependent 6-phosphofructokinase, muscle type | HUMAN |
| **sp\|Q15785\|TOM34_HUMAN** | Mitochondrial import receptor subunit TOM34 | HUMAN |
| **sp\|P45983\|MK08_HUMAN** | Mitogen-activated protein kinase 8 | HUMAN |
| **sp\|O43684\|BUB3_HUMAN** | Mitotic checkpoint protein BUB3 | HUMAN |
| **sp\|P61221\|ABCE1_HUMAN** | ATP-binding cassette sub-family E member 1 | HUMAN |
| **sp\|Q02878\|RL6_HUMAN** | 60S ribosomal protein L6 | HUMAN |
| **sp\|Q9Y6Y8\|S23IP_HUMAN** | SEC23-interacting protein | HUMAN |
| **sp\|Q07666\|KHDR1_HUMAN** | KH domain-containing, RNA-binding, signal transduction-associated protein 1 | HUMAN |
| **sp\|P62280\|RS11_HUMAN** | 40S ribosomal protein S11 | HUMAN |
| **sp\|Q9UNF1\|MAGD2_HUMAN** | Melanoma-associated antigen D2 | HUMAN |
| **sp\|Q9UMR2\|DD19B_HUMAN** | ATP-dependent RNA helicase DDX19B | HUMAN |
| **sp\|Q13363\|CTBP1_HUMAN** | C-terminal-binding protein 1 | HUMAN |
| **sp\|P20042\|IF2B_HUMAN** | Eukaryotic translation initiation factor 2 subunit 2 | HUMAN |
| **sp\|Q12904\|AIMP1_HUMAN** | Aminoacyl tRNA synthase complex-interacting multifunctional protein 1 | HUMAN |
| **sp\|O60716\|CTND1_HUMAN** | Catenin delta-1 | HUMAN |
| **sp\|Q99729\|ROAA_HUMAN** | Heterogeneous nuclear ribonucleoprotein A/B | HUMAN |
| **sp\|Q9UQE7\|SMC3_HUMAN** | Structural maintenance of chromosomes protein 3 | HUMAN |
| **sp\|P55327\|TPD52_HUMAN** | Tumor protein D52 | HUMAN |
| **sp\|P19105\|ML12A_HUMAN** | Myosin regulatory light chain 12A | HUMAN |
| **sp\|P61601\|NCALD_HUMAN** | Neurocalcin-delta | HUMAN |
| **sp\|Q8N6H7\|ARFG2_HUMAN** | ADP-ribosylation factor GTPase-activating protein 2 | HUMAN |
| **sp\|Q92783\|STAM1_HUMAN** | Signal transducing adapter molecule 1 | HUMAN |
| **sp\|P53999\|TCP4_HUMAN** | Activated RNA polymerase II transcriptional coactivator p15 | HUMAN |

| | | |
|---|---|---|
| **sp\|P28066\|PSA5_HUMAN** | Proteasome subunit alpha type-5 | HUMAN |
| **sp\|Q05682\|CALD1_HUMAN** | Caldesmon | HUMAN |
| **sp\|Q9Y285\|SYFA_HUMAN** | Phenylalanine--tRNA ligase alpha subunit | HUMAN |
| **sp\|Q96DI7\|SNR40_HUMAN** | U5 small nuclear ribonucleoprotein 40 kDa protein | HUMAN |
| **sp\|Q9Y376\|CAB39_HUMAN** | Calcium-binding protein 39 | HUMAN |
| **sp\|P46783\|RS10_HUMAN** | 40S ribosomal protein S10 | HUMAN |
| **sp\|P52907\|CAZA1_HUMAN** | F-actin-capping protein subunit alpha-1 | HUMAN |
| **sp\|P36578\|RL4_HUMAN** | 60S ribosomal protein L4 | HUMAN |
| **sp\|P62633\|CNBP_HUMAN** | CCHC-type zinc finger nucleic acid binding protein | HUMAN |
| **sp\|P54619\|AAKG1_HUMAN** | 5'-AMP-activated protein kinase subunit gamma-1 | HUMAN |
| **sp\|Q9H0C8\|ILKAP_HUMAN** | Integrin-linked kinase-associated serine/threonine phosphatase 2C | HUMAN |
| **sp\|Q4G0F5\|VP26B_HUMAN** | Vacuolar protein sorting-associated protein 26B | HUMAN |
| **sp\|Q86U70\|LDB1_HUMAN** | LIM domain-binding protein 1 | HUMAN |
| **sp\|P51665\|PSMD7_HUMAN** | 26S proteasome non-ATPase regulatory subunit 7 | HUMAN |
| **sp\|O15347\|HMGB3_HUMAN** | High mobility group protein B3 | HUMAN |
| **sp\|P62316\|SMD2_HUMAN** | Small nuclear ribonucleoprotein Sm D2 | HUMAN |
| **sp\|Q86U44\|MTA70_HUMAN** | N6-adenosine-methyltransferase catalytic subunit | HUMAN |
| **sp\|Q9H444\|CHM4B_HUMAN** | Charged multivesicular body protein 4b | HUMAN |
| **sp\|P26373\|RL13_HUMAN** | 60S ribosomal protein L13 | HUMAN |
| **sp\|Q15366\|PCBP2_HUMAN** | Poly(rC)-binding protein 2 | HUMAN |
| **sp\|P15531\|NDKA_HUMAN** | Nucleoside diphosphate kinase A | HUMAN |
| **sp\|P62424\|RL7A_HUMAN** | 60S ribosomal protein L7a | HUMAN |
| **sp\|Q14683\|SMC1A_HUMAN** | Structural maintenance of chromosomes protein 1A | HUMAN |
| **sp\|O43395\|PRPF3_HUMAN** | U4/U6 small nuclear ribonucleoprotein Prp3 | HUMAN |
| **sp\|Q9UBE0\|SAE1_HUMAN** | SUMO-activating enzyme subunit 1 | HUMAN |
| **sp\|O15498\|YKT6_HUMAN** | Synaptobrevin homolog YKT6 | HUMAN |
| **sp\|Q16204\|CCDC6_HUMAN** | Coiled-coil domain-containing protein 6 | HUMAN |
| **sp\|O43301\|HS12A_HUMAN** | Heat shock 70 kDa protein 12A | HUMAN |
| **sp\|O95219\|SNX4_HUMAN** | Sorting nexin-4 | HUMAN |
| **sp\|Q9NYB0\|TE2IP_HUMAN** | Telomeric repeat-binding factor 2-interacting protein 1 | HUMAN |
| **sp\|P67775\|PP2AA_HUMAN** | Serine/threonine-protein phosphatase 2A catalytic subunit alpha isoform | HUMAN |
| **sp\|P29373\|RABP2_HUMAN** | Cellular retinoic acid-binding protein 2 | HUMAN |
| **sp\|P11021\|BIP_HUMAN** | Endoplasmic reticulum chaperone BiP | HUMAN |
| **sp\|Q9P258\|RCC2_HUMAN** | Protein RCC2 | HUMAN |
| **sp\|P63000\|RAC1_HUMAN** | Ras-related C3 botulinum toxin substrate 1 | HUMAN |
| **sp\|Q14978\|NOLC1_HUMAN** | Nucleolar and coiled-body phosphoprotein 1 | HUMAN |
| **sp\|P61964\|WDR5_HUMAN** | WD repeat-containing protein 5 | HUMAN |
| **sp\|Q07960\|RHG01_HUMAN** | Rho GTPase-activating protein 1 | HUMAN |
| **sp\|P35241\|RADI_HUMAN** | Radixin | HUMAN |

| | | |
|---|---|---|
| **sp\|P51812\|KS6A3_HUMAN** | Ribosomal protein S6 kinase alpha-3 | HUMAN |
| **sp\|P18124\|RL7_HUMAN** | 60S ribosomal protein L7 | HUMAN |
| **sp\|O43324\|MCA3_HUMAN** | Eukaryotic translation elongation factor 1 epsilon-1 | HUMAN |
| **sp\|Q13442\|HAP28_HUMAN** | 28 kDa heat- and acid-stable phosphoprotein | HUMAN |
| **sp\|P61106\|RAB14_HUMAN** | Ras-related protein Rab-14 | HUMAN |
| **sp\|P62244\|RS15A_HUMAN** | 40S ribosomal protein S15a | HUMAN |
| **sp\|O43396\|TXNL1_HUMAN** | Thioredoxin-like protein 1 | HUMAN |
| **sp\|P20340\|RAB6A_HUMAN** | Ras-related protein Rab-6A | HUMAN |
| **sp\|Q9UKF6\|CPSF3_HUMAN** | Cleavage and polyadenylation specificity factor subunit 3 | HUMAN |
| **sp\|P62277\|RS13_HUMAN** | 40S ribosomal protein S13 | HUMAN |
| **sp\|P25786\|PSA1_HUMAN** | Proteasome subunit alpha type-1 | HUMAN |
| **sp\|O94903\|PLPHP_HUMAN** | Pyridoxal phosphate homeostasis protein | HUMAN |
| **sp\|P61289\|PSME3_HUMAN** | Proteasome activator complex subunit 3 | HUMAN |
| **sp\|Q99733\|NP1L4_HUMAN** | Nucleosome assembly protein 1-like 4 | HUMAN |
| **sp\|Q92930\|RAB8B_HUMAN** | Ras-related protein Rab-8B | HUMAN |
| **sp\|P62081\|RS7_HUMAN** | 40S ribosomal protein S7 | HUMAN |
| **sp\|Q96M27\|PRRC1_HUMAN** | Protein PRRC1 | HUMAN |
| **sp\|P17677\|NEUM_HUMAN** | Neuromodulin | HUMAN |
| **sp\|Q8N684\|CPSF7_HUMAN** | Cleavage and polyadenylation specificity factor subunit 7 | HUMAN |
| **sp\|P50552\|VASP_HUMAN** | Vasodilator-stimulated phosphoprotein | HUMAN |
| **sp\|P60900\|PSA6_HUMAN** | Proteasome subunit alpha type-6 | HUMAN |
| **sp\|P55036\|PSMD4_HUMAN** | 26S proteasome non-ATPase regulatory subunit 4 | HUMAN |
| **sp\|Q16629\|SRSF7_HUMAN** | Serine/arginine-rich splicing factor 7 | HUMAN |
| **sp\|P07737\|PROF1_HUMAN** | Profilin-1 | HUMAN |
| **sp\|O14929\|HAT1_HUMAN** | Histone acetyltransferase type B catalytic subunit | HUMAN |
| **sp\|P20618\|PSB1_HUMAN** | Proteasome subunit beta type-1 | HUMAN |
| **sp\|O00203\|AP3B1_HUMAN** | AP-3 complex subunit beta-1 | HUMAN |
| **sp\|P39687\|AN32A_HUMAN** | Acidic leucine-rich nuclear phosphoprotein 32 family member A | HUMAN |
| **sp\|Q15555\|MARE2_HUMAN** | Microtubule-associated protein RP/EB family member 2 | HUMAN |
| **sp\|P09012\|SNRPA_HUMAN** | U1 small nuclear ribonucleoprotein A | HUMAN |
| **sp\|P46779\|RL28_HUMAN** | 60S ribosomal protein L28 | HUMAN |
| **sp\|Q13526\|PIN1_HUMAN** | Peptidyl-prolyl cis-trans isomerase NIMA-interacting 1 | HUMAN |
| **sp\|Q14019\|COTL1_HUMAN** | Coactosin-like protein | HUMAN |
| **sp\|P22087\|FBRL_HUMAN** | rRNA 2'-O-methyltransferase fibrillarin | HUMAN |
| **sp\|Q9BVG4\|PBDC1_HUMAN** | Protein PBDC1 | HUMAN |
| **sp\|Q9UN37\|VPS4A_HUMAN** | Vacuolar protein sorting-associated protein 4A | HUMAN |
| **sp\|Q9NP79\|VTA1_HUMAN** | Vacuolar protein sorting-associated protein VTA1 homolog | HUMAN |
| **sp\|P27635\|RL10_HUMAN** | 60S ribosomal protein L10 | HUMAN |

| | | |
|---|---|---|
| **sp\|P62750\|RL23A_HUMAN** | 60S ribosomal protein L23a | HUMAN |
| **sp\|Q99879\|H2B1M_HUMAN** | Histone H2B type 1-M | HUMAN |
| **sp\|P27361\|MK03_HUMAN** | Mitogen-activated protein kinase 3 | HUMAN |
| **sp\|Q9BUF5\|TBB6_HUMAN** | Tubulin beta-6 chain | HUMAN |
| **sp\|P62888\|RL30_HUMAN** | 60S ribosomal protein L30 | HUMAN |
| **sp\|Q07020\|RL18_HUMAN** | 60S ribosomal protein L18 | HUMAN |
| **sp\|P46109\|CRKL_HUMAN** | Crk-like protein | HUMAN |
| **sp\|Q92804\|RBP56_HUMAN** | TATA-binding protein-associated factor 2N | HUMAN |
| **sp\|P62805\|H4_HUMAN** | Histone H4 | HUMAN |
| **sp\|P14625\|ENPL_HUMAN** | Endoplasmin | HUMAN |
| **sp\|P67870\|CSK2B_HUMAN** | Casein kinase II subunit beta | HUMAN |
| **sp\|O00139\|KIF2A_HUMAN** | Kinesin-like protein KIF2A | HUMAN |
| **sp\|Q3YEC7\|RABL6_HUMAN** | Rab-like protein 6 | HUMAN |
| **sp\|Q14320\|FA50A_HUMAN** | Protein FAM50A | HUMAN |
| **sp\|Q9UKM9\|RALY_HUMAN** | RNA-binding protein Raly | HUMAN |
| **sp\|Q99615\|DNJC7_HUMAN** | DnaJ homolog subfamily C member 7 | HUMAN |
| **sp\|P52292\|IMA1_HUMAN** | Importin subunit alpha-1 | HUMAN |
| **sp\|Q9UHD9\|UBQL2_HUMAN** | Ubiquilin-2 | HUMAN |
| **sp\|Q96PU8\|QKI_HUMAN** | KH domain-containing RNA-binding protein QKI | HUMAN |
| **sp\|P49915\|GUAA_HUMAN** | GMP synthase [glutamine-hydrolyzing] | HUMAN |
| **sp\|Q7RTV0\|PHF5A_HUMAN** | PHD finger-like domain-containing protein 5A | HUMAN |
| **sp\|Q9BSD7\|NTPCR_HUMAN** | Cancer-related nucleoside-triphosphatase | HUMAN |
| **sp\|P62306\|RUXF_HUMAN** | Small nuclear ribonucleoprotein F | HUMAN |
| **sp\|Q05086\|UBE3A_HUMAN** | Ubiquitin-protein ligase E3A | HUMAN |
| **sp\|P40429\|RL13A_HUMAN** | 60S ribosomal protein L13a | HUMAN |
| **sp\|Q12972\|PP1R8_HUMAN** | Nuclear inhibitor of protein phosphatase 1 | HUMAN |
| **sp\|P82979\|SARNP_HUMAN** | SAP domain-containing ribonucleoprotein | HUMAN |
| **sp\|Q8WXF1\|PSPC1_HUMAN** | Paraspeckle component 1 | HUMAN |
| **sp\|Q92541\|RTF1_HUMAN** | RNA polymerase-associated protein RTF1 homolog | HUMAN |
| **sp\|P62873\|GBB1_HUMAN** | Guanine nucleotide-binding protein G(I)/G(S)/G(T) subunit beta-1 | HUMAN |
| **sp\|O75475\|PSIP1_HUMAN** | PC4 and SFRS1-interacting protein | HUMAN |
| **sp\|P61586\|RHOA_HUMAN** | Transforming protein RhoA | HUMAN |
| **sp\|Q01130\|SRSF2_HUMAN** | Serine/arginine-rich splicing factor 2 | HUMAN |
| **sp\|P84077\|ARF1_HUMAN** | ADP-ribosylation factor 1 | HUMAN |
| **sp\|Q9UL25\|RAB21_HUMAN** | Ras-related protein Rab-21 | HUMAN |
| **sp\|Q96HC4\|PDLI5_HUMAN** | PDZ and LIM domain protein 5 | HUMAN |
| **sp\|P36969\|GPX4_HUMAN** | Phospholipid hydroperoxide glutathione peroxidase | HUMAN |
| **sp\|Q9UPN6\|SCAF8_HUMAN** | SR-related and CTD-associated factor 8 | HUMAN |
| **sp\|Q14576\|ELAV3_HUMAN** | ELAV-like protein 3 | HUMAN |
| **sp\|P51858\|HDGF_HUMAN** | Hepatoma-derived growth factor | HUMAN |
| **sp\|Q9NR46\|SHLB2_HUMAN** | Endophilin-B2 | HUMAN |

| | | |
|---|---|---|
| **sp\|Q15084\|PDIA6_HUMAN** | Protein disulfide-isomerase A6 | HUMAN |
| **sp\|P19174\|PLCG1_HUMAN** | 1-phosphatidylinositol 4,5-bisphosphate phosphodiesterase gamma-1 | HUMAN |
| **sp\|Q96SI9\|STRBP_HUMAN** | Spermatid perinuclear RNA-binding protein | HUMAN |
| **sp\|O60573\|IF4E2_HUMAN** | Eukaryotic translation initiation factor 4E type 2 | HUMAN |
| **sp\|P32119\|PRDX2_HUMAN** | Peroxiredoxin-2 | HUMAN |
| **sp\|Q00169\|PIPNA_HUMAN** | Phosphatidylinositol transfer protein alpha isoform | HUMAN |
| **sp\|Q16527\|CSRP2_HUMAN** | Cysteine and glycine-rich protein 2 | HUMAN |
| **sp\|Q9NQC3\|RTN4_HUMAN** | Reticulon-4 | HUMAN |
| **sp\|Q01081\|U2AF1_HUMAN** | Splicing factor U2AF 35 kDa subunit | HUMAN |
| **sp\|P30050\|RL12_HUMAN** | 60S ribosomal protein L12 | HUMAN |
| **sp\|P25789\|PSA4_HUMAN** | Proteasome subunit alpha type-4 | HUMAN |
| **sp\|P61020\|RAB5B_HUMAN** | Ras-related protein Rab-5B | HUMAN |
| **sp\|Q9UJW0\|DCTN4_HUMAN** | Dynactin subunit 4 | HUMAN |
| **sp\|Q01105\|SET_HUMAN** | Protein SET | HUMAN |
| **sp\|Q14011\|CIRBP_HUMAN** | Cold-inducible RNA-binding protein | HUMAN |
| **sp\|P62851\|RS25_HUMAN** | 40S ribosomal protein S25 | HUMAN |
| **sp\|Q9H2G2\|SLK_HUMAN** | STE20-like serine/threonine-protein kinase | HUMAN |
| **sp\|P52434\|RPAB3_HUMAN** | DNA-directed RNA polymerases I, II, and III subunit RPABC3 | HUMAN |
| **sp\|P32969\|RL9_HUMAN** | 60S ribosomal protein L9 | HUMAN |
| **sp\|P49721\|PSB2_HUMAN** | Proteasome subunit beta type-2 | HUMAN |
| **sp\|Q6P3W7\|SCYL2_HUMAN** | SCY1-like protein 2 | HUMAN |
| **sp\|Q15436\|SC23A_HUMAN** | Protein transport protein Sec23A | HUMAN |
| **sp\|Q92879\|CELF1_HUMAN** | CUGBP Elav-like family member 1 | HUMAN |
| **sp\|O75436\|VP26A_HUMAN** | Vacuolar protein sorting-associated protein 26A | HUMAN |
| **sp\|Q13045\|FLII_HUMAN** | Protein flightless-1 homolog | HUMAN |
| **sp\|Q9NR31\|SAR1A_HUMAN** | GTP-binding protein SAR1a | HUMAN |
| **sp\|P39748\|FEN1_HUMAN** | Flap endonuclease 1 | HUMAN |
| **sp\|O00505\|IMA4_HUMAN** | Importin subunit alpha-4 | HUMAN |
| **sp\|P09211\|GSTP1_HUMAN** | Glutathione S-transferase P | HUMAN |
| **sp\|P28074\|PSB5_HUMAN** | Proteasome subunit beta type-5 | HUMAN |
| **sp\|P61764\|STXB1_HUMAN** | Syntaxin-binding protein 1 | HUMAN |
| **sp\|O14979\|HNRDL_HUMAN** | Heterogeneous nuclear ribonucleoprotein D-like | HUMAN |
| **sp\|Q2TAY7\|SMU1_HUMAN** | WD40 repeat-containing protein SMU1 | HUMAN |
| **sp\|P07197\|NFM_HUMAN** | Neurofilament medium polypeptide | HUMAN |
| **sp\|P83731\|RL24_HUMAN** | 60S ribosomal protein L24 | HUMAN |
| **sp\|P53990\|IST1_HUMAN** | IST1 homolog | HUMAN |
| **sp\|O94973\|AP2A2_HUMAN** | AP-2 complex subunit alpha-2 | HUMAN |
| **sp\|Q13033\|STRN3_HUMAN** | Striatin-3 | HUMAN |
| **sp\|O14737\|PDCD5_HUMAN** | Programmed cell death protein 5 | HUMAN |
| **sp\|P62753\|RS6_HUMAN** | 40S ribosomal protein S6 | HUMAN |

| | | |
|---|---|---|
| **sp\|Q9BUJ2\|HNRL1_HUMAN** | Heterogeneous nuclear ribonucleoprotein U-like protein 1 | HUMAN |
| **sp\|Q9Y6D9\|MD1L1_HUMAN** | Mitotic spindle assembly checkpoint protein MAD1 | HUMAN |
| **sp\|P18621\|RL17_HUMAN** | 60S ribosomal protein L17 | HUMAN |
| **sp\|Q8WVY7\|UBCP1_HUMAN** | Ubiquitin-like domain-containing CTD phosphatase 1 | HUMAN |
| **sp\|P78406\|RAE1L_HUMAN** | mRNA export factor RAE1 | HUMAN |
| **sp\|O75940\|SPF30_HUMAN** | Survival of motor neuron-related-splicing factor 30 | HUMAN |
| **sp\|Q96PK6\|RBM14_HUMAN** | RNA-binding protein 14 | HUMAN |
| **sp\|Q13547\|HDAC1_HUMAN** | Histone deacetylase 1 | HUMAN |
| **sp\|P24534\|EF1B_HUMAN** | Elongation factor 1-beta | HUMAN |
| **sp\|P61254\|RL26_HUMAN** | 60S ribosomal protein L26 | HUMAN |
| **sp\|P25787\|PSA2_HUMAN** | Proteasome subunit alpha type-2 | HUMAN |
| **sp\|Q13451\|FKBP5_HUMAN** | Peptidyl-prolyl cis-trans isomerase FKBP5 | HUMAN |
| **sp\|Q9NUP9\|LIN7C_HUMAN** | Protein lin-7 homolog C | HUMAN |
| **sp\|Q96CT7\|CC124_HUMAN** | Coiled-coil domain-containing protein 124 | HUMAN |
| **sp\|P61026\|RAB10_HUMAN** | Ras-related protein Rab-10 | HUMAN |
| **sp\|Q9NZZ3\|CHMP5_HUMAN** | Charged multivesicular body protein 5 | HUMAN |
| **sp\|Q96MZ0\|GD1L1_HUMAN** | Ganglioside-induced differentiation-associated protein 1-like 1 | HUMAN |
| **sp\|P51513\|NOVA1_HUMAN** | RNA-binding protein Nova-1 | HUMAN |
| **sp\|P25705\|ATPA_HUMAN** | ATP synthase subunit alpha, mitochondrial | HUMAN |
| **sp\|Q9UNS2\|CSN3_HUMAN** | COP9 signalosome complex subunit 3 | HUMAN |
| **sp\|O43776\|SYNC_HUMAN** | Asparagine--tRNA ligase, cytoplasmic | HUMAN |
| **sp\|Q8IYB3\|SRRM1_HUMAN** | Serine/arginine repetitive matrix protein 1 | HUMAN |
| **sp\|P31946\|1433B_HUMAN** | 14-3-3 protein beta/alpha | HUMAN |
| **sp\|P48556\|PSMD8_HUMAN** | 26S proteasome non-ATPase regulatory subunit 8 | HUMAN |
| **sp\|Q9H0D6\|XRN2_HUMAN** | 5'-3' exoribonuclease 2 | HUMAN |
| **sp\|Q04917\|1433F_HUMAN** | 14-3-3 protein eta | HUMAN |
| **sp\|E9PAV3\|NACAM_HUMAN** | Nascent polypeptide-associated complex subunit alpha, muscle-specific form | HUMAN |
| **sp\|Q01518\|CAP1_HUMAN** | Adenylyl cyclase-associated protein 1 | HUMAN |
| **sp\|Q96I25\|SPF45_HUMAN** | Splicing factor 45 | HUMAN |
| **sp\|Q04726\|TLE3_HUMAN** | Transducin-like enhancer protein 3 | HUMAN |
| **sp\|P27816\|MAP4_HUMAN** | Microtubule-associated protein 4 | HUMAN |
| **sp\|P49770\|EI2BB_HUMAN** | Translation initiation factor eIF-2B subunit beta | HUMAN |
| **sp\|P61160\|ARP2_HUMAN** | Actin-related protein 2 | HUMAN |
| **sp\|P50897\|PPT1_HUMAN** | Palmitoyl-protein thioesterase 1 | HUMAN |
| **sp\|O43399\|TPD54_HUMAN** | Tumor protein D54 | HUMAN |
| **sp\|Q13242\|SRSF9_HUMAN** | Serine/arginine-rich splicing factor 9 | HUMAN |
| **sp\|O60869\|EDF1_HUMAN** | Endothelial differentiation-related factor 1 | HUMAN |
| **sp\|P62314\|SMD1_HUMAN** | Small nuclear ribonucleoprotein Sm D1 | HUMAN |
| **sp\|P27348\|1433T_HUMAN** | 14-3-3 protein theta | HUMAN |

| | | |
|---|---|---|
| **sp\|P62906\|RL10A_HUMAN** | 60S ribosomal protein L10a | HUMAN |
| **sp\|Q92530\|PSMF1_HUMAN** | Proteasome inhibitor PI31 subunit | HUMAN |
| **sp\|O94819\|KBTBB_HUMAN** | Kelch repeat and BTB domain-containing protein 11 | HUMAN |
| **sp\|Q9NRV9\|HEBP1_HUMAN** | Heme-binding protein 1 | HUMAN |
| **sp\|Q02750\|MP2K1_HUMAN** | Dual specificity mitogen-activated protein kinase kinase 1 | HUMAN |
| **sp\|P25788\|PSA3_HUMAN** | Proteasome subunit alpha type-3 | HUMAN |
| **sp\|Q7Z739\|YTHD3_HUMAN** | YTH domain-containing family protein 3 | HUMAN |
| **sp\|Q13418\|ILK_HUMAN** | Integrin-linked protein kinase | HUMAN |
| **sp\|P0DP25\|CALM3_HUMAN** | Calmodulin-3 | HUMAN |
| **sp\|P62263\|RS14_HUMAN** | 40S ribosomal protein S14 | HUMAN |
| **sp\|Q15814\|TBCC_HUMAN** | Tubulin-specific chaperone C | HUMAN |
| **sp\|P15927\|RFA2_HUMAN** | Replication protein A 32 kDa subunit | HUMAN |
| **sp\|P62829\|RL23_HUMAN** | 60S ribosomal protein L23 | HUMAN |
| **sp\|Q9UIV1\|CNOT7_HUMAN** | CCR4-NOT transcription complex subunit 7 | HUMAN |
| **sp\|P54577\|SYYC_HUMAN** | Tyrosine--tRNA ligase, cytoplasmic | HUMAN |
| **sp\|P55735\|SEC13_HUMAN** | Protein SEC13 homolog | HUMAN |
| **sp\|Q9Y5K5\|UCHL5_HUMAN** | Ubiquitin carboxyl-terminal hydrolase isozyme L5 | HUMAN |
| **sp\|Q9Y3S1\|WNK2_HUMAN** | Serine/threonine-protein kinase WNK2 | HUMAN |
| **sp\|P28070\|PSB4_HUMAN** | Proteasome subunit beta type-4 | HUMAN |
| **sp\|Q86YP4\|P66A_HUMAN** | Transcriptional repressor p66-alpha | HUMAN |
| **sp\|O00487\|PSDE_HUMAN** | 26S proteasome non-ATPase regulatory subunit 14 | HUMAN |
| **sp\|P49321\|NASP_HUMAN** | Nuclear autoantigenic sperm protein | HUMAN |
| **sp\|P41227\|NAA10_HUMAN** | N-alpha-acetyltransferase 10 | HUMAN |
| **sp\|O95433\|AHSA1_HUMAN** | Activator of 90 kDa heat shock protein ATPase homolog 1 | HUMAN |
| **sp\|P62841\|RS15_HUMAN** | 40S ribosomal protein S15 | HUMAN |
| **sp\|Q9Y2T2\|AP3M1_HUMAN** | AP-3 complex subunit mu-1 | HUMAN |
| **sp\|Q9Y3A3\|PHOCN_HUMAN** | MOB-like protein phocein | HUMAN |
| **sp\|Q9UBQ5\|EIF3K_HUMAN** | Eukaryotic translation initiation factor 3 subunit K | HUMAN |
| **sp\|P10636\|TAU_HUMAN** | Microtubule-associated protein tau | HUMAN |
| **sp\|Q9BWD1\|THIC_HUMAN** | Acetyl-CoA acetyltransferase, cytosolic | HUMAN |
| **sp\|P49419\|AL7A1_HUMAN** | Alpha-aminoadipic semialdehyde dehydrogenase | HUMAN |
| **sp\|P84085\|ARF5_HUMAN** | ADP-ribosylation factor 5 | HUMAN |
| **sp\|P19388\|RPAB1_HUMAN** | DNA-directed RNA polymerases I, II, and III subunit RPABC1 | HUMAN |
| **sp\|Q9H4L7\|SMRCD_HUMAN** | SWI/SNF-related matrix-associated actin-dependent regulator of chromatin subfamily A containing DEAD/H box 1 | HUMAN |
| **sp\|Q9NYF8\|BCLF1_HUMAN** | Bcl-2-associated transcription factor 1 | HUMAN |
| **sp\|Q9UEE9\|CFDP1_HUMAN** | Craniofacial development protein 1 | HUMAN |
| **sp\|Q7Z3B4\|NUP54_HUMAN** | Nucleoporin p54 | HUMAN |
| **sp\|Q14141\|SEPT6_HUMAN** | Septin-6 | HUMAN |

| | | |
|---|---|---|
| **sp\|P07196\|NFL_HUMAN** | Neurofilament light polypeptide | HUMAN |
| **sp\|Q5SSJ5\|HP1B3_HUMAN** | Heterochromatin protein 1-binding protein 3 | HUMAN |
| **sp\|P61088\|UBE2N_HUMAN** | Ubiquitin-conjugating enzyme E2 N | HUMAN |
| **sp\|Q96G61\|NUD11_HUMAN** | Diphosphoinositol polyphosphate phosphohydrolase 3-beta | HUMAN |
| **sp\|Q9UJU6\|DBNL_HUMAN** | Drebrin-like protein | HUMAN |
| **sp\|Q96BM9\|ARL8A_HUMAN** | ADP-ribosylation factor-like protein 8A | HUMAN |
| **sp\|Q9BQ61\|TRIR_HUMAN** | Telomerase RNA component interacting RNase | HUMAN |
| **sp\|P55795\|HNRH2_HUMAN** | Heterogeneous nuclear ribonucleoprotein H2 | HUMAN |
| **sp\|Q9C040\|TRIM2_HUMAN** | Tripartite motif-containing protein 2 | HUMAN |
| **sp\|P63173\|RL38_HUMAN** | 60S ribosomal protein L38 | HUMAN |
| **sp\|Q14677\|EPN4_HUMAN** | Clathrin interactor 1 | HUMAN |
| **sp\|P67936\|TPM4_HUMAN** | Tropomyosin alpha-4 chain | HUMAN |
| **sp\|P55769\|NH2L1_HUMAN** | NHP2-like protein 1 | HUMAN |
| **sp\|Q9P0L2\|MARK1_HUMAN** | Serine/threonine-protein kinase MARK1 | HUMAN |
| **sp\|Q5EBL4\|RIPL1_HUMAN** | RILP-like protein 1 | HUMAN |
| **sp\|Q9HD42\|CHM1A_HUMAN** | Charged multivesicular body protein 1a | HUMAN |
| **sp\|Q9UGI8\|TES_HUMAN** | Testin | HUMAN |
| **sp\|Q15370\|ELOB_HUMAN** | Elongin-B | HUMAN |
| **sp\|O43633\|CHM2A_HUMAN** | Charged multivesicular body protein 2a | HUMAN |
| **sp\|P84103\|SRSF3_HUMAN** | Serine/arginine-rich splicing factor 3 | HUMAN |
| **sp\|O75391\|SPAG7_HUMAN** | Sperm-associated antigen 7 | HUMAN |
| **sp\|Q92600\|CNOT9_HUMAN** | CCR4-NOT transcription complex subunit 9 | HUMAN |
| **sp\|Q13155\|AIMP2_HUMAN** | Aminoacyl tRNA synthase complex-interacting multifunctional protein 2 | HUMAN |
| **sp\|P60866\|RS20_HUMAN** | 40S ribosomal protein S20 | HUMAN |
| **sp\|Q15942\|ZYX_HUMAN** | Zyxin | HUMAN |
| **sp\|P36405\|ARL3_HUMAN** | ADP-ribosylation factor-like protein 3 | HUMAN |
| **sp\|P04792\|HSPB1_HUMAN** | Heat shock protein beta-1 | HUMAN |
| **sp\|P61353\|RL27_HUMAN** | 60S ribosomal protein L27 | HUMAN |
| **sp\|P16401\|H15_HUMAN** | Histone H1.5 | HUMAN |
| **sp\|P33240\|CSTF2_HUMAN** | Cleavage stimulation factor subunit 2 | HUMAN |
| **sp\|Q9Y295\|DRG1_HUMAN** | Developmentally-regulated GTP-binding protein 1 | HUMAN |
| **sp\|O76094\|SRP72_HUMAN** | Signal recognition particle subunit SRP72 | HUMAN |
| **sp\|Q9Y3B4\|SF3B6_HUMAN** | Splicing factor 3B subunit 6 | HUMAN |
| **sp\|Q9UBQ0\|VPS29_HUMAN** | Vacuolar protein sorting-associated protein 29 | HUMAN |
| **sp\|Q9Y3F4\|STRAP_HUMAN** | Serine-threonine kinase receptor-associated protein | HUMAN |
| **sp\|P20339\|RAB5A_HUMAN** | Ras-related protein Rab-5A | HUMAN |
| **sp\|P62917\|RL8_HUMAN** | 60S ribosomal protein L8 | HUMAN |
| **sp\|P37840\|SYUA_HUMAN** | Alpha-synuclein | HUMAN |
| **sp\|O95400\|CD2B2_HUMAN** | CD2 antigen cytoplasmic tail-binding protein 2 | HUMAN |
| **sp\|Q92522\|H1X_HUMAN** | Histone H1.10 | HUMAN |

| | | |
|---|---|---|
| **sp\|Q9H8Y8\|GORS2_HUMAN** | Golgi reassembly-stacking protein 2 | HUMAN |
| **sp\|O00422\|SAP18_HUMAN** | Histone deacetylase complex subunit SAP18 | HUMAN |
| **sp\|Q12792\|TWF1_HUMAN** | Twinfilin-1 | HUMAN |
| **sp\|P35658\|NU214_HUMAN** | Nuclear pore complex protein Nup214 | HUMAN |
| **sp\|P04350\|TBB4A_HUMAN** | Tubulin beta-4A chain | HUMAN |
| **sp\|Q9BWW4\|SSBP3_HUMAN** | Single-stranded DNA-binding protein 3 | HUMAN |
| **sp\|P62899\|RL31_HUMAN** | 60S ribosomal protein L31 | HUMAN |
| **sp\|Q9Y570\|PPME1_HUMAN** | Protein phosphatase methylesterase 1 | HUMAN |
| **sp\|P98175\|RBM10_HUMAN** | RNA-binding protein 10 | HUMAN |
| **sp\|P14314\|GLU2B_HUMAN** | Glucosidase 2 subunit beta | HUMAN |
| **sp\|P62318\|SMD3_HUMAN** | Small nuclear ribonucleoprotein Sm D3 | HUMAN |
| **sp\|O14773\|TPP1_HUMAN** | Tripeptidyl-peptidase 1 | HUMAN |
| **sp\|Q9Y2Z0\|SGT1_HUMAN** | Protein SGT1 homolog | HUMAN |
| **sp\|Q9H8S9\|MOB1A_HUMAN** | MOB kinase activator 1A | HUMAN |
| **sp\|Q71UM5\|RS27L_HUMAN** | 40S ribosomal protein S27-like | HUMAN |
| **sp\|Q13642\|FHL1_HUMAN** | Four and a half LIM domains protein 1 | HUMAN |
| **sp\|P37108\|SRP14_HUMAN** | Signal recognition particle 14 kDa protein | HUMAN |
| **sp\|P40222\|TXLNA_HUMAN** | Alpha-taxilin | HUMAN |
| **sp\|Q71UI9\|H2AV_HUMAN** | Histone H2A.V | HUMAN |
| **sp\|P59998\|ARPC4_HUMAN** | Actin-related protein 2/3 complex subunit 4 | HUMAN |
| **sp\|Q9BT78\|CSN4_HUMAN** | COP9 signalosome complex subunit 4 | HUMAN |
| **sp\|P60953\|CDC42_HUMAN** | Cell division control protein 42 homolog | HUMAN |
| **sp\|O15160\|RPAC1_HUMAN** | DNA-directed RNA polymerases I and III subunit RPAC1 | HUMAN |
| **sp\|O75340\|PDCD6_HUMAN** | Programmed cell death protein 6 | HUMAN |
| **sp\|Q86U42\|PABP2_HUMAN** | Polyadenylate-binding protein 2 | HUMAN |
| **sp\|P47813\|IF1AX_HUMAN** | Eukaryotic translation initiation factor 1A, X-chromosomal | HUMAN |
| **sp\|P20336\|RAB3A_HUMAN** | Ras-related protein Rab-3A | HUMAN |
| **sp\|P62266\|RS23_HUMAN** | 40S ribosomal protein S23 | HUMAN |
| **sp\|Q9P013\|CWC15_HUMAN** | Spliceosome-associated protein CWC15 homolog | HUMAN |
| **sp\|Q96KB5\|TOPK_HUMAN** | Lymphokine-activated killer T-cell-originated protein kinase | HUMAN |
| **sp\|Q8IZP0\|ABI1_HUMAN** | Abl interactor 1 | HUMAN |
| **sp\|Q9UEY8\|ADDG_HUMAN** | Gamma-adducin | HUMAN |
| **sp\|Q96A72\|MGN2_HUMAN** | Protein mago nashi homolog 2 | HUMAN |
| **sp\|Q9BY77\|PDIP3_HUMAN** | Polymerase delta-interacting protein 3 | HUMAN |
| **sp\|Q9C0C9\|UBE2O_HUMAN** | (E3-independent) E2 ubiquitin-conjugating enzyme | HUMAN |
| **sp\|P49458\|SRP09_HUMAN** | Signal recognition particle 9 kDa protein | HUMAN |
| **sp\|Q9Y5X3\|SNX5_HUMAN** | Sorting nexin-5 | HUMAN |
| **sp\|P23258\|TBG1_HUMAN** | Tubulin gamma-1 chain | HUMAN |
| **sp\|P62913\|RL11_HUMAN** | 60S ribosomal protein L11 | HUMAN |
| **sp\|Q00535\|CDK5_HUMAN** | Cyclin-dependent kinase 5 | HUMAN |

| | | |
|---|---|---|
| **sp\|Q53GS9\|SNUT2_HUMAN** | U4/U6.U5 tri-snRNP-associated protein 2 | HUMAN |
| **sp\|Q96FJ2\|DYL2_HUMAN** | Dynein light chain 2, cytoplasmic | HUMAN |
| **sp\|Q12824\|SNF5_HUMAN** | SWI/SNF-related matrix-associated actin-dependent regulator of chromatin subfamily B member 1 | HUMAN |
| **sp\|Q15059\|BRD3_HUMAN** | Bromodomain-containing protein 3 | HUMAN |
| **sp\|Q8N1G2\|CMTR1_HUMAN** | Cap-specific mRNA (nucleoside-2'-O-)-methyltransferase 1 | HUMAN |
| **sp\|Q86Y82\|STX12_HUMAN** | Syntaxin-12 | HUMAN |
| **sp\|P36543\|VATE1_HUMAN** | V-type proton ATPase subunit E 1 | HUMAN |
| **sp\|P19387\|RPB3_HUMAN** | DNA-directed RNA polymerase II subunit RPB3 | HUMAN |
| **sp\|Q99436\|PSB7_HUMAN** | Proteasome subunit beta type-7 | HUMAN |
| **sp\|Q6P996\|PDXD1_HUMAN** | Pyridoxal-dependent decarboxylase domain-containing protein 1 | HUMAN |
| **sp\|P18077\|RL35A_HUMAN** | 60S ribosomal protein L35a | HUMAN |
| **sp\|P63220\|RS21_HUMAN** | 40S ribosomal protein S21 | HUMAN |
| **sp\|P51116\|FXR2_HUMAN** | RNA-binding protein FXR2 | HUMAN |
| **sp\|Q04323\|UBXN1_HUMAN** | UBX domain-containing protein 1 | HUMAN |
| **sp\|Q9H115\|SNAB_HUMAN** | Beta-soluble NSF attachment protein | HUMAN |
| **sp\|P30041\|PRDX6_HUMAN** | Peroxiredoxin-6 | HUMAN |
| **sp\|P09936\|UCHL1_HUMAN** | Ubiquitin carboxyl-terminal hydrolase isozyme L1 | HUMAN |
| **sp\|O60493\|SNX3_HUMAN** | Sorting nexin-3 | HUMAN |
| **sp\|P49841\|GSK3B_HUMAN** | Glycogen synthase kinase-3 beta | HUMAN |
| **sp\|Q9UQ16\|DYN3_HUMAN** | Dynamin-3 | HUMAN |
| **sp\|O43447\|PPIH_HUMAN** | Peptidyl-prolyl cis-trans isomerase H | HUMAN |
| **sp\|Q02543\|RL18A_HUMAN** | 60S ribosomal protein L18a | HUMAN |
| **sp\|O60828\|PQBP1_HUMAN** | Polyglutamine-binding protein 1 | HUMAN |
| **sp\|P52594\|AGFG1_HUMAN** | Arf-GAP domain and FG repeat-containing protein 1 | HUMAN |
| **sp\|Q9Y5Y2\|NUBP2_HUMAN** | Cytosolic Fe-S cluster assembly factor NUBP2 | HUMAN |
| **sp\|Q92905\|CSN5_HUMAN** | COP9 signalosome complex subunit 5 | HUMAN |
| **sp\|O75534\|CSDE1_HUMAN** | Cold shock domain-containing protein E1 | HUMAN |
| **sp\|Q7Z6K5\|ARPIN_HUMAN** | Arpin | HUMAN |
| **sp\|O75934\|SPF27_HUMAN** | Pre-mRNA-splicing factor SPF27 | HUMAN |
| **sp\|Q9ULJ6\|ZMIZ1_HUMAN** | Zinc finger MIZ domain-containing protein 1 | HUMAN |
| **sp\|Q9BUL8\|PDC10_HUMAN** | Programmed cell death protein 10 | HUMAN |
| **sp\|Q7L775\|EPMIP_HUMAN** | EPM2A-interacting protein 1 | HUMAN |
| **sp\|O43670\|ZN207_HUMAN** | BUB3-interacting and GLEBS motif-containing protein ZNF207 | HUMAN |
| **sp\|Q9Y371\|SHLB1_HUMAN** | Endophilin-B1 | HUMAN |
| **sp\|P09496\|CLCA_HUMAN** | Clathrin light chain A | HUMAN |
| **sp\|P46940\|IQGA1_HUMAN** | Ras GTPase-activating-like protein IQGAP1 | HUMAN |
| **sp\|Q9NP97\|DLRB1_HUMAN** | Dynein light chain roadblock-type 1 | HUMAN |
| **sp\|Q09161\|NCBP1_HUMAN** | Nuclear cap-binding protein subunit 1 | HUMAN |

| | | |
|---|---|---|
| **sp\|Q9NZN8\|CNOT2_HUMAN** | CCR4-NOT transcription complex subunit 2 | HUMAN |
| **sp\|Q15102\|PA1B3_HUMAN** | Platelet-activating factor acetylhydrolase IB subunit alpha1 | HUMAN |
| **sp\|Q9Y608\|LRRF2_HUMAN** | Leucine-rich repeat flightless-interacting protein 2 | HUMAN |
| **sp\|O00233\|PSMD9_HUMAN** | 26S proteasome non-ATPase regulatory subunit 9 | HUMAN |
| **sp\|P15311\|EZRI_HUMAN** | Ezrin | HUMAN |
| **sp\|Q9GZZ1\|NAA50_HUMAN** | N-alpha-acetyltransferase 50 | HUMAN |
| **sp\|Q5T0N5\|FBP1L_HUMAN** | Formin-binding protein 1-like | HUMAN |
| **sp\|P62847\|RS24_HUMAN** | 40S ribosomal protein S24 | HUMAN |
| **sp\|P40616\|ARL1_HUMAN** | ADP-ribosylation factor-like protein 1 | HUMAN |
| **sp\|Q9NZ32\|ARP10_HUMAN** | Actin-related protein 10 | HUMAN |
| **sp\|O75935\|DCTN3_HUMAN** | Dynactin subunit 3 | HUMAN |
| **sp\|P57723\|PCBP4_HUMAN** | Poly(rC)-binding protein 4 | HUMAN |
| **sp\|Q9Y5A9\|YTHD2_HUMAN** | YTH domain-containing family protein 2 | HUMAN |
| **sp\|P48729\|KC1A_HUMAN** | Casein kinase I isoform alpha | HUMAN |
| **sp\|P49418\|AMPH_HUMAN** | Amphiphysin | HUMAN |
| **sp\|Q15369\|ELOC_HUMAN** | Elongin-C | HUMAN |
| **sp\|Q9HD15\|SRA1_HUMAN** | Steroid receptor RNA activator 1 | HUMAN |
| **sp\|P54578\|UBP14_HUMAN** | Ubiquitin carboxyl-terminal hydrolase 14 | HUMAN |
| **sp\|O14787\|TNPO2_HUMAN** | Transportin-2 | HUMAN |
| **sp\|P19784\|CSK22_HUMAN** | Casein kinase II subunit alpha' | HUMAN |
| **sp\|P11441\|UBL4A_HUMAN** | Ubiquitin-like protein 4A | HUMAN |
| **sp\|P07237\|PDIA1_HUMAN** | Protein disulfide-isomerase | HUMAN |
| **sp\|Q9UHV9\|PFD2_HUMAN** | Prefoldin subunit 2 | HUMAN |
| **sp\|Q7L5N1\|CSN6_HUMAN** | COP9 signalosome complex subunit 6 | HUMAN |
| **sp\|P51452\|DUS3_HUMAN** | Dual specificity protein phosphatase 3 | HUMAN |
| **sp\|Q92599\|SEPT8_HUMAN** | Septin-8 | HUMAN |
| **sp\|P62330\|ARF6_HUMAN** | ADP-ribosylation factor 6 | HUMAN |
| **sp\|Q9BWJ5\|SF3B5_HUMAN** | Splicing factor 3B subunit 5 | HUMAN |
| **sp\|P46776\|RL27A_HUMAN** | 60S ribosomal protein L27a | HUMAN |
| **sp\|Q03426\|KIME_HUMAN** | Mevalonate kinase | HUMAN |
| **sp\|Q6UN15\|FIP1_HUMAN** | Pre-mRNA 3'-end-processing factor FIP1 | HUMAN |
| **sp\|O15240\|VGF_HUMAN** | Neurosecretory protein VGF | HUMAN |
| **sp\|Q9H910\|JUPI2_HUMAN** | Jupiter microtubule associated homolog 2 | HUMAN |
| **sp\|Q53GG5\|PDLI3_HUMAN** | PDZ and LIM domain protein 3 | HUMAN |
| **sp\|Q9Y3Y2\|CHTOP_HUMAN** | Chromatin target of PRMT1 protein | HUMAN |
| **sp\|Q16799\|RTN1_HUMAN** | Reticulon-1 | HUMAN |
| **sp\|P63172\|DYLT1_HUMAN** | Dynein light chain Tctex-type 1 | HUMAN |
| **sp\|P62857\|RS28_HUMAN** | 40S ribosomal protein S28 | HUMAN |
| **sp\|P50914\|RL14_HUMAN** | 60S ribosomal protein L14 | HUMAN |
| **sp\|P28072\|PSB6_HUMAN** | Proteasome subunit beta type-6 | HUMAN |
| **sp\|P09234\|RU1C_HUMAN** | U1 small nuclear ribonucleoprotein C | HUMAN |
| **sp\|O14994\|SYN3_HUMAN** | Synapsin-3 | HUMAN |

| | | |
|---|---|---|
| **sp\|Q96DH6\|MSI2H_HUMAN** | RNA-binding protein Musashi homolog 2 | HUMAN |
| **sp\|Q8WXI9\|P66B_HUMAN** | Transcriptional repressor p66-beta | HUMAN |
| **sp\|Q99426\|TBCB_HUMAN** | Tubulin-folding cofactor B | HUMAN |
| **sp\|Q8IZQ5\|SELH_HUMAN** | Selenoprotein H | HUMAN |
| **sp\|O75175\|CNOT3_HUMAN** | CCR4-NOT transcription complex subunit 3 | HUMAN |
| **sp\|P49207\|RL34_HUMAN** | 60S ribosomal protein L34 | HUMAN |
| **sp\|P63279\|UBC9_HUMAN** | SUMO-conjugating enzyme UBC9 | HUMAN |
| **sp\|P62273\|RS29_HUMAN** | 40S ribosomal protein S29 | HUMAN |
| **sp\|Q9BRP8\|PYM1_HUMAN** | Partner of Y14 and mago | HUMAN |
| **sp\|A6ZKI3\|RTL8C_HUMAN** | Retrotransposon Gag-like protein 8C | HUMAN |
| **sp\|Q8WW12\|PCNP_HUMAN** | PEST proteolytic signal-containing nuclear protein | HUMAN |
| **sp\|O43660\|PLRG1_HUMAN** | Pleiotropic regulator 1 | HUMAN |
| **sp\|Q15427\|SF3B4_HUMAN** | Splicing factor 3B subunit 4 | HUMAN |
| **sp\|P62308\|RUXG_HUMAN** | Small nuclear ribonucleoprotein G | HUMAN |
| **sp\|Q96GY0\|ZC21A_HUMAN** | Zinc finger C2HC domain-containing protein 1A | HUMAN |
| **sp\|Q9P0L0\|VAPA_HUMAN** | Vesicle-associated membrane protein-associated protein A | HUMAN |
| **sp\|Q15691\|MARE1_HUMAN** | Microtubule-associated protein RP/EB family member 1 | HUMAN |
| **sp\|Q9BVM2\|DPCD_HUMAN** | Protein DPCD | HUMAN |
| **sp\|Q13573\|SNW1_HUMAN** | SNW domain-containing protein 1 | HUMAN |
| **sp\|P18615\|NELFE_HUMAN** | Negative elongation factor E | HUMAN |
| **sp\|P18754\|RCC1_HUMAN** | Regulator of chromosome condensation | HUMAN |
| **sp\|P62995\|TRA2B_HUMAN** | Transformer-2 protein homolog beta | HUMAN |
| **sp\|Q9NWS0\|PIHD1_HUMAN** | PIH1 domain-containing protein 1 | HUMAN |
| **sp\|Q8WXX5\|DNJC9_HUMAN** | DnaJ homolog subfamily C member 9 | HUMAN |
| **sp\|Q71DI3\|H32_HUMAN** | Histone H3.2 | HUMAN |
| **sp\|Q8IYD1\|ERF3B_HUMAN** | Eukaryotic peptide chain release factor GTP-binding subunit ERF3B | HUMAN |
| **sp\|Q9BRG1\|VPS25_HUMAN** | Vacuolar protein-sorting-associated protein 25 | HUMAN |
| **sp\|P61081\|UBC12_HUMAN** | NEDD8-conjugating enzyme Ubc12 | HUMAN |
| **sp\|P46937\|YAP1_HUMAN** | Transcriptional coactivator YAP1 | HUMAN |
| **sp\|Q9Y5V3\|MAGD1_HUMAN** | Melanoma-associated antigen D1 | HUMAN |
| **sp\|P53680\|AP2S1_HUMAN** | AP-2 complex subunit sigma | HUMAN |
| **sp\|Q86TG7\|PEG10_HUMAN** | Retrotransposon-derived protein PEG10 | HUMAN |
| **sp\|P08247\|SYPH_HUMAN** | Synaptophysin | HUMAN |
| **sp\|Q16186\|ADRM1_HUMAN** | Proteasomal ubiquitin receptor ADRM1 | HUMAN |
| **sp\|P05204\|HMGN2_HUMAN** | Non-histone chromosomal protein HMG-17 | HUMAN |
| **sp\|O14770\|MEIS2_HUMAN** | Homeobox protein Meis2 | HUMAN |
| **sp\|Q8WUD4\|CCD12_HUMAN** | Coiled-coil domain-containing protein 12 | HUMAN |
| **sp\|Q15404\|RSU1_HUMAN** | Ras suppressor protein 1 | HUMAN |
| **sp\|Q8IV08\|PLD3_HUMAN** | 5'-3' exonuclease PLD3 | HUMAN |
| **sp\|P78318\|IGBP1_HUMAN** | Immunoglobulin-binding protein 1 | HUMAN |

| | | |
|---|---|---|
| **sp\|P52943\|CRIP2_HUMAN** | Cysteine-rich protein 2 | HUMAN |
| **sp\|Q96D71\|REPS1_HUMAN** | RalBP1-associated Eps domain-containing protein 1 | HUMAN |
| **sp\|Q15286\|RAB35_HUMAN** | Ras-related protein Rab-35 | HUMAN |
| **sp\|P49006\|MRP_HUMAN** | MARCKS-related protein | HUMAN |
| **sp\|Q99719\|SEPT5_HUMAN** | Septin-5 | HUMAN |
| **sp\|P49720\|PSB3_HUMAN** | Proteasome subunit beta type-3 | HUMAN |
| **sp\|P62140\|PP1B_HUMAN** | Serine/threonine-protein phosphatase PP1-beta catalytic subunit | HUMAN |
| **sp\|O95721\|SNP29_HUMAN** | Synaptosomal-associated protein 29 | HUMAN |
| **sp\|Q9H944\|MED20_HUMAN** | Mediator of RNA polymerase II transcription subunit 20 | HUMAN |
| **sp\|P48739\|PIPNB_HUMAN** | Phosphatidylinositol transfer protein beta isoform | HUMAN |
| **sp\|P07305\|H10_HUMAN** | Histone H1.0 | HUMAN |
| **sp\|P61923\|COPZ1_HUMAN** | Coatomer subunit zeta-1 | HUMAN |
| **sp\|Q9H2P0\|ADNP_HUMAN** | Activity-dependent neuroprotector homeobox protein | HUMAN |
| **sp\|Q9H7C9\|AAMDC_HUMAN** | Mth938 domain-containing protein | HUMAN |
| **sp\|Q99496\|RING2_HUMAN** | E3 ubiquitin-protein ligase RING2 | HUMAN |
| **sp\|P45973\|CBX5_HUMAN** | Chromobox protein homolog 5 | HUMAN |
| **sp\|Q92733\|PRCC_HUMAN** | Proline-rich protein PRCC | HUMAN |
| **sp\|P27797\|CALR_HUMAN** | Calreticulin | HUMAN |
| **sp\|Q9NRN7\|ADPPT_HUMAN** | L-aminoadipate-semialdehyde dehydrogenase-phosphopantetheinyl transferase | HUMAN |
| **sp\|P05387\|RLA2_HUMAN** | 60S acidic ribosomal protein P2 | HUMAN |
| **sp\|O95218\|ZRAB2_HUMAN** | Zinc finger Ran-binding domain-containing protein 2 | HUMAN |
| **sp\|Q92747\|ARC1A_HUMAN** | Actin-related protein 2/3 complex subunit 1A | HUMAN |
| **sp\|Q13247\|SRSF6_HUMAN** | Serine/arginine-rich splicing factor 6 | HUMAN |
| **sp\|Q96PU5\|NED4L_HUMAN** | E3 ubiquitin-protein ligase NEDD4-like | HUMAN |
| **sp\|Q69YQ0\|CYTSA_HUMAN** | Cytospin-A | HUMAN |
| **sp\|P62993\|GRB2_HUMAN** | Growth factor receptor-bound protein 2 | HUMAN |
| **sp\|P83916\|CBX1_HUMAN** | Chromobox protein homolog 1 | HUMAN |
| **sp\|P13591\|NCAM1_HUMAN** | Neural cell adhesion molecule 1 | HUMAN |
| **sp\|Q15813\|TBCE_HUMAN** | Tubulin-specific chaperone E | HUMAN |
| **sp\|Q9HBM1\|SPC25_HUMAN** | Kinetochore protein Spc25 | HUMAN |
| **sp\|Q08209\|PP2BA_HUMAN** | Protein phosphatase 3 catalytic subunit alpha | HUMAN |
| **sp\|P61158\|ARP3_HUMAN** | Actin-related protein 3 | HUMAN |
| **sp\|Q9UNE7\|CHIP_HUMAN** | E3 ubiquitin-protein ligase CHIP | HUMAN |
| **sp\|P35268\|RL22_HUMAN** | 60S ribosomal protein L22 | HUMAN |
| **sp\|P49902\|5NTC_HUMAN** | Cytosolic purine 5'-nucleotidase | HUMAN |
| **sp\|Q9H492\|MLP3A_HUMAN** | Microtubule-associated proteins 1A/1B light chain 3A | HUMAN |
| **sp\|P26378\|ELAV4_HUMAN** | ELAV-like protein 4 | HUMAN |

| | | |
|---|---|---|
| **sp\|P49750\|YLPM1_HUMAN** | YLP motif-containing protein 1 | HUMAN |
| **sp\|P52209\|6PGD_HUMAN** | 6-phosphogluconate dehydrogenase, decarboxylating | HUMAN |
| **sp\|Q96JE9\|MAP6_HUMAN** | Microtubule-associated protein 6 | HUMAN |
| **sp\|P42766\|RL35_HUMAN** | 60S ribosomal protein L35 | HUMAN |
| **sp\|P98179\|RBM3_HUMAN** | RNA-binding protein 3 | HUMAN |
| **sp\|Q8N9N7\|LRC57_HUMAN** | Leucine-rich repeat-containing protein 57 | HUMAN |
| **sp\|P27824\|CALX_HUMAN** | Calnexin | HUMAN |
| **sp\|O14579\|COPE_HUMAN** | Coatomer subunit epsilon | HUMAN |
| **sp\|Q99614\|TTC1_HUMAN** | Tetratricopeptide repeat protein 1 | HUMAN |
| **sp\|Q9NP72\|RAB18_HUMAN** | Ras-related protein Rab-18 | HUMAN |
| **sp\|Q9Y6A4\|CFA20_HUMAN** | Cilia- and flagella-associated protein 20 | HUMAN |
| **sp\|Q9Y678\|COPG1_HUMAN** | Coatomer subunit gamma-1 | HUMAN |
| **sp\|Q8N5M4\|TTC9C_HUMAN** | Tetratricopeptide repeat protein 9C | HUMAN |
| **sp\|Q05639\|EF1A2_HUMAN** | Elongation factor 1-alpha 2 | HUMAN |
| **sp\|Q7L5D6\|GET4_HUMAN** | Golgi to ER traffic protein 4 homolog | HUMAN |
| **sp\|Q9BYV8\|CEP41_HUMAN** | Centrosomal protein of 41 kDa | HUMAN |
| **sp\|Q14653\|IRF3_HUMAN** | Interferon regulatory factor 3 | HUMAN |
| **sp\|Q9BW83\|IFT27_HUMAN** | Intraflagellar transport protein 27 homolog | HUMAN |
| **sp\|P53582\|MAP11_HUMAN** | Methionine aminopeptidase 1 | HUMAN |
| **sp\|P62166\|NCS1_HUMAN** | Neuronal calcium sensor 1 | HUMAN |
| **sp\|P31350\|RIR2_HUMAN** | Ribonucleoside-diphosphate reductase subunit M2 | HUMAN |
| **sp\|P24666\|PPAC_HUMAN** | Low molecular weight phosphotyrosine protein phosphatase | HUMAN |
| **sp\|Q9Y6B6\|SAR1B_HUMAN** | GTP-binding protein SAR1b | HUMAN |
| **sp\|Q92688\|AN32B_HUMAN** | Acidic leucine-rich nuclear phosphoprotein 32 family member B | HUMAN |
| **sp\|P42677\|RS27_HUMAN** | 40S ribosomal protein S27 | HUMAN |
| **sp\|Q9UNT1\|RBL2B_HUMAN** | Rab-like protein 2B | HUMAN |
| **sp\|Q9Y3D6\|FIS1_HUMAN** | Mitochondrial fission 1 protein | HUMAN |
| **sp\|Q9Y296\|TPPC4_HUMAN** | Trafficking protein particle complex subunit 4 | HUMAN |
| **sp\|P41567\|EIF1_HUMAN** | Eukaryotic translation initiation factor 1 | HUMAN |
| **sp\|P30040\|ERP29_HUMAN** | Endoplasmic reticulum resident protein 29 | HUMAN |
| **sp\|P62304\|RUXE_HUMAN** | Small nuclear ribonucleoprotein E | HUMAN |
| **sp\|P21291\|CSRP1_HUMAN** | Cysteine and glycine-rich protein 1 | HUMAN |
| **sp\|P14174\|MIF_HUMAN** | Macrophage migration inhibitory factor | HUMAN |
| **sp\|P05386\|RLA1_HUMAN** | 60S acidic ribosomal protein P1 | HUMAN |
| **sp\|O94811\|TPPP_HUMAN** | Tubulin polymerization-promoting protein | HUMAN |
| **sp\|O43768\|ENSA_HUMAN** | Alpha-endosulfine | HUMAN |
| **sp\|Q8IUR0\|TPPC5_HUMAN** | Trafficking protein particle complex subunit 5 | HUMAN |
| **sp\|Q9NWB6\|ARGL1_HUMAN** | Arginine and glutamate-rich protein 1 | HUMAN |
| **sp\|P63272\|SPT4H_HUMAN** | Transcription elongation factor SPT4 | HUMAN |
| **sp\|P61960\|UFM1_HUMAN** | Ubiquitin-fold modifier 1 | HUMAN |

| | | |
|---|---|---|
| **sp\|Q7LBR1\|CHM1B_HUMAN** | Charged multivesicular body protein 1b | HUMAN |
| **sp\|Q9H0U4\|RAB1B_HUMAN** | Ras-related protein Rab-1B | HUMAN |
| **sp\|O75351\|VPS4B_HUMAN** | Vacuolar protein sorting-associated protein 4B | HUMAN |
| **sp\|Q9BX40\|LS14B_HUMAN** | Protein LSM14 homolog B | HUMAN |
| **sp\|Q92572\|AP3S1_HUMAN** | AP-3 complex subunit sigma-1 | HUMAN |
| **sp\|Q9NYB9\|ABI2_HUMAN** | Abl interactor 2 | HUMAN |
| **sp\|P21579\|SYT1_HUMAN** | Synaptotagmin-1 | HUMAN |
| **sp\|O15144\|ARPC2_HUMAN** | Actin-related protein 2/3 complex subunit 2 | HUMAN |
| **sp\|Q6IBS0\|TWF2_HUMAN** | Twinfilin-2 | HUMAN |
| **sp\|P04632\|CPNS1_HUMAN** | Calpain small subunit 1 | HUMAN |
| **sp\|Q6VY07\|PACS1_HUMAN** | Phosphofurin acidic cluster sorting protein 1 | HUMAN |
| **sp\|P47914\|RL29_HUMAN** | 60S ribosomal protein L29 | HUMAN |
| **sp\|Q04724\|TLE1_HUMAN** | Transducin-like enhancer protein 1 | HUMAN |
| **sp\|Q04727\|TLE4_HUMAN** | Transducin-like enhancer protein 4 | HUMAN |
| **sp\|Q07812\|BAX_HUMAN** | Apoptosis regulator BAX | HUMAN |
| **sp\|Q13098\|CSN1_HUMAN** | COP9 signalosome complex subunit 1 | HUMAN |
| **sp\|Q15819\|UB2V2_HUMAN** | Ubiquitin-conjugating enzyme E2 variant 2 | HUMAN |
| **sp\|Q9BXJ9\|NAA15_HUMAN** | N-alpha-acetyltransferase 15, NatA auxiliary subunit | HUMAN |
| **sp\|Q9BXS5\|AP1M1_HUMAN** | AP-1 complex subunit mu-1 | HUMAN |
| **sp\|Q9P0V9\|SEP10_HUMAN** | Septin-10 | HUMAN |
| **sp\|Q53T59\|H1BP3_HUMAN** | HCLS1-binding protein 3 | HUMAN |
| **sp\|Q8IUE6\|H2A2B_HUMAN** | Histone H2A type 2-B | HUMAN |
| **sp\|Q9GZT4\|SRR_HUMAN** | Serine racemase | HUMAN |
| **sp\|Q9BZL1\|UBL5_HUMAN** | Ubiquitin-like protein 5 | HUMAN |
| **sp\|Q3MHD2\|LSM12_HUMAN** | Protein LSM12 | HUMAN |
| **sp\|P52298\|NCBP2_HUMAN** | Nuclear cap-binding protein subunit 2 | HUMAN |
| **sp\|P19367\|HXK1_HUMAN** | Hexokinase-1 | HUMAN |
| **sp\|Q05193\|DYN1_HUMAN** | Dynamin-1 | HUMAN |
| **sp\|Q15382\|RHEB_HUMAN** | GTP-binding protein Rheb | HUMAN |
| **sp\|P60520\|GBRL2_HUMAN** | Gamma-aminobutyric acid receptor-associated protein-like 2 | HUMAN |
| **sp\|Q86WG3\|ATCAY_HUMAN** | Caytaxin | HUMAN |
| **sp\|Q15287\|RNPS1_HUMAN** | RNA-binding protein with serine-rich domain 1 | HUMAN |
| **sp\|O95104\|SCAF4_HUMAN** | SR-related and CTD-associated factor 4 | HUMAN |
| **sp\|Q9UK41\|VPS28_HUMAN** | Vacuolar protein sorting-associated protein 28 homolog | HUMAN |
| **sp\|Q9Y530\|OARD1_HUMAN** | ADP-ribose glycohydrolase OARD1 | HUMAN |
| **sp\|P35659\|DEK_HUMAN** | Protein DEK | HUMAN |
| **sp\|Q96T60\|PNKP_HUMAN** | Bifunctional polynucleotide phosphatase/kinase | HUMAN |
| **sp\|Q8WUX9\|CHMP7_HUMAN** | Charged multivesicular body protein 7 | HUMAN |
| **sp\|P30044\|PRDX5_HUMAN** | Peroxiredoxin-5, mitochondrial | HUMAN |
| **sp\|Q9BTE1\|DCTN5_HUMAN** | Dynactin subunit 5 | HUMAN |

| | | |
|---|---|---|
| **sp\|Q8IYB5\|SMAP1_HUMAN** | Stromal membrane-associated protein 1 | HUMAN |
| **sp\|P62875\|RPAB5_HUMAN** | DNA-directed RNA polymerases I, II, and III subunit RPABC5 | HUMAN |
| **sp\|Q9NVM6\|DJC17_HUMAN** | DnaJ homolog subfamily C member 17 | HUMAN |
| **sp\|Q9BY43\|CHM4A_HUMAN** | Charged multivesicular body protein 4a | HUMAN |
| **sp\|Q9NQ48\|LZTL1_HUMAN** | Leucine zipper transcription factor-like protein 1 | HUMAN |
| **sp\|Q9H814\|PHAX_HUMAN** | Phosphorylated adapter RNA export protein | HUMAN |
| **sp\|Q7Z5L9\|I2BP2_HUMAN** | Interferon regulatory factor 2-binding protein 2 | HUMAN |
| **sp\|P62834\|RAP1A_HUMAN** | Ras-related protein Rap-1A | HUMAN |
| **sp\|Q8N1B4\|VPS52_HUMAN** | Vacuolar protein sorting-associated protein 52 homolog | HUMAN |
| **sp\|P53041\|PPP5_HUMAN** | Serine/threonine-protein phosphatase 5 | HUMAN |
| **sp\|Q07157\|ZO1_HUMAN** | Tight junction protein ZO-1 | HUMAN |
| **sp\|P46778\|RL21_HUMAN** | 60S ribosomal protein L21 | HUMAN |
| **sp\|O75150\|BRE1B_HUMAN** | E3 ubiquitin-protein ligase BRE1B | HUMAN |
| **sp\|Q12979\|ABR_HUMAN** | Active breakpoint cluster region-related protein | HUMAN |
| **sp\|Q9NSC5\|HOME3_HUMAN** | Homer protein homolog 3 | HUMAN |
| **sp\|P41208\|CETN2_HUMAN** | Centrin-2 | HUMAN |
| **sp\|Q92609\|TBCD5_HUMAN** | TBC1 domain family member 5 | HUMAN |
| **sp\|O95793\|STAU1_HUMAN** | Double-stranded RNA-binding protein Staufen homolog 1 | HUMAN |
| **sp\|Q16775\|GLO2_HUMAN** | Hydroxyacylglutathione hydrolase, mitochondrial | HUMAN |
| **sp\|Q9NPA8\|ENY2_HUMAN** | Transcription and mRNA export factor ENY2 | HUMAN |
| **sp\|Q08117\|TLE5_HUMAN** | TLE family member 5 | HUMAN |
| **sp\|P17600\|SYN1_HUMAN** | Synapsin-1 | HUMAN |
| **sp\|P48059\|LIMS1_HUMAN** | LIM and senescent cell antigen-like-containing domain protein 1 | HUMAN |
| **sp\|Q9HB71\|CYBP_HUMAN** | Calcyclin-binding protein | HUMAN |
| **sp\|P20962\|PTMS_HUMAN** | Parathymosin | HUMAN |
| **sp\|P61006\|RAB8A_HUMAN** | Ras-related protein Rab-8A | HUMAN |
| **sp\|P45985\|MP2K4_HUMAN** | Dual specificity mitogen-activated protein kinase kinase 4 | HUMAN |
| **sp\|Q9UJY5\|GGA1_HUMAN** | ADP-ribosylation factor-binding protein GGA1 | HUMAN |
| **sp\|O15145\|ARPC3_HUMAN** | Actin-related protein 2/3 complex subunit 3 | HUMAN |
| **sp\|O15511\|ARPC5_HUMAN** | Actin-related protein 2/3 complex subunit 5 | HUMAN |
| **sp\|Q9UMZ2\|SYNRG_HUMAN** | Synergin gamma | HUMAN |
| **sp\|Q66K74\|MAP1S_HUMAN** | Microtubule-associated protein 1S | HUMAN |
| **sp\|P12268\|IMDH2_HUMAN** | Inosine-5'-monophosphate dehydrogenase 2 | HUMAN |
| **sp\|Q9H0Q0\|CYRIA_HUMAN** | CYFIP-related Rac1 interactor A | HUMAN |
| **sp\|Q9NPD3\|EXOS4_HUMAN** | Exosome complex component RRP41 | HUMAN |
| **sp\|Q15836\|VAMP3_HUMAN** | Vesicle-associated membrane protein 3 | HUMAN |
| **sp\|Q9P035\|HACD3_HUMAN** | Very-long-chain (3R)-3-hydroxyacyl-CoA dehydratase 3 | HUMAN |
| **sp\|Q9UKZ1\|CNO11_HUMAN** | CCR4-NOT transcription complex subunit 11 | HUMAN |

| | | |
|---|---|---|
| **sp\|Q8NCA5\|FA98A_HUMAN** | Protein FAM98A | HUMAN |
| **sp\|Q06210\|GFPT1_HUMAN** | Glutamine--fructose-6-phosphate aminotransferase [isomerizing] 1 | HUMAN |
| **sp\|P06753\|TPM3_HUMAN** | Tropomyosin alpha-3 chain | HUMAN |
| **sp\|Q93045\|STMN2_HUMAN** | Stathmin-2 | HUMAN |
| **sp\|Q32MZ4\|LRRF1_HUMAN** | Leucine-rich repeat flightless-interacting protein 1 | HUMAN |
| **sp\|Q13185\|CBX3_HUMAN** | Chromobox protein homolog 3 | HUMAN |
| **sp\|Q9H3H9\|TCAL2_HUMAN** | Transcription elongation factor A protein-like 2 | HUMAN |
| **sp\|Q9BW30\|TPPP3_HUMAN** | Tubulin polymerization-promoting protein family member 3 | HUMAN |
| **sp\|Q8NB37\|GALD1_HUMAN** | Glutamine amidotransferase-like class 1 domain-containing protein 1 | HUMAN |
| **sp\|Q00765\|REEP5_HUMAN** | Receptor expression-enhancing protein 5 | HUMAN |
| **sp\|P68371\|TBB4B_HUMAN** | Tubulin beta-4B chain | HUMAN |
| **sp\|Q13885\|TBB2A_HUMAN** | Tubulin beta-2A chain | HUMAN |
| **sp\|P62491\|RB11A_HUMAN** | Ras-related protein Rab-11A | HUMAN |
| **sp\|Q14240\|IF4A2_HUMAN** | Eukaryotic initiation factor 4A-II | HUMAN |
| **sp\|Q9NRW1\|RAB6B_HUMAN** | Ras-related protein Rab-6B | HUMAN |
| **sp\|P22694\|KAPCB_HUMAN** | cAMP-dependent protein kinase catalytic subunit beta | HUMAN |
| **sp\|Q01085\|TIAR_HUMAN** | Nucleolysin TIAR | HUMAN |
| **sp\|P63167\|DYL1_HUMAN** | Dynein light chain 1, cytoplasmic | HUMAN |
| **sp\|P47755\|CAZA2_HUMAN** | F-actin-capping protein subunit alpha-2 | HUMAN |
| **sp\|O95197\|RTN3_HUMAN** | Reticulon-3 | HUMAN |
| **sp\|Q96LR5\|UB2E2_HUMAN** | Ubiquitin-conjugating enzyme E2 E2 | HUMAN |
| **sp\|P40123\|CAP2_HUMAN** | Adenylyl cyclase-associated protein 2 | HUMAN |
| **sp\|Q969Q0\|RL36L_HUMAN** | 60S ribosomal protein L36a-like | HUMAN |
| **sp\|P62861\|RS30_HUMAN** | FAU ubiquitin-like and ribosomal protein S30 | HUMAN |
| **sp\|P09132\|SRP19_HUMAN** | Signal recognition particle 19 kDa protein | HUMAN |
| **sp\|Q9Y2V2\|CHSP1_HUMAN** | Calcium-regulated heat-stable protein 1 | HUMAN |
| **sp\|Q9Y2S6\|TMA7_HUMAN** | Translation machinery-associated protein 7 | HUMAN |
| **sp\|Q9UHR5\|S30BP_HUMAN** | SAP30-binding protein | HUMAN |
| **sp\|Q9H832\|UBE2Z_HUMAN** | Ubiquitin-conjugating enzyme E2 Z | HUMAN |
| **sp\|Q9BSV6\|SEN34_HUMAN** | tRNA-splicing endonuclease subunit Sen34 | HUMAN |
| **sp\|Q13510\|ASAH1_HUMAN** | Acid ceramidase | HUMAN |
| **sp\|P80723\|BASP1_HUMAN** | Brain acid soluble protein 1 | HUMAN |
| **sp\|P78356\|PI42B_HUMAN** | Phosphatidylinositol 5-phosphate 4-kinase type-2 beta | HUMAN |
| **sp\|O75396\|SC22B_HUMAN** | Vesicle-trafficking protein SEC22b | HUMAN |
| **sp\|Q96P47\|AGAP3_HUMAN** | Arf-GAP with GTPase, ANK repeat and PH domain-containing protein 3 | HUMAN |
| **sp\|P22061\|PIMT_HUMAN** | Protein-L-isoaspartate(D-aspartate) O-methyltransferase | HUMAN |

| | | |
|---|---|---|
| **sp\|Q6QNY0\|BL1S3_HUMAN** | Biogenesis of lysosome-related organelles complex 1 subunit 3 | HUMAN |
| **sp\|Q9NY12\|GAR1_HUMAN** | H/ACA ribonucleoprotein complex subunit 1 | HUMAN |
| **sp\|P35813\|PPM1A_HUMAN** | Protein phosphatase 1A | HUMAN |
| **sp\|Q8NFW8\|NEUA_HUMAN** | N-acylneuraminate cytidylyltransferase | HUMAN |
| **sp\|Q9Y312\|AAR2_HUMAN** | Protein AAR2 homolog | HUMAN |
| **sp\|Q8ND56\|LS14A_HUMAN** | Protein LSM14 homolog A | HUMAN |
| **sp\|Q9Y4C2\|TCAF1_HUMAN** | TRPM8 channel-associated factor 1 | HUMAN |
| **sp\|Q8NAF0\|ZN579_HUMAN** | Zinc finger protein 579 | HUMAN |
| **sp\|P62910\|RL32_HUMAN** | 60S ribosomal protein L32 | HUMAN |
| **sp\|Q6VEQ5\|WASH2_HUMAN** | WAS protein family homolog 2 | HUMAN |
| **sp\|P07355\|ANXA2_HUMAN** | Annexin A2 | HUMAN |
| **sp\|P60709\|ACTB_HUMAN** | Actin, cytoplasmic 1 | HUMAN |
| **sp\|O14910\|LIN7A_HUMAN** | Protein lin-7 homolog A | HUMAN |
| **sp\|Q96I24\|FUBP3_HUMAN** | Far upstream element-binding protein 3 | HUMAN |
| **sp\|P11717\|MPRI_HUMAN** | Cation-independent mannose-6-phosphate receptor | HUMAN |
| **sp\|Q14919\|NC2A_HUMAN** | Dr1-associated corepressor | HUMAN |
| **sp\|Q9Y490\|TLN1_HUMAN** | Talin-1 | HUMAN |
| **sp\|P06746\|DPOLB_HUMAN** | DNA polymerase beta | HUMAN |
| **sp\|P84022\|SMAD3_HUMAN** | Mothers against decapentaplegic homolog 3 | HUMAN |
| **sp\|P37802\|TAGL2_HUMAN** | Transgelin-2 | HUMAN |
| **sp\|Q16352\|AINX_HUMAN** | Alpha-internexin | HUMAN |
| **sp\|Q6UUV9\|CRTC1_HUMAN** | CREB-regulated transcription coactivator 1 | HUMAN |
| **sp\|O14602\|IF1AY_HUMAN** | Eukaryotic translation initiation factor 1A, Y-chromosomal | HUMAN |
| **sp\|P37235\|HPCL1_HUMAN** | Hippocalcin-like protein 1 | HUMAN |
| **sp\|Q9BRJ7\|TIRR_HUMAN** | Tudor-interacting repair regulator protein | HUMAN |
| **sp\|O95292\|VAPB_HUMAN** | Vesicle-associated membrane protein-associated protein B/C | HUMAN |
| **sp\|O75525\|KHDR3_HUMAN** | KH domain-containing, RNA-binding, signal transduction-associated protein 3 | HUMAN |
| **sp\|Q8WUH6\|TM263_HUMAN** | Transmembrane protein 263 | HUMAN |
| **sp\|Q9H6Z4\|RANB3_HUMAN** | Ran-binding protein 3 | HUMAN |

**Supp. Table S3.**

**Supplementary Table S3: List of 40 kinases bound by PS-NPs, detected by LC-MS/MS**

| Accession | Protein Name | Unique to PS-NPs (Not bound by silica) |
|---|---|---|
| sp\|P78527\|PRKDC_HUMAN | DNA-dependent protein kinase catalytic subunit | ✗ |
| sp\|P14618\|KPYM_HUMAN | Pyruvate kinase PKM | ✗ |
| sp\|P17858\|PFKAL_HUMAN | ATP-dependent 6-phosphofructokinase, liver type | ✗ |
| sp\|Q13177\|PAK2_HUMAN | Serine/threonine-protein kinase PAK 2 | ✗ |
| sp\|O95819\|M4K4_HUMAN | Mitogen-activated protein kinase kinase kinase kinase 4 | ✓ |
| sp\|P00558\|PGK1_HUMAN | Phosphoglycerate kinase 1 | ✗ |
| sp\|P28482\|MK01_HUMAN | Mitogen-activated protein kinase 1 | ✓ |
| sp\|P12277\|KCRB_HUMAN | Creatine kinase B-type | ✗ |
| sp\|P36507\|MP2K2_HUMAN | Dual specificity mitogen-activated protein kinase kinase 2 | ✓ |
| sp\|P06493\|CDK1_HUMAN | Cyclin-dependent kinase 1 | ✗ |
| sp\|O15075\|DCLK1_HUMAN | Serine/threonine-protein kinase DCLK1 | ✗ |
| sp\|O95747\|OXSR1_HUMAN | Serine/threonine-protein kinase OSR1 | ✗ |
| sp\|P68400\|CSK21_HUMAN | Casein kinase II subunit alpha | ✓ |
| sp\|Q13557\|KCC2D_HUMAN | Calcium/calmodulin-dependent protein kinase type II subunit delta | ✓ |
| sp\|P13861\|KAP2_HUMAN | cAMP-dependent protein kinase type II-alpha regulatory subunit | ✓ |
| sp\|Q01813\|PFKAP_HUMAN | ATP-dependent 6-phosphofructokinase, platelet type | ✓ |
| sp\|P17612\|KAPCA_HUMAN | cAMP-dependent protein kinase catalytic subunit alpha | ✓ |
| sp\|P08237\|PFKAM_HUMAN | ATP-dependent 6-phosphofructokinase, muscle type | ✓ |
| sp\|P45983\|MK08_HUMAN | Mitogen-activated protein kinase 8 | ✓ |
| sp\|P54619\|AAKG1_HUMAN | 5'-AMP-activated protein kinase subunit gamma-1 | ✗ |
| sp\|P15531\|NDKA_HUMAN | Nucleoside diphosphate kinase A | ✗ |
| sp\|P51812\|KS6A3_HUMAN | Ribosomal protein S6 kinase alpha-3 | ✓ |
| sp\|P27361\|MK03_HUMAN | Mitogen-activated protein kinase 3 | ✓ |
| sp\|P67870\|CSK2B_HUMAN | Casein kinase II subunit beta | ✗ |
| sp\|Q9H2G2\|SLK_HUMAN | STE20-like serine/threonine-protein kinase | ✓ |
| sp\|Q02750\|MP2K1_HUMAN | Dual specificity mitogen-activated protein kinase kinase 1 | ✓ |
| sp\|Q13418\|ILK_HUMAN | Integrin-linked protein kinase | ✓ |
| sp\|Q9Y3S1\|WNK2_HUMAN | Serine/threonine-protein kinase WNK2 | ✓ |

| | | |
|---|---|---|
| sp\|Q9P0L2\|MARK1_HUMAN | Serine/threonine-protein kinase MARK1 | ✓ |
| sp\|Q96KB5\|TOPK_HUMAN | Lymphokine-activated killer T-cell-originated protein kinase | ✓ |
| sp\|Q00535\|CDK5_HUMAN | Cyclin-dependent kinase 5 | ✓ |
| sp\|P49841\|GSK3B_HUMAN | Glycogen synthase kinase-3 beta | ✓ |
| sp\|P48729\|KC1A_HUMAN | Casein kinase I isoform alpha | ✓ |
| sp\|P19784\|CSK22_HUMAN | Casein kinase II subunit alpha' | ✓ |
| sp\|Q03426\|KIME_HUMAN | Mevalonate kinase | ✓ |
| sp\|P19367\|HXK1_HUMAN | Hexokinase-1 | ✓ |
| sp\|Q96T60\|PNKP_HUMAN | Bifunctional polynucleotide phosphatase/kinase | ✓ |
| sp\|P45985\|MP2K4_HUMAN | Dual specificity mitogen-activated protein kinase kinase 4 | ✓ |
| sp\|P22694\|KAPCB_HUMAN | cAMP-dependent protein kinase catalytic subunit beta | ✓ |
| sp\|P78356\|PI42B_HUMAN | Phosphatidylinositol 5-phosphate 4-kinase type-2 beta | ✓ |

**Colour legend for Table S3:**

Kinases known to phosphorylate TDP43 are shown in red.
Kinases known to phosphorylate Tau are shown in blue.
Kinases known to phosphorylate both TDP43 and Tau are shown in green.
